# Gut Microbiome-derived Propionate Reprograms Alveolar Macrophages Metabolically and Regulates Lung Injury Responses

**DOI:** 10.1101/2025.04.14.648797

**Authors:** Daisuke Maruyama, Xiaoli Tian, Thien N. M. Doan, Wen-I Liao, Tomohiro Chaki, Hiroki Taenaka, Mazharul Maishan, Brian Layden, Michael A. Matthay, Arun Prakash

**Author notes:** Corresponding Author: Arun Prakash, M.D., Ph.D., Department of Anesthesia and Perioperative Care; San Francisco General Hospital, University of California, San Francisco. Department of Emergency Medicine, Tri-Service General Hospital, National Defense Medical Center, Taipei, Taiwan (Wen-I Liao). Home institution: Department of Anesthesiology, Sapporo Medical University School of Medicine, Sapporo, Japan (Tomohiro Chaki). Department of Medicine, University of Illinois Chicago (Brian Layden). Department of Medicine and Anesthesia, University of California San Francisco (Hiroki Taenaka, Mazharul Maishan, Michael A. Matthay). **Department/Institution to which the work is attributed:** Department of Anesthesia and Perioperative Care, University of California San Francisco and San Francisco General Hospital, San Francisco, California. **Meetings at which portions of this work have been presented:** American Society of Anesthesiologists (ASA) Annual meeting, October 2021, San Diego; American Thoracic Society (ATS) Annual meeting, May 2023, Washington, DC; Cell Symposia: Infection biology in the age of the microbiome, June 2023; Inserm Workshop 272: Les microbiotes – de l’exploration à l’application thérapeutique, May 2023; Lung Development, Injury and Repair GRC, August 2023. **Contributions of Authors:** AP designed the study, prepared figures, wrote manuscript and is the corresponding author. DM, TNMD, WIL, TC and XT designed and carried out experiments, contributed to manuscript writing. MB performed bioinformatics analyses and prepared figures. MM and TH performed pneumonia infection experiments and were overseen by MAM. BL provided FFAR2KO and FFAR3KO mice.

## Abstract

Responses to lung injury can vary between individuals and the diet and gut microbiome represent two underappreciated sources of inter-individual variability. The gut microbiome can influence lung injury outcomes through the gut-lung axis, but exactly how diet and its effects on the microbiota are involved remains unclear. We hypothesized that a dietary fiber intervention would enrich for the presence of short-chain fatty acid (SCFA)-producing fermentative bacteria in the gut microbiome, thereby influencing resting lung immunometabolic tone and influence downstream immune responses to lung injury and infection. To test this hypothesis, we fed mice fiber-rich (FR) and fiber-free (FF) diets and observed changes in the steady-state transcriptional programming of alveolar macrophages (AMs) in the absence of injury. Next, we examined the effects of FR and FF diets on the responses to sterile and infectious lung injury *in vivo* while simultaneously profiling gut microbiota and SCFA levels transmitted along the gut-lung axis. Finally, we validated our *in vivo* observations with mechanistic studies of the metabolic, signaling, and chromatin-modifying effects of specific SCFAs on lung AMs *ex vivo* and *in vitro*. Overall, a FR diet reprogrammed AMs and attenuated lung inflammation after sterile injury while exacerbating lung infection. This effect of FR diets could be transferred to germ-free (GF) mice by fecal microbiome transplantation (FMT) and depended on the ability of the microbiota to produce propionate. Mechanistically, SCFAs altered the metabolic programming of AMs and lung tissue *ex vivo* without a role for free fatty acid receptors (FFAR) or chromatin remodeling. These findings demonstrate that the gut-lung axis can regulate the resting metabolic tone through dietary fiber intake and the enrichment of SCFA-producing gut bacteria, as well as influence sterile and non-sterile lung injury responses through the activity of gut metabolites. These results provide further evidence to support the development of therapeutic dietary interventions or the use of specific gut metabolites to preserve or enhance specific aspects of host pulmonary immunity.

## INTRODUCTION

The influence of the gut microbiome on human health and disease has been closely studied over the past decade^1–4^ with great progress made, but how gut microbiota control lung immune injury responses via the gut-lung axis is not well understood^2,5,6^. High-fiber diets have been shown to promote the expansion of fermenting gut bacteria resulting in the production of important metabolites, including short-chain fatty acids (SCFA) – those with acyl chains up to 6 carbons in length. SCFA metabolites are now well recognized as immune and metabolic regulators of gut health and permeability and have also been implicated in modulating health and disease states throughout the body^1,6–10^. However, the key mechanisms for how these changes in diet and gut microbiome tune lung metabolic and immune states remain undefined as do the specific effects of key bacterial species and metabolites. The emerging intersection and interactions between metabolism and immunity has been termed immunometabolism and can be studied both at the cellular and organismal levels, with key roles for the diet and the gut microbiome (reviewed in ^11–17)^. Therefore, the identification of dietary factors, specific gut bacteria and bioactive metabolites that influence lung immune and injury responses could lead to a better understanding of inter-individual immune variability and could be exploited for personalized therapeutic approaches and applications.

The immunometabolic tone of the lung at a tissue level refers to the level of immune priming and the baseline metabolic set-point present in the lung, which in turn may dictate the degree of inflammatory response following injury. Two key inflammasome-dependent cytokines, IL-1β and IL-18 may contribute to this tone and have been implicated in a variety of lung pathologies, including sterile injury ^18–21^, infection^22,23^, and asthma ^24–26^. We previously reported that SCFA metabolites derived from the gut microbiome can translocate from the gut to the lung in mice and can regulate the IL-1β-dependent inflammatory output of alveolar macrophages (AMs) *in vitro* ^21,27^.

SCFA (acetate, propionate, and butyrate) are primarily produced by fiber fermentation by gut bacteria and are thought to exert their biological effects on the host through three primary mechanisms (reviewed in ^1,7,28,29^). First, they can serve as a local energy source for colonic epithelia and potentially other cell types. Second, propionate and butyrate can act as histone deacetylase inhibitors (HDACi) targeting specific gene loci. Finally, all three SCFAs can signal via free fatty acid receptors (FFARs – primarily FFAR2 and FFAR3), which are G-protein coupled receptors that can regulate gene transcription in the affected cell and influence inflammatory output.

In this study, we used pectin-based fiber-rich (FR) and fiber-free (FF) diets to investigate the effects of fiber on lung immunometabolic tone at steady-state, specifically on alveolar macrophages (AMs). We observed that dietary fiber could also regulate lung inflammatory responses after sterile ischemia reperfusion (IR) injury, and correlated changes in lung inflammatory markers with specific gut bacterial taxa and SCFAs. We then investigated whether the diet-induced lung IR injury responses could be transferred to germ-free (GF) mice via fecal microbiome transplantation (FMT) and how dietary fiber might affect the course of a bacterial pneumonia in mice. Finally, using *in vivo* as well as *in vitro* and *ex vivo* cellular and tissue approaches, we mechanistically studied how propionate was able to alter the inflammatory responses and metabolic activity of AMs and whether these effects depended on FFAR signaling and/or changes in chromatin accessibility.

## MATERIALS AND METHODS

### Animal care

All studies were approved by the Institutional Animal Care and Use Committee at the University of California, San Francisco (UCSF) (IACUC protocol# AN197325-01). All mice were purchased from the Jackson Laboratory (Bar Harbor, ME) or bred in the animal facilities at UCSF. Wild-type (WT) C57BL/6 and germ-free (GF aka gnotobiotic) male mice aged 10-15 weeks were used in this study. Commercially purchased mice were acclimated to their new housing for at least 1 week before the initiation of any experiments procedures. FFAR2 and FFAR3KO mice were generated as reported previously 30,31 and generously provided by B. Layden (University of Illinois Chicago).

### Fiber diet interventions

Purchased mice received standard animal facility chow (normal diet, i.e. ND) for at least 1 week or more after arrival at UCSF, after which they received either 2 weeks of 0% fiber (FF), 2 weeks of 35% pectin fiber (FR 2wk or FR), 1 week of FF followed by 1 week of FR (FR 1wk or FF->FR), or continued on ND. Diets were purchased from Newco Distributors, Inc. The FF diet was TestDiet® Modified AIN-93G without cellulose (catalog number 5GCX), while the FR diet was TestDiet® Modified 57W5 with pectin (catalog number 5Z6R). The ND consisted of standard UCSF mouse chow containing 2.4% dietary fiber. Although these three diets were not isocaloric, the mice had *ad libitum* access to food. The choice of FF and FR diet allowed us to evaluate the role of fiber without introducing potential confounding effects associated with making the diets isocaloric by adding fat or other calorie sources (**Figure S1A**). Fecal quality and intestinal morphology were assessed after IR surgery (**Figure S1B**), along with monitoring weight changes throughout the dietary intervention. No significant weight loss was observed during the study period (**Figure S1C**). Furthermore, energy contributions from fiber fermentation and other bacterial breakdown were not accounted for in the dietary caloric calculations.

### Left lung ischemia reperfusion (IR) surgery

A mouse model of unilateral left pulmonary artery occlusion was used, as previously described ^30^. Briefly, mice were anesthetized with intraperitoneal (IP) 2,2,2-tribromoethanol (Avertin®; catalog number T48402, Sigma-Aldrich), orally intubated, administered IP buprenorphine (catalog number 059122, Covetrus North America), and placed on a rodent ventilator with a tidal volumes of 225 uL (7.5 cc/kg) and a respiratory rate of 180 breaths/min (assuming an average mouse weight of 30 g). A left thoracotomy was performed via the intercostal space between the 2nd and 3rd ribs. The left pulmonary artery (PA) was identified and ligated with a slipknot using 7-0 monofilament suture (catalog number 8696G, Ethicon). The end of the suture was externalized to the anterior chest wall through a narrow bore (27g) needle. Before closing the chest cavity, the left lung was reinflated with positive end pressure. Local anesthesia was provided using 3-4 drops of 0.25% bupivacaine (catalog number NDC 0409-1159-19, Hospira, Inc.) applied topically prior to skin closure. The total duration of mechanical ventilation and surgery was approximately 20-25 min. After skin closure, mice were extubated and allowed to recover from anesthesia. After 60 min of ischemia, the pulmonary artery ligature was released, and left lung reperfusion was initiated. At the experimental end point (1h post reperfusion), mice were euthanized, and the blood, feces, and lungs were collected.

Blood was collected from anesthetized mice through inferior vena cava (IVC) and portal vein puncture with a heparinized syringe, centrifuged (14,000g, 5 min) and the plasma was separated, snap frozen in liquid nitrogen and stored at −80°C. The lower portions of the left lungs were excised and divided into two parts: one part was placed in TRIzol® (catalog number 15596018, Invitrogen) for RNA preparation and the second part was frozen at −80°C for homogenization for ELISA. Cytokines and chemokine levels were quantified in plasma or homogenized lung tissue by ELISA or in lung tissue by qPCR.

All mice received equivalent durations of mechanical ventilation (∼40-45 min) and were allowed to breathe spontaneously during their recovery from anesthesia and the remainder of the ischemia and subsequent reperfusion periods. Pre- and postoperative care was performed according to ARRIVE guidelines.

The success rate for the lung IR surgery is approximately 80-90%. Mice that did not survive the IR surgery or the reperfusion period due to technical complications in the surgical procedure (predominantly, left bronchus or left PA injury) were mostly excluded from the study, except for the analysis of effects of the dietary fiber on 16S rRNA sequence microbiome composition. In the case of experiments involving limited availability of mice, namely, experiments involving the GF mice given FMTs, both mice that survived IR surgery and those that did not were included (and the injury termed “lung injury”) with the rationale that the entry into the left thorax, left lung collapse, left lung manipulation – all generated sterile lung injury and based on our prior work with human lung tissue, the affected lung tissue and cells would still be alive and undergoing inflammatory responses 2h following the start of the surgical procedure^21^. In the experiments to investigate direct effects of C3 on *in vivo* lung IR, C3 (10 mmol/kg) was administered IP 1.5h before the start of the ischemic period.

### Bulk RNAseq Analysis of Bronchoalveolar Lavage (BAL) cells

Two groups of WT C57BL/6 mice (n=10 each) were fed FR and FF diets for 2 weeks. BAL was performed as previously described ^21^ and RNA quantity and quality were measured. Briefly, mice were euthanized under anesthesia (Avertin plus buprenorphine), and their tracheas were surgically exposed.

Alveolar lavage was performed by injecting and withdrawing 10 mL of ice-cold phosphate-buffered saline (PBS; catalog number 14190144, Gibco) containing 2 mM EDTA (catalog number 45001-122, Corning), into the trachea, 1 mL at a time. Cells were pelleted at 300g x 4 min at 4°C and then resuspended in TRIzol^®^.

One hundred ng of RNA for each sample was submitted to the UCSF ImmunoX Genomics CoLabs for bulk RNAseq. RIN scores were all between 9-10. Illumina compatible RNA-sequencing libraries were generated from purified RNA using the Tecan Universal mRNA Plus kit (9156-A01) by the UCSF Genomics CoLabs. Libraries were sequenced on an Illumina NovaSeq X using paired end 50bp reads at the UCSF Center for Advanced Technology (https://cat.ucsf.edu). Sequencing reads were aligned to the mouse reference genome (GRCm38) and reads per gene matrix were counted using the Ensemble annotation build version 96 with STAR v2.7.5c^31^. Read counts per gene were used as input to DESeq2 ^32^ to test for differential gene expression between conditions using a Wald test, while correcting for possible covariates. Genes passing a multiple testing correction p-value of 0.1 (FDR method) were considered significant. Differential gene expression was performed and heat maps were generated as shown. Gene enrichment analysis was also performed using MSigDB, KEGG, and GO databases.

### Reagents and Cell Lines

Propionate (catalog number 18108), acetate (catalog number 241245), butyrate (catalog number B5887), indole-3-propionate (IPA, catalog number 57410), and lipopolysaccharide (LPS; catalog number L4391) were purchased from Sigma-Aldrich (St. Louis, MO). Seahorse XF calibrant solution (part number 100840-000), Seahorse XF Dulbecco’s Modified Eagle Medium (DMEM), pH7.4 (catalog number 103575-100), Seahorse XF RPMI medium, pH 7.4 (catalog number 103576-100), Seahorse XF 1.0 M glucose solution (catalog number 103577-100), Seahorse XF 100 mM pyruvate solution (catalog number 103578-100), Seahorse XF 200 mM glutamine solution (catalog number 103579-100), and Seahorse XF Cell Mito Stress Test Kit (catalog number 103015-100; includes oligomycin, FCCP, and rotenone/antimycin A) were all purchased from Agilent Technologies, Inc. (Santa Clara, CA).

The cell lines used in this study were MLE-12 (wild-type [WT] FVB/N murine lung epithelial alveolar type II [AT2] cells; catalog number CRL-2110) and MH-S (WT BALB/c alveolar macrophage cells; catalog number CRL-2019), both purchased from the American Type Culture Collection (ATCC; Manassas, VA).

### Collection of Murine Primary Alveolar Macrophages (AMs) by BAL

AMs were isolated via BAL from WT, FFAR2 KO and FFAR3 KO mice fed a ND as well as WT mice fed an FF or FR diet for 2 weeks. None of the mice had undergone *in vivo* lung injury. BAL was performed as described earlier, and cells from up to 4 mice were pooled, or processed individually if only one mouse was used, then pelleted at 300g x 4 min at 4°C. The pellets were resuspended in Roswell Park Memorial Institute (RPMI) 1640 medium (catalog number 72400047, Gibco) supplemented with 10% fetal bovine serum (FBS; catalog number 10438026, Life Technologies) and 1% penicillin-streptomycin (P/S; catalog number 30-002-CI, Corning) and plated in a 48-well plate at 200,000 cells per well and were incubated overnight at 37°C under humidified 5% CO_2_.

Additionally, cytospin preparations were performed on a portion of the cell suspensions to verify the morphology of mouse AMs. A 100 µL aliquot of cell suspension was centrifuged onto a glass slide at 500 rpm for 5 min using a Cytospin 4 centrifuge (Thermo Fisher). The slides were air-dried and stained with Kwik-Diff™ stain (catalog number 9990700, Epredia Shandon) to visualize cell morphology. Macrophages were identified based on their large size, irregular shape, abundant cytoplasm, and distinctive nuclear morphology.

### Collection of Human Alveolar Macrophage by BAL

Primary human alveolar macrophages were isolated from the adult human lungs (multiple donors) which were declined for transplantation^33^. Explicit approval for the use of donor lungs for research from each donor’s family was obtained by Donor Network West. All experiments using cadaver human lung tissue were approved by the UCSF Biosafety Committee. Donor information: Donor #1: White/Asian American male, 36 years old, never tobacco smoker; Donor #2: African American male, 55 years old, current tobacco smoker; Donor #3: White female, 61 years old, former smoker; Donor #4: White female, 54 years old, former smoker; Donor #5: Asian male, 50 years old, never tobacco smoker; Donor #8: White male, 68 years old, current tobacco smoker.

BAL fluid was obtained from a human lung lobe free of gross consolidation or injury as previously reported^34^. Briefly, BAL was obtained by injecting and withdrawing 500 mL of ice-cold PBS containing 2 mM EDTA into the left upper lobe bronchus, 50 mL at a time. Red blood cells in BAL were lysed with 1x RBC Lysis Buffer (catalog number 501129751, Invitrogen) according to the manufacturer’s instructions. Cells were pelleted at 300g for 4 min at 4°C. Pellets were resuspended in RPMI 1640 supplemented with 10% FBS and 1% P/S. BAL cells were plated in a 48-well plate at 50,000 cells per well and incubated for 2h at 37°C under humidified 5% CO_2_. Based on our past experience with both human and mouse BAL AMs, we used only the described collection and plating method as a selection process. This approach was chosen to minimize *ex vivo*, preserving their *in situ* conditions and preventing alterations in gene expression and responses to inflammatory challenges^21^.

### *Ex vivo* LPS Stimulation and Gut-Derived Metabolite Treatment to Human Primary Alveolar Macrophages

Two to 12 h after BAL cell seeding, non-adherent cells (non AMs) were washed away, and attached AMs (visualized by microscopy as the predominant cell population) were exposed to LPS (0, 25, or 500 ng/mL) with C3 (0, 0.3, or 3 mM) at 37°C under humidified 5% CO_2_. After overnight incubation, supernatants were collected for enzyme-linked immunosorbent assay (ELISA) analysis. For experiments where C2/C3/C4 were compared (Figure 7C), 200ng/mL LPS was used to stimulate BAL cells overnight in the presence of C2/C3/C4 (0, 0.3, or 3 mM) after 4h of pretreatment with the same type and dose of SCFA. For IPA experiments (Figure 7D), BAL cells were exposed overnight to LPS (10ng/mL) with C3 (0mM, 0.3mM and 3 mM) and/or IPA (0mM, 0.1mM and 1 mM).

### *Ex vivo* LPS Stimulation to Primary Alveolar Macrophages Obtained from Mice fed FF and FR Diets

BAL was performed as described above to isolate primary AMs from WT C57BL/6 mice fed ND, FF and FR diets for 2 weeks. Two to 12 h after BAL cell seeding, non-adherent cells were washed away, and attached AMs were exposed to LPS (0, 10, or 100 ng/mL) at 37°C in a humidified 5% CO_2_ atmosphere. Following overnight incubation, supernatants were collected for enzyme-linked immunosorbent assay (ELISA) analysis.

### *In vitro* LPS Stimulation and SCFA Treatment to Murine Alveolar Macrophage Cell Line

MH-S cells (10,000 cells/well in RPMI 1640) were seeded into 96-well plates supplemented with 10% FBS and 1% P/S. All cells were incubated at 37°C in humidified 5% CO_2_, and all treatments were administered 12-24h after cell seeding. All cells were pre-treated with C3 (0 or 1 mM) for 24h before exposure to LPS (10 ng/mL), with or without co-treatment with C3 (1mM). After 24h of stimulation, supernatants were collected for enzyme-linked immunosorbent assay (ELISA) analysis.

For IPA experiments (Figure 7D), cells were pre-treated with C3 (0, 0.3 and 3mM) and/or IPA (0, 0.1 and 1mM) for 24h followed by overnight exposure to LPS (50ng/mL) in the presence of the same concentrations of C3 and/or IPA. For experiments comparing C2, C3, and C4, BAL cells were pre-treated with the respective SCFA (C2, C3, or C4) at varying concentrations for 4h, followed by overnight stimulation with 200 ng/mL LPS in the presence of the same type and concentration of SCFA.

### *In vitro* and *ex vivo* nutritional IR injury

Primary alveolar macrophages (200,000 cells/well in RPMI 1640) or MH-S (200,000 cells/well in RPMI 1640) were seeded on a 48-well plate supplemented with 10% FBS and 1% P/S. All cells were incubated at 37°C in humidified 5% CO_2_ and all treatments were given 12-24h after cell seeding. All cells were exposed to LPS (200 ng/mL) with SCFAs overnight, and SCFAs were administered 4h before LPS stimulation.

*In vitro* and *ex vivo* nutritional IR conditions were performed as previously described^30^. Briefly, nutritional “ischemia” was established by first washing the respective cells twice with PBS and then replacing the medium with PBS for 1h. We did not create hypoxia for the period of nutritional IR injury because our *in vivo* ventilated IR model experiences minimal periods of hypoxia (as described above). After the *in vitro* “ischemia” period (1h), medium supplemented with 10% FBS was added for the “reperfusion” period (1h), and supernatants were collected at denoted times for ELISA analysis.

### Fecal Microbiome Transplant (FMT) in Germ-free (GF) Mice

Two groups of GF mice (n=10 per group) were colonized with microbiota via FMT. Donor homogenized colonic contents were obtained from specific pathogen-free (SPF) control mice fed FR or FF diets for two weeks. The FMT was performed by oral gavage 2–3 times over a two-week period, during which GF recipient mice were maintained on their respective FR or FF diets to match the dietary conditions of the donors. Following this period, the mice underwent lung injury procedures as previously described. Lung tissues (left and right lungs) were harvested, homogenized, and analyzed by ELISA to quantify lung injury markers. Additionally, colonic contents from recipient mice were collected and analyzed using 16S rRNA sequencing, along with the original FMT donor samples (FR and FF groups).

### Monocolonization with *Bacteroides thetaiotaomicron* (*B.theta*) in Germ-free (GF) Mice

A separate group of GF mice maintained on a ND was monocolonized with either propionate production capable *Bacteroides thetaiotaomicron* (*B. theta DeltaTDK*) or a propionate-deficient mutant strain of *B.theta (Delta1686-89)*. These strains were generously provided by Eric Martens and have been previously described^35^. Information regarding the nature of the strains has also been provided in **Table S3**. Colonization was achieved by administering 200 µL of bacterial suspension via oral gavage three times over a three-week period. Following monocolonization, the mice underwent *in vivo* lung IR injury. After 1h of reperfusion, lung tissues, plasma, colonic stool, and other samples were collected and analyzed using methods described elsewhere.

### Bacterial Lung Infections with *Streptococcus pneumoniae*

Three groups of WT C57BL/6 mice (n=10 each) were fed ND, FR or FF diets for 2 weeks. These mice had baseline measurements of body weight, rectal temperature, and oxygen saturation by pulse oximetry. Pulse oximetry was measured using the Mouse Ox+ cervical collar system (Starr Life Sciences, Oakmont, PA) by monitoring mice for 5 min for each measurement and taking the mean SpO_2_ for 10 consecutive seconds when mice were not active as calculated by the system software. Mice were inoculated intranasally with *S.pneumoniae* (1×10^8^ CFU) as previously reported ^36,37^ without the use of any antibiotics. Body weight and temperature were also measured at 12h, 24h, and 36h post infection, and oxygen saturation was measured at 36 h prior to collection of blood and BAL. BAL protein was measured by Pierce BCA Protein Assay (catalog number 23227, Thermo Scientific) and cell and differential counts performed. Postmortem bacterial titers of BAL were measured by serial dilution and counting colony-forming units on sheep blood agar plates.

### Precision cut lung slices (PCLS) preparation and culture

Preparation of PCLS was based on previous reports in the literature ^38,39^, with some modifications as described here. WT C57BL/6 mice were used to prepare precision-cut lung slices (PCLS). Anesthesia and analgesia were induced via IP injection of avertin at a dose of 250 mg/kg body weight, supplemented with IP buprenorphine (0.1 mg/kg body weight), to minimize pain and distress. A midline incision was then made to expose the thoracic cavity, and blood was thoroughly flushed from the vasculature by trans-cardiac perfusion with ice-cold PBS to ensure complete removal of blood from the pulmonary vasculature. The trachea was carefully exposed, and securely tied off with a suture and the mouse lung was inflated with 1.5 mL of pre-warmed 2% low melting point agarose (catalog number 16520100, Invitrogen) via intratracheal injection. After inflation, the inflated lungs were covered with chilled gauze and kept at 4°C for 10 min to allow the agarose to solidify, thereby preserving lung morphology before the lungs were dissected out.

Isolated lung tissue was processed using a compresstome VF-310-0Z (Precisionary, Natick, MA, USA) vibrating microtome. Lung tissue was affixed to the piston with cyanoacrylate adhesive, embedded in 4% low melting point agarose, and sliced into sections of 300 μm thickness. All PCLS were collected in chilled Hanks’ Balanced Salt solution (HBSS; catalog number 14185052, Gibco) and transferred to a pre-warmed 24-well plate at a rate of 2 slices per well. Shortly after, all PCLS were standardized into uniform round discs with a diameter of 4 mm. This was achieved using a sterile Biopunch tool with an inner diameter (ID) of 4.0 mm (TED PELLA, INC., Redding, CA). The Biopunch was cleaned and sterilized before use to ensure the integrity of the samples.

Each PCLS was carefully aligned on a flat, sterile surface, and the Biopunch was pressed firmly and evenly to create consistent, circular discs. Excess tissue around the discs was carefully removed to avoid contamination or interference with subsequent assays. This standardization step ensured uniformity across all samples, facilitating reproducibility and consistency in downstream analyses. The processed discs were immediately transferred to appropriate culture media to maintain tissue viability. The agarose was removed by washing the sections three times in pre-warmed DMEM/F12 medium (catalog number 21041025, Gibco) containing 10% FBS, 100 U/mL penicillin, 100 μg/mL streptomycin, and 1.5 μg/mL amphotericin B (catalog number 15290018, Thermo Fisher). Each wash involved incubating the slices in 500 μL of medium per well at 37°C with 5% for 1h.

After the final wash, the discs were cultured at 37°C with 5% CO_2_. The culture medium was replenished within the first 24h, and then slices were subsequently maintained in fresh DMEM/F12 medium supplemented with 10% FBS, 100 U/mL penicillin, and 100 μg/mL streptomycin. The AlamarBlue cell viability assay (Thermo Fisher) was used to confirm PCLS viability per manufacturer’s instructions. Viability assessments were repeated at intervals throughout the experiment.

### Quantification of Short-chain Fatty Acids (SCFAs)

SCFAs in mouse lung samples were quantified at the Metabolomics Core Facility at the University of Michigan. SCFA extraction was performed using an aqueous extraction solvent containing 3% (v/v) 1 M HCl and isotope-labeled internal standards (d7-buytric acid and d11-hexanoic acid). Samples were homogenized, centrifuged, and the supernatants subjected to liquid-liquid extraction with anhydrous diethyl ether. Following phase separation, the upper organic layer was transferred to autosampler vials for gas chromatography-mass spectrometry (GC-MS) analysis. GC separation was performed on an Agilent 6890 gas chromatograph with a ZB-Wax plus column (0.25 μm × 0.25 mm × 30 m) (Phenomenex, Torrance, CA). SCFA identification and quantification were achieved using a single quadrupole mass spectrometer (Agilent, 5973 inert MSD). Data were processed with Agilent Masshunter software (version B.06) and quantification was normalized to sample wet/dry weight for comparability across samples.

### Seahorse^TM^ Extracellular Flux Metabolism Analysis

The Agilent Seahorse XFe24 analyzer (Agilent Technologies, Santa Clara, CA) was used to assess metabolic responses in mouse PCLS and cell lines exposed to LPS and SCFAs. Mitochondrial function was evaluated using the Seahorse XF Cell Mito Stress Test, which measured oxygen consumption rate (OCR) to monitor mitochondrial respiration in real-time.

The MH-S (alveolar macrophages) and MLE-12 (murine lung epithelial cells) cell lines were cultured in RPMI 1640 and DMEM (catalog no. 11995065, Gibco), respectively, supplemented with 10% FBS and 1% P/S, at 37°C in a humidified 5% CO_2_ incubator. Cells were seeded onto an XFe24 assay plate (40,000 cells/well) one day before the assay and treated overnight with LPS (10 ng/mL) and C3 (0.1 or 5 mM).

Mouse PCLS were placed individually into XFe24 plate wells (one slice per well) in DMEM/F12 medium supplemented with 1% P/S and 1.5 μg/mL amphotericin B. Treatments, including LPS (100 ng/mL) and C3 (0.1 or 5 mM), were applied 24h after PCLS preparation. The sensor cartridge was hydrated in XF calibrant solution at 37°C in a non-CO_2_ incubator for at least 12 h.

On the assay day, XF assay medium was prepared by supplementing XF RPMI medium with 1 mM pyruvate, 2 mM glutamine, and 10 mM glucose, with the pH adjusted to 7.4. Working concentrations of compounds (1.5 μM oligomycin, 1 μM FCCP, and 0.5 μM rotenone/antimycin A) were prepared according to the XF Cell Mito Stress Test protocol. Cells and PCLS were gently washed with warm XF assay medium to remove residual treatment medium and incubated in the assay medium at 37°C in a non-CO_2_ incubator for 1h. Before analysis, cell confluence and PCLS integrity were confirmed microscopically.

The Seahorse analyzer measured OCR and metabolic parameters, including basal respiration, maximal respiration, ATP-coupled respiration, spare respiratory capacity, and metabolic potential.

### Quantitative real-time reverse transcription polymerase chain reaction (RT-qPCR)

TaqMan-specific inventoried gene primers for glyceraldehyde 3-phosphate dehydrogenase (GAPDH), beta actin (Actb), ribosomal protein L19 (Rpl19), interleukin (IL)-6, IL-1β, IL-10, IL-18, NOD-like receptor family, pyrin domain containing (NLRP) 3, chemokine (C-X-C motif) ligand (CXCL) 1, CXCL2 (Life Technologies, Carlsbad, CA) were used to measure the mRNA levels of these human or mouse genes in cells or lung tissue.

Lung tissue was homogenized using Tissue-Tearor (Biospec Products, Bartlesville, OK), and total RNA was isolated with TRIzol^®^ and the RNAeasy Mini Kit (catalog number 74104, Qiagen) for RNA purification. Reverse transcription was performed using the High-Capacity RNA-to-cDNA Kit (catalog number 4387406, Applied Biosystems) with 1 μg messenger RNA per reaction.

Quantitative real-time polymerase chain reaction was performed using TaqMan fast advance master mix (catalog number 4444557), TaqMan gene expression assay (reagent), and QuantStudio^TM^ 6 and 7 Flex Real-time PCR systems. Run method: Polymerase chain reaction activation at 95°C for 20 s was followed by 40 cycles of 1 s at 95°C and 20 s at 60°C.

The average threshold cycle count (Ct) value of 3 technical replicates was used in all calculations. The average Ct values of the internal controls (GAPDH, actb, and Rpl19) were used to calculate ΔCt values for the array samples. Data analysis was performed using the 2^-ΔΔCt^ method, data are presented as relative quantification (RQ) ^40,41^.

### Single-nucleus ATAC sequencing (SnATAC-seq) on Whole Lung Digests

Two groups of WT C57BL/6 mice (n=4 per group) were fed FR and FF diets for 2 weeks. At the conclusion of the dietary intervention, whole lungs were harvested under sterile conditions and placed in cold PBS to minimize enzymatic activity during transport. The lungs were then processed to isolate nuclei for single-nucleus ATAC sequencing (snATAC-seq) using the Chromium Single Cell ATAC Library & Gel Bead Kit (10x Genomics, Pleasanton, CA), following the manufacturer’s protocol. Raw sequencing data were analyzed using the Cell Ranger ATAC pipeline (10x Genomics) for alignment, barcode assignment, and peak calling. Subsequent analyses included dimensionality reduction, clustering, and identification of regulatory elements and cell-type-specific chromatin accessibility. Differential chromatin accessibility analysis between the FR and FF diet groups was performed to identify potential diet-induced epigenetic modifications in lung tissue.

### Sandwich Enzyme-Linked Immunosorbent Assay (ELISA)

Cytokines and chemokine concentrations in cell culture supernatant, plasma, and lung homogenates were quantified using DuoSet or Quantikine ELISA kits (R&D Systems, Minneapolis, MN). All assays were performed according to the manufacturer’s protocol. Standard curves were generated for each assay using serial dilutions of known concentrations of the target analyte. Sample cytokine and chemokine concentrations were calculated from the standard curves and expressed as the mean ± standard deviation (SD).

### Correlation Analyses of 16S rRNA Sequencing Data, Metabolomic Data, and Immune/Inflammatory Markers

The size of the sequenced paired-end libraries ranges from 5,976 bp to 285,864bp, representing a total of over 15 million 151bp reads. The resultant paired-end 150bp data were demultiplexed by the unique barcode and converted into fastq format via the bcl2fastq software. Forward and reverse reads were processed separately, and quality filtered using the DADA2 software package version 1.16.0. in R.4.1.2. Reads having more than two expected errors and/or <150 base pairs in length were removed. Error rates of the filtered dereplicated reads were estimated using 100,000 sequences. Paired reads with a minimum overlap of 25 base pairs were merged to obtain the full denoised sequences.

Amplicon sequence variants (ASVs)were inferred exactly, resolving variants that differ by as little as one nucleotide. ASVs determined to be chimeric were discarded. Taxonomic assignment was performed using the naïve Bayesian classifier method utilizing the SILVA v138 16SrRNA reference database.

Additionally, taxa known to be common reagent contaminants and observed in greater than 15% of negative extraction control were discarded. The maximum read count of each remaining ASV in any single negative extraction control was subtracted from the reads counts of that ASV for each sample. Read counts for ASVs which summed across all samples that were less than 0.001% of the total read count for the dataset were discarded to minimize noise in the dataset. Sample read numbers were rarefied on minimum sequencing depth obtained for this study (mentioned below under Read-Depth section).

Reads were processed through Qiime2 (version 2020.8.0)^42^:low-quality reads and sequencing adaptors were removed using Cutadapt^43^, and sequencing errors were corrected using Dada2^44^ using custom parameters (--p-trunc-len-f 150--p-trunc-len-r 140). Taxonomic assignation of resulted ASVs was done using SILVA trained database (version 138-99)^45^ based on scikit-learn’s naïve Bayes algorithm^46^. For the subsequent analyses (except for alpha-diversity calculations), the abundance of each taxon present in a sample was normalized using the relative method to allow sample-to-sample comparison. Results were deep analyzed using the Phyloseq package (version 1.34.0)^47^ as for the taxonomic and alpha diversity analyses. Statistical analyses were performed using rstatix (version 0.7)^48^ and figures were plotted using the ggplot2 package (version 3.3.5)^49^. Principal coordinate analyses (PCoA) were performed with the Vegan package (version 2.5-7)^50^ on the Bray-Curtis dissimilarity matrices constructed from the relative abundance of ASVs. The communities that emerged were verified using a PERMANOVA test and the confidence intervals were plotted with 95% and 97% confidence limits, using the standard deviation method. Spearman correlation analyses were performed to determine associations between metabolites and microbiota species using the R package energy (version 1.7-8).

Procrustes analysis were performed to determine the relationship between Euclidean distances lung protein/transcipt/M_data and Bray-Curtis distances of microbiome. The length of lines on Procrustes plots indicate dissimilarity within-subject of the microbiome and metabolome. The smaller the sum of squares values (M-square), the identical the matrices are, and vice-versa.

Corr_Procrustes_rotation is a corr value of the Procrustes rotation. Procrustes analysis results includes the Procrustes sum of squares, correlation Procrustes rotation and p-values for relationship between profile1 (microbiome) and profile2 (protein, Transcripts and M_data sub-groups) data sets. The metabolomic data was sub-grouped into M_data_pb (portal vein plasma), M_data_p (peripheral plasma), and M_data_RL (right lung tissue) and then there correlation was established with microbiome.

### Gut Permeability Measurement

Gut permeability was assessed by measuring Lipopolysaccharide Binding Protein (LBP) levels in mouse plasma using the Mouse LBP Immunoassay Kit (catalog number EEL115, Invitrogen). Plasma samples collected from mice fed FR and FF diets were analyzed using ELISA according to the manufacturer’s instructions.

### Statistical Analyses

Data in the figures are expressed as mean +/- SD. Data from *in vivo* studies comparing two conditions were analyzed using 2-tailed nonparametric Mann–Whitney analyses. For bacterial infection in vivo experiments, 2-way ANOVA (body temperature) and mixed-effects analysis with multiple comparisons (SpO2) were used. Aseesa’s Stars analysis tool (www.aseesa.com) was used to analyze and generate principal analysis donut and scatter plots, and correlations ^51^. Also shown are linear trendlines, or an nth-degree polynomial trendline if its goodness-of-fit is either 50% greater than, or if it explains at least half of the variance not explained by the (n – 1)th-degree polynomial. R^2^, r and p denote goodness-of-fit, Pearson’s correlation coefficient and significance of the correlation, respectively. Data from *in vitro* studies comparing two conditions were analyzed using standard Student’s t-test with equal SD to generate *P* values. GraphPad Prism was used for statistical analysis (GraphPad Software, La Jolla, CA). P values < 0.05 were considered significant. P values are presented in the figures as follows: *< 0.05; **< 0.01; ***< 0.001; ****< 0.0001. *In vitro* experiments were repeated 2 or more times, and representative are data shown. The number of animals used in *in vivo* experiments are described elsewhere within the methods section. One cage from the FF->FR group (FR 1wk) with lung IR surgery was excluded due to the presence of *E. coli/Shigella* in the fecal microbiome composition indicating a potential GI infection in this one cage.

## RESULTS

### Fiber-rich (FR) diet results in metabolically reprogrammed BAL alveolar macrophages (AMs) in uninjured lungs

In our prior work, we reported that direct pulmonary administration of SCFA diminished lung immunity, that altering the gut microbiome with antibiotics enriched for SCFA-producing *lachnospiraceae* species and reduced sterile lung injury^27^, and that alveolar macrophages mediated this injury^30^. Therefore, we set out to investigate the impact of FR diets, which are expected to naturally generate SCFAs after fiber fermentation by gut microbiota, on the transcriptional programming of AMs. We fed mice a FR diet containing 35% pectin for 2 weeks and used a FF diet (0% fiber) as our control (schematized in **Figure 1A**). We chose pectin over inulin and other fiber-containing prebiotics for its known ability to reliably generate fecal SCFAs and based on previous published work^9,24,52^.

**Figure 1.**
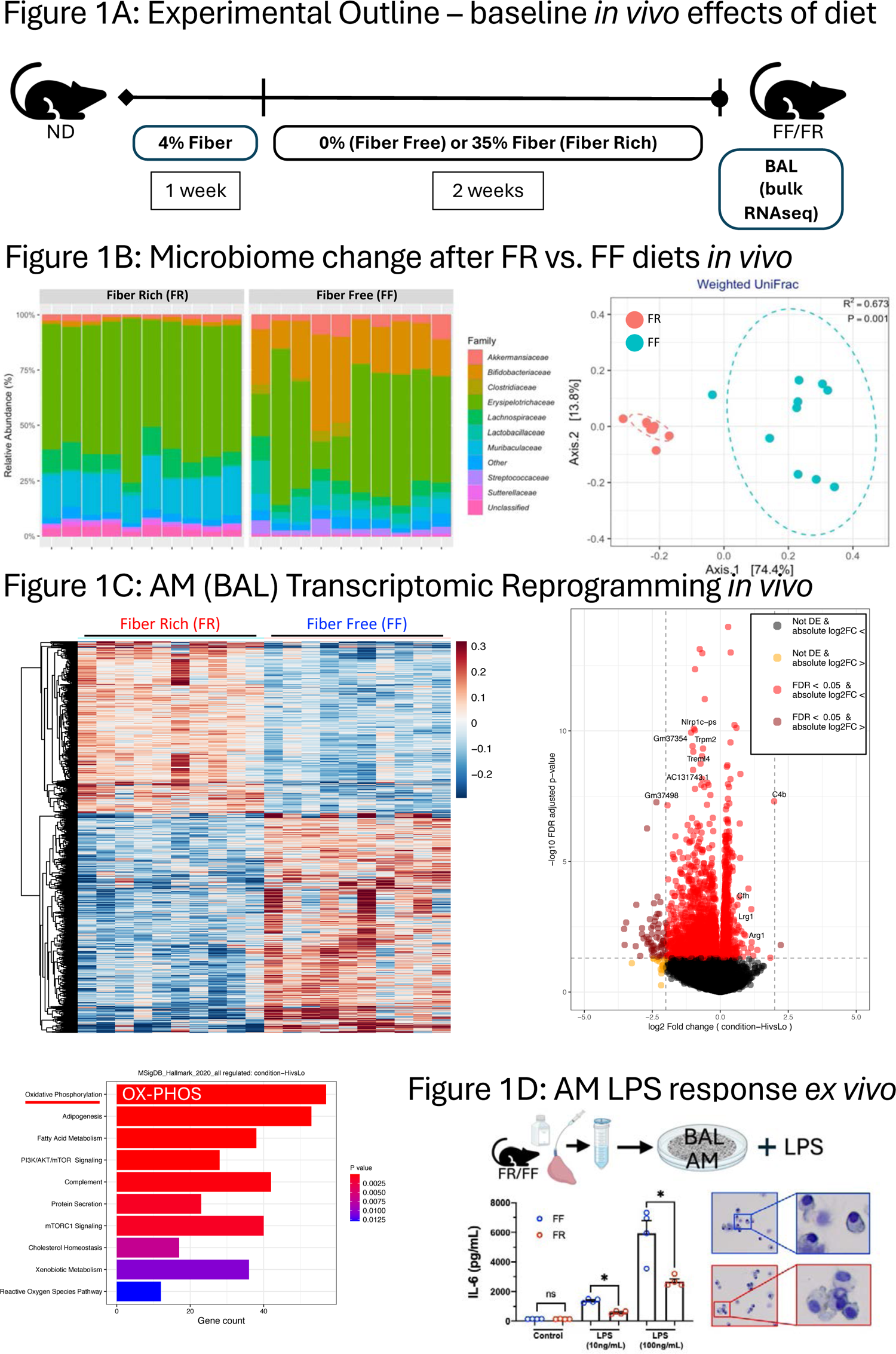
FR diet changes gut microbiome composition and reprograms BAL AMs towards oxidative phosphorylation (pre-injury). (A) Experimental outline for fiber-free (0% fiber/FF) and fiber-rich (35% fiber/FR) dietary intervention switched from normal fiber (ND), followed by bronchoalveolar lavage and bulk RNAseq analysis. (B) Family level changes in gut microbiome comparing mice on FR and FF groups (after 2 weeks of dietary intervention) (left); Weighted UniFrac beta diversity analysis (right). (C) Heat map of the differentially expressed genes between these two groups (top, left); Volcano plot of differentially expressed genes (FR/FF) (top, right) with Arg1 highlighted. Differential gene expression analysis using the MSigDB Hallmark (bottom, left) reference database shows enrichment of key pathways (OX-PHOS highlighted and underlined in red) in the FR group compared to the FF group. n=10 mice in each group. (D) IL-6 response in AMs from mice on fiber-rich (FR) and fiber-free (FF) diets after being challenged with LPS (10 ng/mL and 100 ng/mL, left). P values are represented comparing IL-6 in AM cells between FF vs FR diet group: *< 0.05; Microscopic images (right) of cytospin preparations from AMs from FR (red box) and FF (blue box) mice. These images provide a visual representation of the AM morphology.

To confirm that our dietary intervention resulted in a change in the resident gut microbiota, we performed 16S rRNA sequencing of fecal contents and observed distinct taxa profiles between the two groups, which clustered separately by beta diversity analysis (**Figure 1B, left and right**). Performing bulk RNAseq on the BAL cells, we observed airway cells (predominantly AMs) from mice that were exposed to FR diets for 2 weeks were strikingly different from those obtained from FF diet mice (**Figure 1C, top left**). Of note, our analyses did not reveal any meaningful differences in the transcript levels of AMs (CD11c and SiglecF) or neutrophil markers (Ly6G and CD11b) expressed alleviating concerns that these fiber diets might affect the number of AMs present in the mouse lungs or that the FF diet might cause baseline lung inflammation (data not shown). In addition, we observed no evidence of differences in gut leakiness between these two groups (plasma LPB levels, **Figure S2**). Gene transcript enrichment analysis revealed a handful of differentially expressed genes, most notably Arg1, that were enriched in the FR group (**Figure 1C, top right**) suggesting that FR AMs may be skewed toward the M2 (anti-inflammatory) and away from the M1 (inflammatory) macrophage phenotypes. Pathway enrichment analyses also showed that oxidative phosphorylation (OX-PHOS) was the #1 (MSigDB) and #4 (KEGG) most highly enriched pathway in the FR group (**Figure 1C, bottom left** and data not shown) consistent with M2 programming^13^. Other gene set enrichment analyses also showed OX-PHOS as a top enriched pathway as well as inflammatory response, complement, and interferon alpha response. The interferon gamma response pathways was negatively enriched (**Table S1**).

### FR diet mice BAL alveolar macrophages are less responsive to LPS injury

To investigate the effect of dietary fiber content on alveolar macrophage (AM) responses to sterile LPS injury, BAL AMs (enriched by adherence plating) were stimulated with two doses of LPS (10 ng/mL and 100 ng/mL). AMs from FF mice produced significantly higher IL-6 levels compared to FR AMs (**Figure 1D, left**). Cytospins were performed to image the BAL populations and these did not show any morphological differences between the AMs (**Figure 1D, right**).

### Fiber-rich (FR) diet results in reduced sterile lung IR injury with distinct regulation of IL-1β and IL-18 responses

Next, the mice that were exposed to FR (1 or 2 weeks, henceforth referred to as FR 1wk and FR 2wks) or FF (2 weeks) diets were subjected to lung IR injury (1h left lung ischemia, 1h reperfusion) and were then examined for <u>immediate-early</u> lung injury responses (schematized in **Figure 2A**). This early time point after IR injury (1h post-reperfusion) has been validated by our previous work to focus on the first wave cytokine and chemokine release *prior* to any histopathologic signs of lung injury (which peak at 3h post-reperfusion and include edema, hyaline membrane casts, and neutrophil infiltration)^30^.

**Figure 2.**
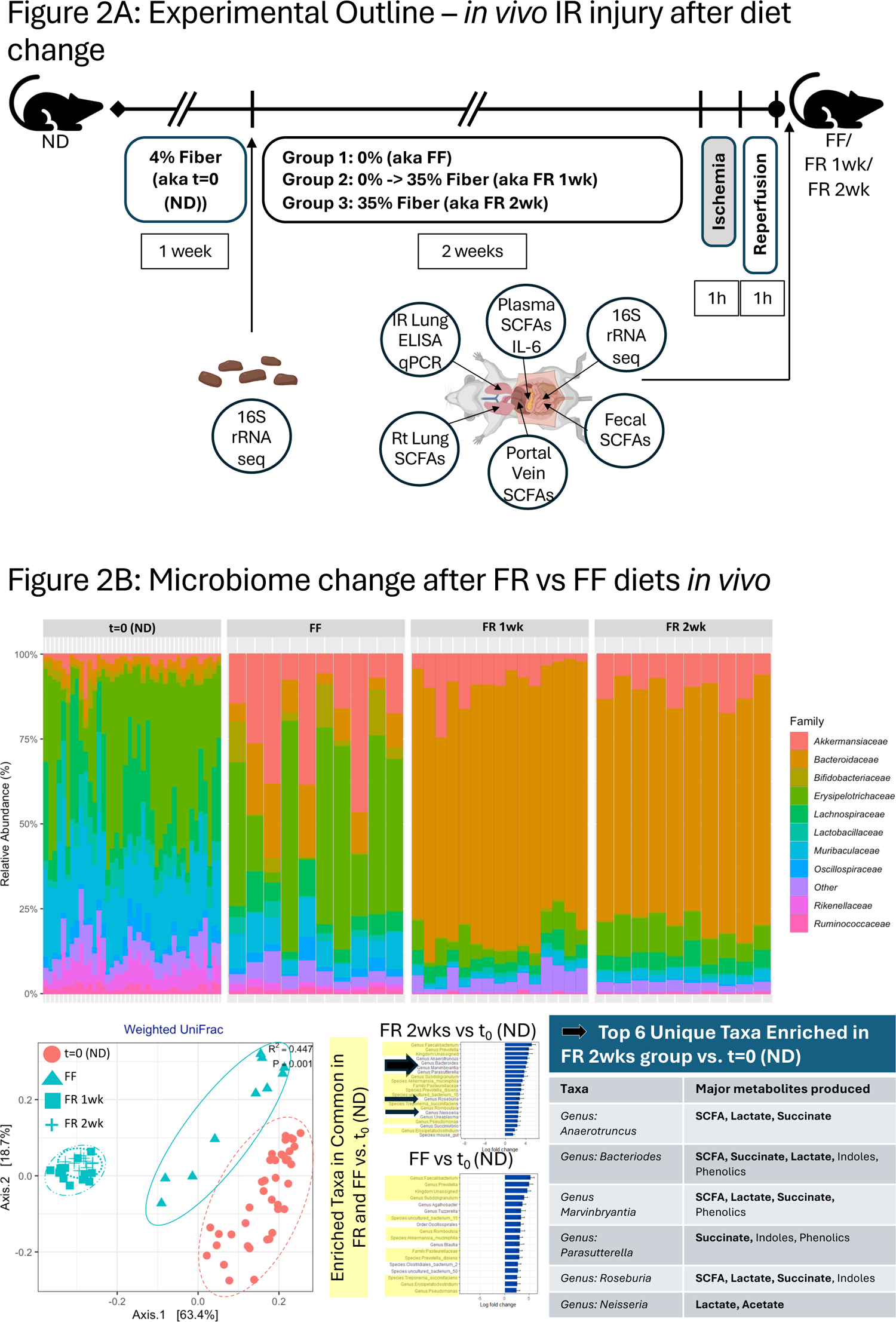
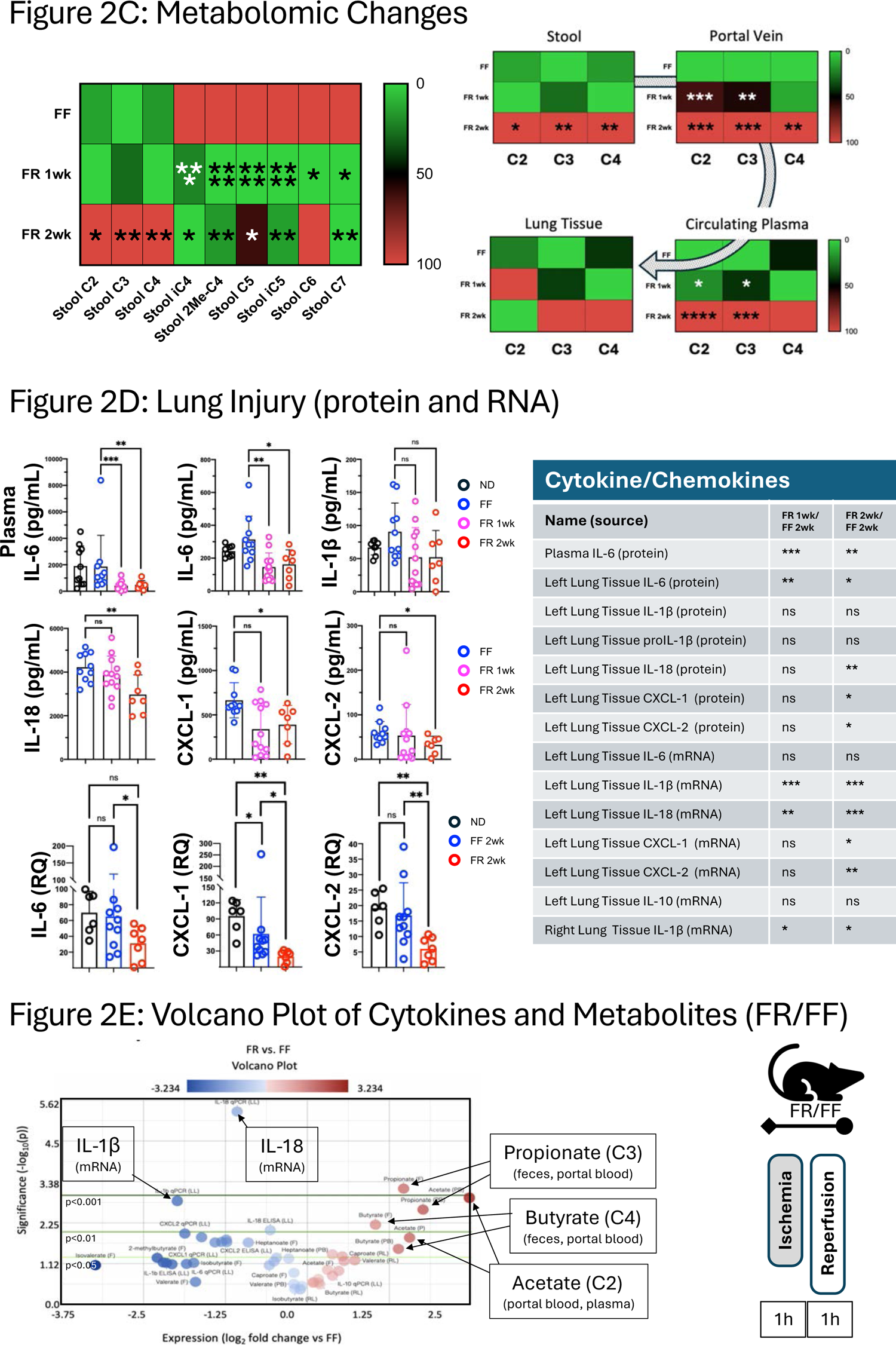
FR diet increases levels of SCFAs and blunts lung IR injury-related inflammatory responses with distinct gut microbiome regulation of IL-1β and IL-18. (A) Experimental outline for FF and FR dietary intervention, followed two weeks later by lung ischemia reperfusion (IR) injury. Different measurements at the start of the fiber diet intervention and after the IR injury are shown. (B) Family level changes in gut microbiome comparing mice on normal (2.4% fiber) chow (t=0, prior to dietary intervention; n=40 mice), FF, and FR groups (top). Beta diversity clustering of mouse groups showing FF, FR 1wk, FR 2wk, and t=0 (ND) (bottom left). Positively enriched taxa in FR group vs. baseline (t=0 (ND)) and FF vs. baseline (bottom middle); taxa highlighted in yellow were common to both FR 2wk and FF diets vs. baseline. The top 6 taxa uniquely enriched in the FR 2wk group and the major metabolites produced are listed with SCFAs and metabolites that can be easily converted to SCFAs highlighted in bold (bottom right). (C) Heatmaps showing changes in SCFA levels in feces between the three dietary groups (left). Focus on C2-C4 comparison between dietary groups in stool, portal blood, plasma, and right lung tissue (right). P values are represented comparing FF vs FR 1wk and FF vs FR 2wk as follows: *< 0.05; **< 0.01; ***< 0.001; ****< 0.0001. (D) Post-IR injury inflammation: plasma IL-6 and lung tissue inflammatory markers measured in normal diet (ND), FF, FR 1wk, and FR 2wk groups (top and middle right 6 panels). IL-6, CXCL-1, and CXCL-2 mRNA levels post-IR injury in ND, FF and FR 2wk (bottom right three panels) groups. Summary of all plasma and lung tissue analytes measured and level of significant for values different between FR and FF groups (right table). P values: *< 0.05; **< 0.01; ***< 0.001. (E) Volcano plot showing increased and decreased metabolites, and cytokine/chemokine in all tissue and fluid samples examined in FR group versus FF. Acetate (C2) in portal blood was the most increased measure and isovalerate in feces was the most decreased measure in FR diet group. Other key significantly changed factors are highlighted in black boxes. Green lines denote significance cut off levels (lighter to darker, p<0.05, p<0.01, and p<0.001).

As shown in **Figure 2B**, the gut microbiota profile was strongly influenced by the dietary change, with 1 and 2 weeks of FR diet resulting in a distinct profile compared to FF diet and the baseline (t=0, normal diet (ND) – 2.4% fiber) with the FR and FF groups well separated in beta-diversity analyses (**Figure 2B, bottom left**). Interestingly, the FF profile was clustered closer to the baseline (t=0) ND microbiome profile. We observed an expansion of the *Bacteriodetes* phyla and a strong reduction of the *Firmicutes:Bacterioides* ratio in the FR groups (data not shown). The FF diet resulted in an enrichment of *Verrucomicrobiota* (including *Akkermansia*), in agreement with other reports^53^.

Analysis of the enriched bacteria in the FR 2wks group over baseline (t=0) ND microbiota revealed unique taxa present at >3 fold magnitude increase including members of the genera *Anaerotruncus, Bacteriodes, Marvinbryantia, Parasutterella, Roseburia, and Neisseria* (**Figure 2B, bottom middle)**. These taxa’s reported metabolic output primarily consisted of SCFAs as might have been expected given the composition of the FR diet (**Figure 2B, bottom right**). Other metabolites produced by these taxa included lactate and succinate, which are also known to be converted into SCFAs (butyrate and propionate, respectively) by gut microbiota through bacterial cross-feeding^54–58^.

We next examined FR and FF mouse cecal contents for SCFA levels and observed the expected increase in C2 (acetate), C3 (propionate), and C4 (butyrate) in the FR groups, especially in the FR 2wks diet group (**Figure 2C, left**). Interestingly, we observed an increase in medium and branch chain FAs (iC4, 2Me-C4, C5, iC5, C6, and C7) in the FF group. We then followed the transit of SCFAs from the colonic contents (stool) to the portal vein, to circulating plasma, and finally to lung tissue (**Figure 2C, right**). The levels of SCFAs in lung tissue were elevated in the FR groups but not significantly different, and as we later propose, we believe that this is due to SCFA consumption by injured lung cells.

We observed that FR diet mice had significantly reduced lung IR injury as measured by lower systemic and local IL-6 levels (compared to both FF and ND mice). Lung tissue levels of IL-1β and IL-18 were also reduced. These findings were confirmed at the transcript level with reduced levels of IL-6, CXCL-1 and CXCL-2 mRNA levels in injured lung tissue from the FR diet mice, particularly in the 2 weeks FR diet group (**Figure 2D, left**). A summary of all the metabolites and inflammatory markers that were differentially present in the FR groups (compared to the FF group) is shown in a table (**Figure 2D, right**) as well as in the volcano plot representation in **Figure 2E**. Overall, acetate and propionate were the factors most elevated in the portal blood, feces and plasma, while lung IL-18 and IL-1β mRNA were the most significantly decreased (all measurements included in **Figure S3**).

### Association of SCFA levels and Gut Microbiota with IL-1β and IL-18 injury responses after lung IR

Next, we performed multi-omic analyses of our combined multivariate datasets consisting of multiple measurements (gut bacterial 16S rRNA gene sequences, SCFA metabolomics, targeted transcriptomics, and targeted proteomics) from individual mice (3 different diets) with the goal of uncovering potentially important correlations between our measured factors.

To do this, we first examined the individual datasets from the three dietary intervention groups and the combined data from all three groups. Despite the relatively small number of mice (n=30) and the targeted nature of our inflammatory gene measurements, we observed that lung IL-1β mRNA levels negatively correlated at the phylum level with fecal proteobacteria levels (p=10^-6.4^) and lung IL-18 mRNA positively correlated with fecal firmicutes levels (p=10^-10.1^) (**Figure 3A, top two graphs**). In addition, lung IL-18 mRNA levels positively correlated with fecal heptanoate (C7) levels (p=10^-4.8^) and negatively correlated with fecal propionate (C3) levels (p=10^-7^) (**Figure 3A, bottom two graphs**). Other significant correlations are shown in **Table S2** and **Figure S4**.

To further elucidate the relationships among these datasets, we applied Procrustes multivariate dataset comparison tests. These revealed the expected correlations (**Figure 3B, top left table**) between the gut microbiome 16S rRNA gene sequenced dataset and the SCFA metabolomic datasets (consistent with gut microbiota being the primary source of SCFA production). Additionally, we also found highly significant correlations between the 16S rRNA sequencing dataset and the lung inflammatory marker transcriptomic dataset, as well as with the lung injury protein dataset. The visual representation of the Procrustes analysis highlighted close relationships between the 16S rRNA dataset and transcript dataset (**Figure 3B, top right**), as well as between the 16S rRNA dataset and the other datasets (**Figure 3B, bottom 5 panels**). These close relationships were particularly evident for the FR group datasets (shorter line distances on the Procrustes plots and highlighted by the black dotted boxes) suggesting that these two datasets were highly correlated.

**Figure 3.**
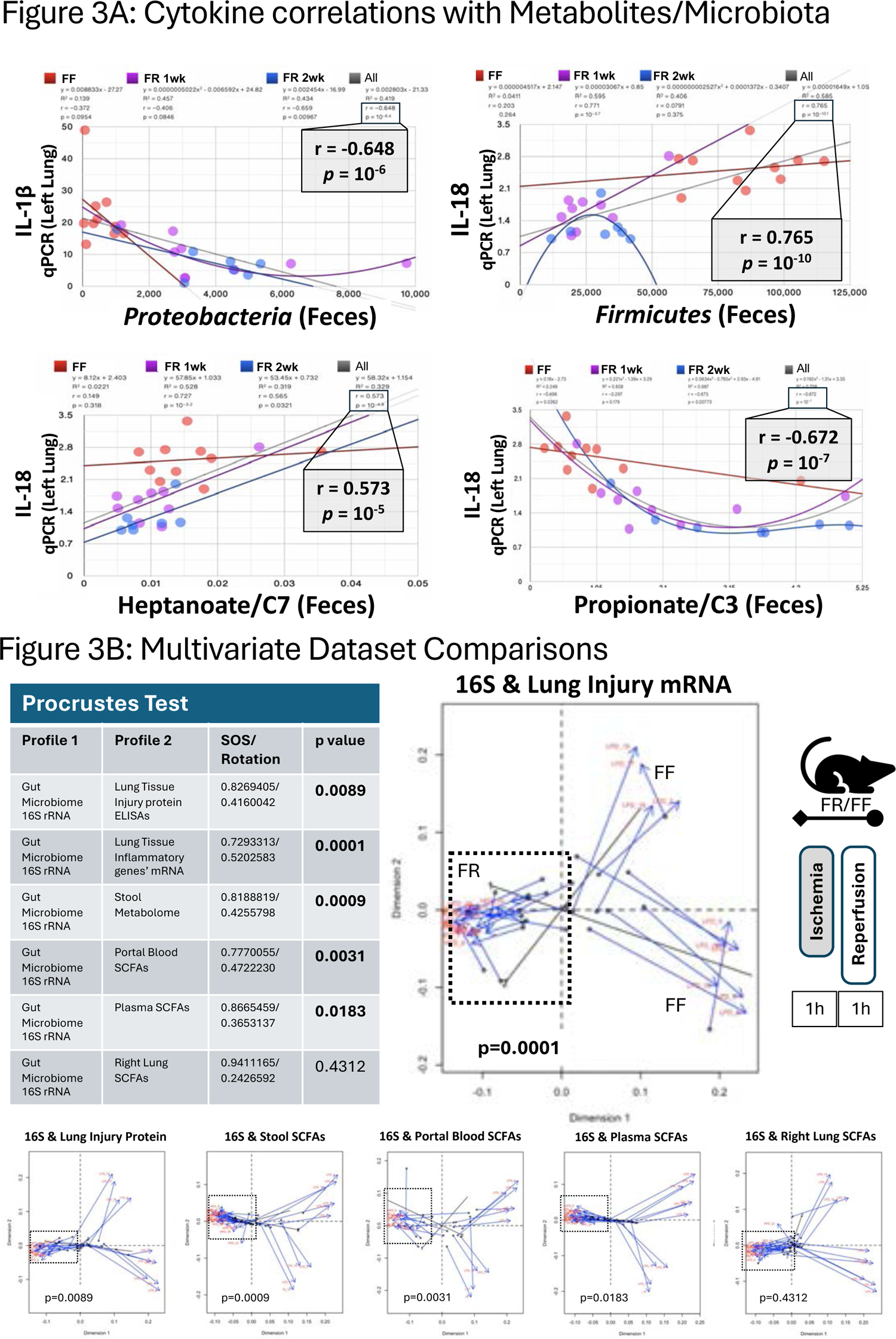
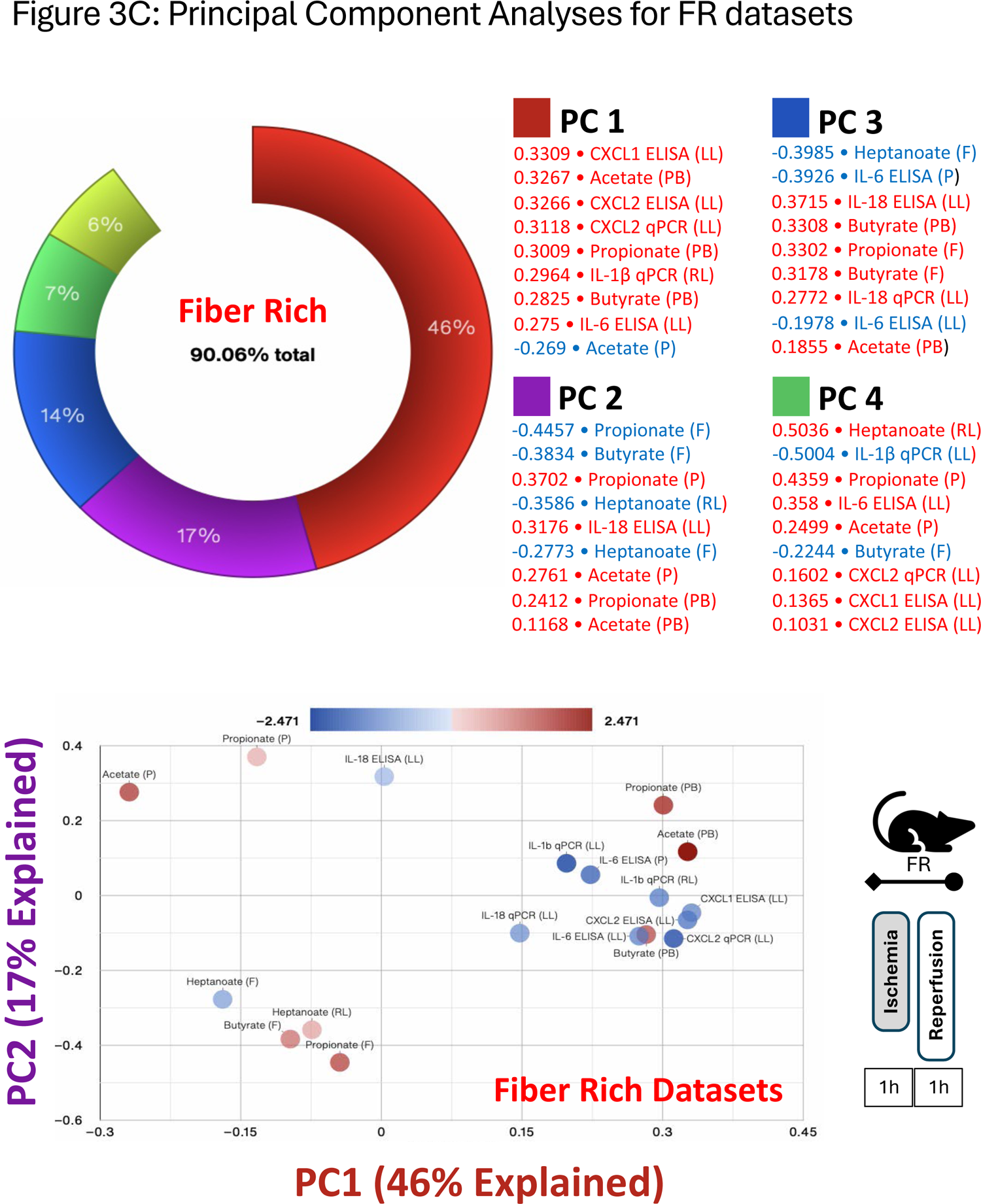
Enrichment of SCFAs and reduction of lung inflammatory markers after FR diet and post lung-IR injury. (A) IL-1β levels in left lung as measured using qPCR correlated with fecal proteobacteria levels, and IL-18 levels correlated with firmicutes (top left and right), heptanoate, and propionate levels (bottom left and right). All three groups are included separately (FF - orange, FR 1wk - purple, and FR 2wk – blue lines) and combined (gray line). Individual r and p values for each group are shown. (B) Procrustes test analyses results correlating gut microbiome 16S rRNA sequencing dataset with inflammatory transcriptomic/proteomic and SCFA metabolomic datasets. Significant data set correlations’ p values shown in bold (top left). Visual representation of Procrustes analyses shown at top right (16S rRNA sequence dataset vs. lung tissue injury transcript qPCR dataset) and below with dotted black boxes denoting the combined FR 1wk and FR 2wk data showing the strongest correlations. Bottom row (left to right) – 16S rRNA sequence dataset vs. lung tissue injury protein ELISA dataset; 16S rRNA sequence dataset vs. stool SCFA metabolome dataset; 16S rRNA sequence dataset vs. portal blood SCFA metabolome dataset; 16S rRNA sequence dataset vs. plasma SCFA metabolome dataset; 16S rRNA sequence dataset vs. right lung SCFA metabolome dataset. (C) PCA Donut chart showing contributions of the five principal components accounting for 90+% of FR group’s datasets variance and the 9 most correlated measures for each of the first 4 principal components are listed (top). Measures in red/blue text are positively/negatively correlated with the respective component. PC1 and PC2 shown in a 2D PCA plot (bottom).

Principal component analysis (PCA) performed on the 2-week FR dataset focusing on metabolites (from all sample sources), protein and transcript data (from lung and plasma) revealed five modules or principal components that explained 90% of the variation observed in this dataset (**Figure 3C, top**).

Modules combining SCFAs and inflammatory markers were observed suggesting potential relationships between these factors within the gut-lung axis (as shown in the PC1 vs PC2 plot, **Figure 3C, bottom**).

### Transfer of the Fiber-driven Lung Injury Phenotype with Fecal Microbiota Transplantation (FMT)

Next, we wanted to determine the importance of the gut microbiome in driving the impact of dietary fiber on lung immunometabolic programming and injury responses. We performed FMTs in germ-free (GF) mice using colon contents from SPF mice fed FR or FF diets for 2 weeks. After gavaging the mice with these microbial communities, and allowing the gut microbiome to colonize the intestinal systems of the GF mice, we subjected the two groups to lung injury.

We observed that the FMT from the FR group was able to suppress inflammatory responses compared to the FMT from the FF group both in the directly injured left lung (**Figure 4A, top**) as well as in the indirectly injured right lung – which interestingly showed similar levels of injury as the directly injured left lung (**Figure S5**). To confirm that the gavage FMT input matched the gut microbiota present 2 weeks later, we performed 16S sequencing (**Figure 4A, bottom left**) and beta diversity analyses, which showed that the FR and FF FMT inputs clustered closely with the microbiota from GF mice colonized with FR and FF FMTs (**Figure 4A, bottom right three panels**).

**Figure 4.**
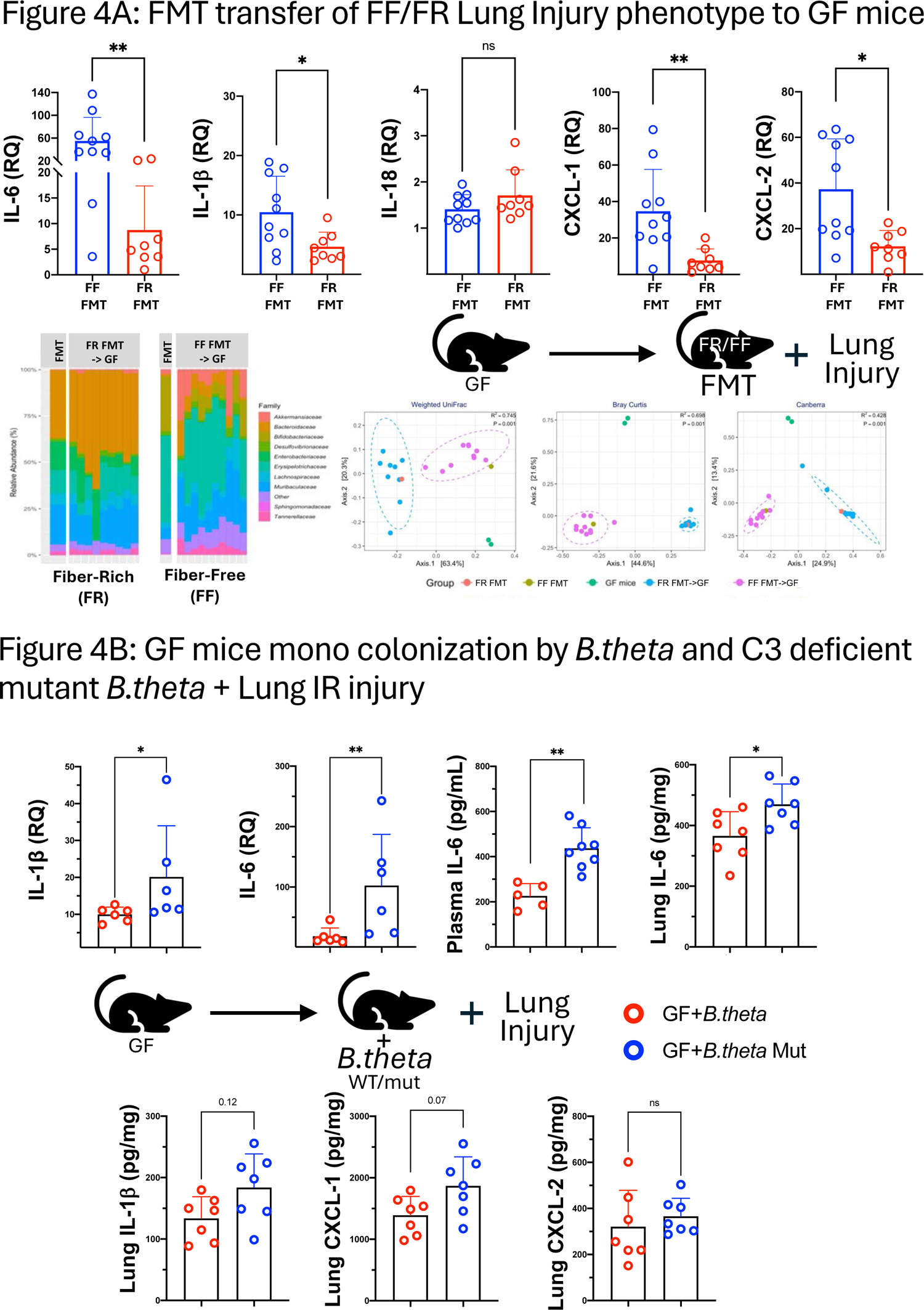
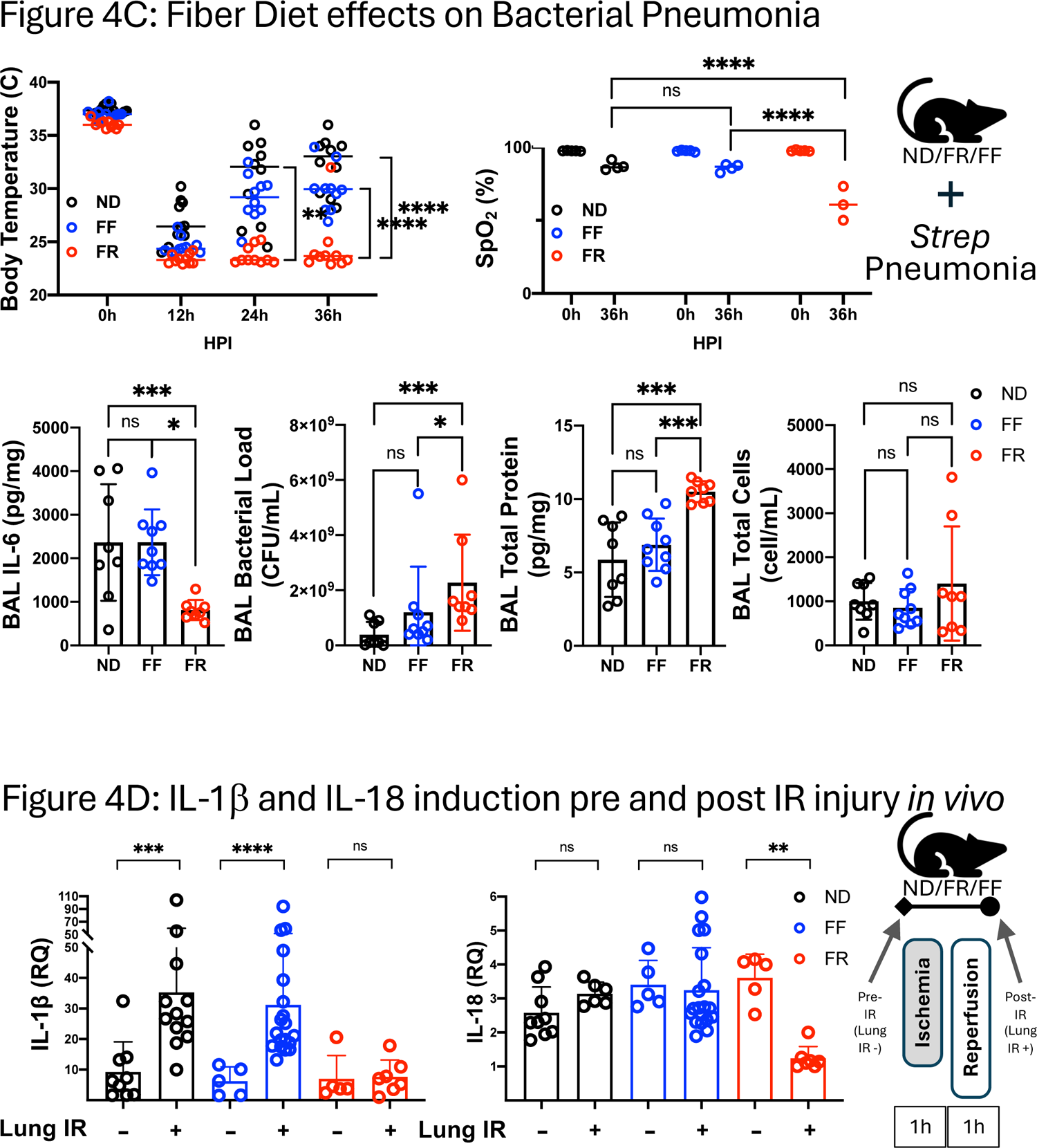
FF and FR phenotypes can be transferred with FMT and differentially affect bacterial pneumonia course. (A) GF mice received FMT using FF and FR homogenized fecal content prior to lung injury. Levels of injury in left lungs shown as measured by qPCR (top). Family-level composition of input FMT inoculum as well as colonized gut microbiota after FMT by 16S rRNA sequencing (bottom left); Weighted UniFrac beta diversity analysis of input and stool microbiota (bottom right). (B) GF mice received either FMT oral gavage inoculation with either *B.theta* or mut *B.theta* (mutant unable to produce propionate) for 3 weeks (1 gavage/week) and then subjected to lung IR injury. Levels of injury in left lungs as measured by ELISA and qPCR. (C) *S.pneumoniae* intranasal (IN) infection of ND, FF and FR diet mice with body temperature (top left), oxygen saturation/SpO2 (top right) with data shown for hours post infection (HPI) time points. P-values represent the significance values for the comparisons of the FR diet group against both the FF and ND groups. BAL IL-6 levels, BAL bacterial load, BAL total protein, and BAL total cells were also measured (bottom left to right). (D) Regulation of IL-1β and IL-18 pre- and post-lung IR injury comparing normal diet (ND) to FF to FR (2 wk) diets. This figure includes data from multiple experiments using the 3 different diets +/- lung IR injury. P values: *< 0.05; **< 0.01; ***< 0.001.

### Lung Injury responses depend on the ability of gut microbiota to produce propionate

We confirmed the importance of propionate production in controlling lung IR injury by inoculating more GF mice with a commonly prevalent propionate-producing gut microbe from the *Bacteriodes* genus, namely *B.thetaiotomicron* (aka *B.theta*). We used either *B.theta* (that produces propionate) or mutant *B.theta* (unable to produce propionate)^35^ and observed that the mice colonized with the mutant *B.theta* had elevated lung injury markers post-IR injury (**Figure 4B**) supporting our model that that the bacterial production of propionate was important for controlling or reducing lung injury responses.

### Fiber-rich (FR) diet influences Bacterial Pneumonia Outcomes and Associated Lung Damage

To investigate whether FR diets could positively or negatively affect host defense against lung infection, we used a bacterial pneumonia model (10^8^ CFU *S.pneumoniae* infection without any antibiotics) in mice after they were fed 2 weeks of FR, FF diets with normal diet (ND) as a control. We observed that the FR group had significantly lower body temperature and oxygen saturation levels 36h after pneumococcal infection, but no significant change in body weights post infection (**Figure 4C, top** and data not shown). Further analysis of these mice, revealed that FR mice had reduced BAL IL-6 levels, elevated BAL bacterial load and increased BAL total protein without any difference in BAL cell number or composition (**Figure 4C, bottom left to right** and data not shown), consistent with the FR mice having the combination of the least controlled infection and worst lung injury compared to FF and ND mice.

### Reprogramming of lung injury responses by FR diets does not occur at baseline via modulation of IL-1β and IL-18 resting levels

Since we had previously reported that IL-1β and IL-18 are key early cytokines for the generation of lung injury under different contexts, including IL-1β specifically for lung IR injury^18,21^, we delved deeper into the regulation of mRNA levels of these two cytokines before and after IR injury (pre-IR and post-IR). We had initially predicted that FR diets would reduce the levels of resting IL-1β mRNA within the lung prior to injury. Instead we observed the opposite – baseline levels of IL-1β and IL-18 were essentially identical in uninjured ND, FF and FR mice. Strikingly, we observed that these cytokines were distinctly altered in IR-exposed mice 2h post-injury, with expected IL-1β mRNA induction seen in the ND and FF groups but not in the FR group (**Figure 4D, left**). IL-18, on the other hand, was suppressed 2h post-injury in the FR group compared to the FF and ND groups (**Figure 4D, right**). These observations that ND and FF groups had similar post-injury responses was supported by earlier observations that their other injury responses and microbiomes were more similar to each other than to the FR group (**Figure 2B, bottom left** and **Figure 2D, left**).

### SCFA regulate lung inflammatory responses and metabolic programming in alveolar macrophage in vitro

If the combination of a FR diet, production of SCFAs and transit of SCFAs to the lungs *in vivo* could result in the immunometabolic reprogramming of AMs from M1 to M2 phenotypes, we wanted to examine if this reprogramming would happen as well *in vitro* in the presence of propionate (C3). We stimulated AMs (MH-S cell line) with LPS alone or in the presence of increasing C3 concentrations and observed the clear change from an M1-like to M2-like phenotype with inflammatory IL-6 levels decreasing while IL-10 levels increased dramatically (**Figure 5A**).

**Figure 5.**
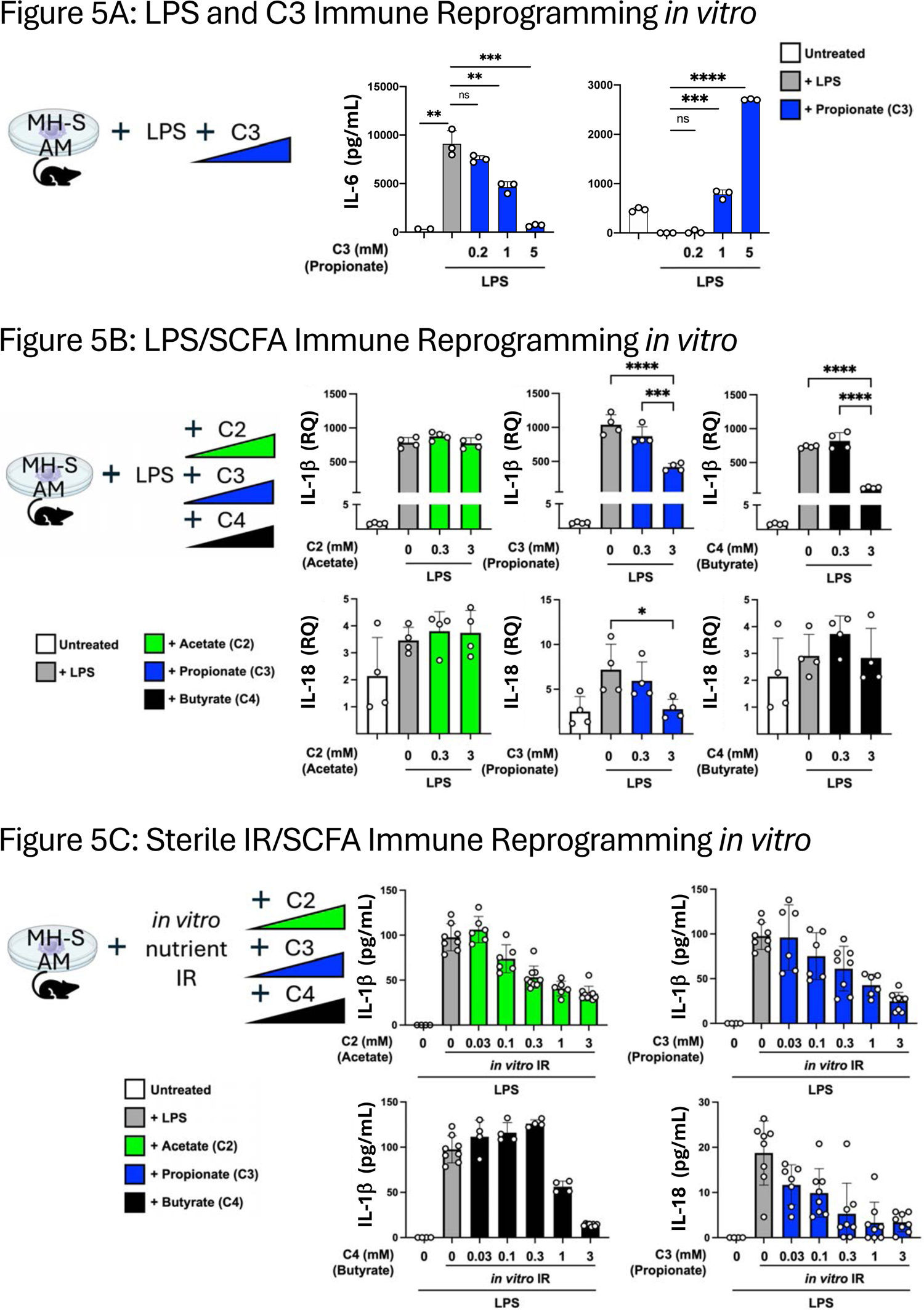
SCFAs can reprogram alveolar macrophage immune responses to LPS and IR *in vitro*. (A) Effect of increasing amounts of C3 (0.2, 1, and 5mM) on LPS (100ng/mL) inflammatory responses (added simultaneously) in MH-S alveolar macrophages *in vitro*. IL-6 and IL-10 levels were measured by ELISA at 24h (IL-6) and 48h (IL-10). (B) The alveolar macrophage cell line (MH-S) was pre-treated with different doses of C2, C3, and C4 (0, 0.3, and 3mM) for four hours and then challenged overnight with LPS (200ng/mL). IL-1β and IL-18 mRNA levels measured by qPCR. (C) The alveolar macrophage cell line (MH-S) was pre-treated with a range of C2, C3, and C4 concentrations (0, 0.03, 0.1, 0.3, 1, and 3mM) and then primed with LPS (200ng/mL) in the presence of the same type and dose of SCFA prior to *in vitro* IR. IL-1β and IL-18 levels measured after 1h reperfusion by ELISA. P values: *< 0.05; **< 0.01; ***< 0.001.

We next examined if all three SCFAs could affect the LPS-dependent induction of IL-1β and IL-18 mRNA similar to what we observed after lung injury *in vivo* with FR diets. We indeed observed that C3 (and C4) was able to blunt the induction of IL-1β and also prevent the increase of IL-18 (**Figure 5B**). Using an *in vitro* nutrient IR model, we also observed SCFAs altereing the inflammatory IR responses in AMs, with C2, C3 and C4 able to blunt the secretion of IL-1β and C3 able to do the same for IL-18 (**Figure 5C**).

### SCFAs reprogram AMs via metabolic shifts and nutrient intake versus chromatin accessibility changes and FFAR signaling

SCFAs are thought to act on immune and non-immune cells via three distinct (and potentially overlapping) mechanisms as shown in **Figure 6A**: 1) they can serve as an energy source and thereby affect cellular metabolism; 2) they can signal via GPCRs (with FFAR2 and FFAR3 being 2 primary SCFA receptors); and 3) they can act as HDAC inhibitors by modifying inflammatory and other gene loci. Furthermore, the actions of gut-derived SCFAs could act *directly* via metabolite transit to lung tissue, *indirectly* via actions on the bone marrow to alter the nature of cells that are trafficked at steady state to the lung, or both. To better understand the mechanistic basis for how SCFAs might affect AM immunometabolism and the shift from M1 to M2 phenotypes, we performed experiments to test the contribution of each of these 3 mechanisms and 3 routes of action. We also included this schematic in each of the following experiments to help confirm or eliminate the contribution of each of these mechanisms/routes.

**Figure 6.**
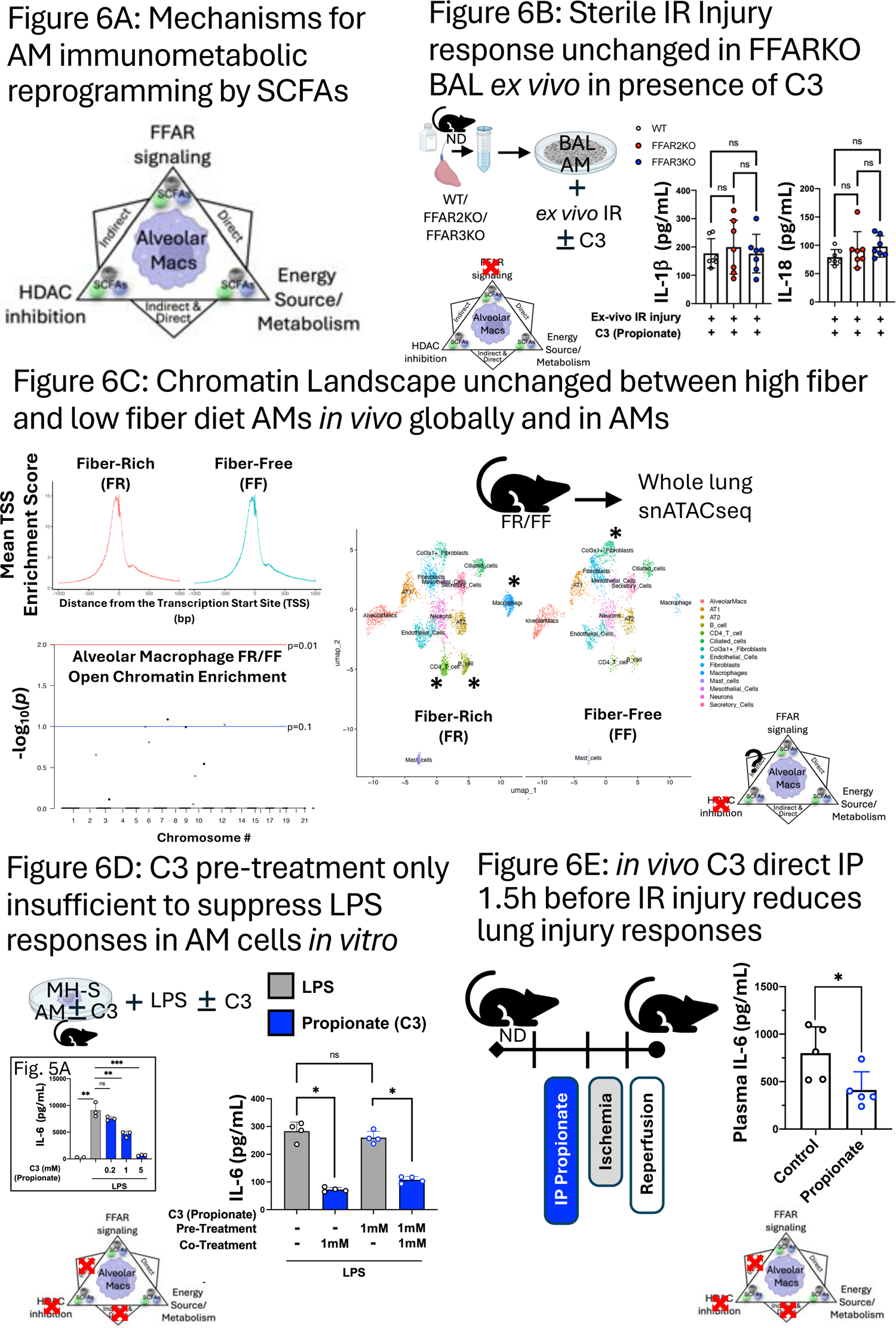
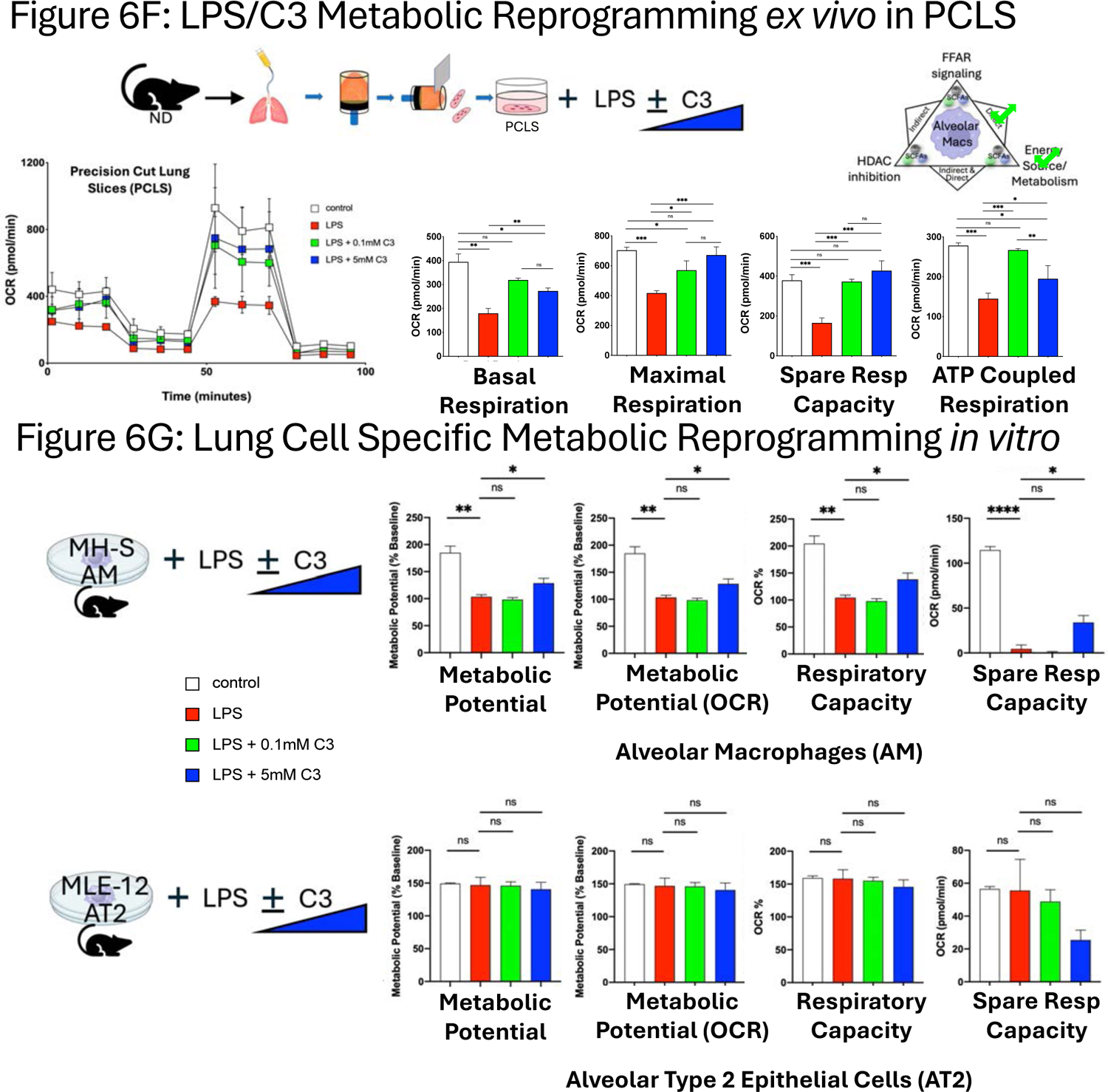
Propionate (C3) specifically alters AM immunometabolic programming independent of FFAR signaling and chromatin remodeling. _(A)_ Three possible mechanisms for SCFAs affecting AM immunometabolic programming. Alveolar macrophage image from Biorender^TM^. (B) WT, FFAR2 and FFAR3KO BAL AMs response to *ex vivo* nutrient IR after pre-treatment with 3mM C3. IL-1β and IL-18 levels measured 1h after *ex vivo* reperfusion by ELISA. (C) ATACseq of lung cell populations after 2 weeks of FR and FF diet. Transcription start site (TSS) enrichment comparison between the two groups (left top); AM population differences in chromatin accessible regions of the genome by chromosome number (left bottom). Blue line represents p=0.1, red line represents p=0.01. t-SNE comparison of calculated frequency of lung cell populations in FR and FF diet mice (right). Significantly different frequencies of cell clusters are denoted with an asterisk (*). (D) MH-S cells were pre-treated with C3 (1 mM) or medium 24h, followed by exposure to LPS (10 ng/mL) with or without co-treatment with C3 (1 mM) for 24h. IL-6 levels in the supernatant were measured by ELISA. (E) Propionate (10mg/kg) was administered IP 1.5h prior to onset of *in vivo* lung IR injury. Following IR injury, IL-6 levels in circulating plasma was measured by ELISA. (F) Oxygen consumption rate (OCR) as a measure of oxidative phosphorylation was performed on precision cut lung slices (PCLS) in the presence of 100ng/mL LPS and indicated C3 concentrations (0.1 and 5mM) for 18-24h assessed using the Seahorse Mito Stress Test. The line graph illustrates dynamic changes in OCR (left) and accompanying bar graphs present key metabolic parameters, including basal respiration, maximal respiration, spare respiratory capacity, and ATP-coupled respiration (right). (G) Seahorse metabolic measurements on alveolar macrophages (MH-S cell line, top graphs) alveolar type 2 cells (MLE-12 cell line, bottom graphs) treated *in vitro* with LPS (10ng/mL) and indicated C3 concentrations (0.1mM and 5mM) for 18-24h. Bar graphs (from left to right) detail the metabolic potential (as a percentage of baseline), metabolic potential OCR, respiratory capacity, and spare respiratory capacity for each cell type. P values are represented as follows in the figures: *< 0.05; **< 0.01; ***< 0.001; ****< 0.0001. All experiments were repeated at least twice and representative data shown.

**Figure 7.**
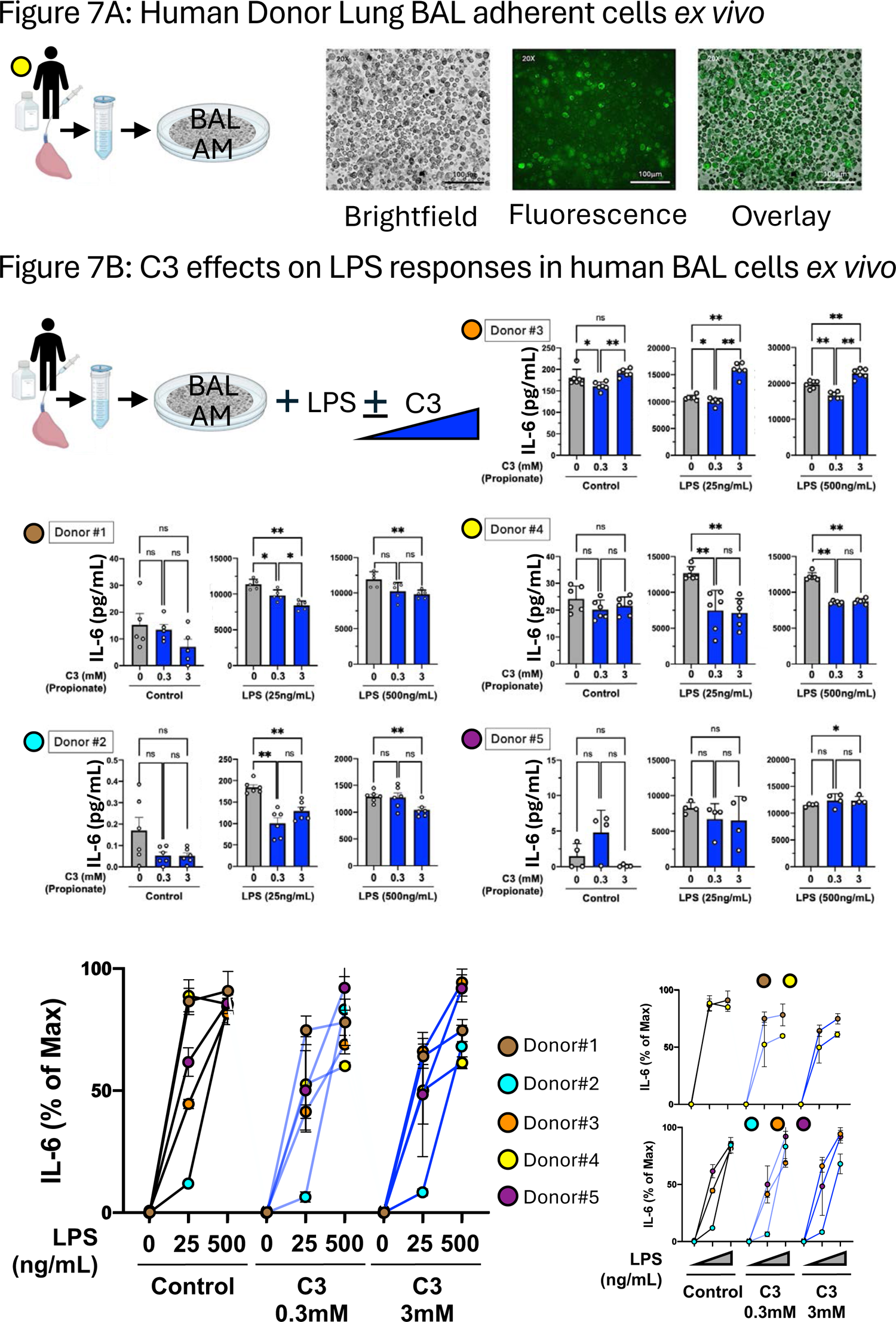
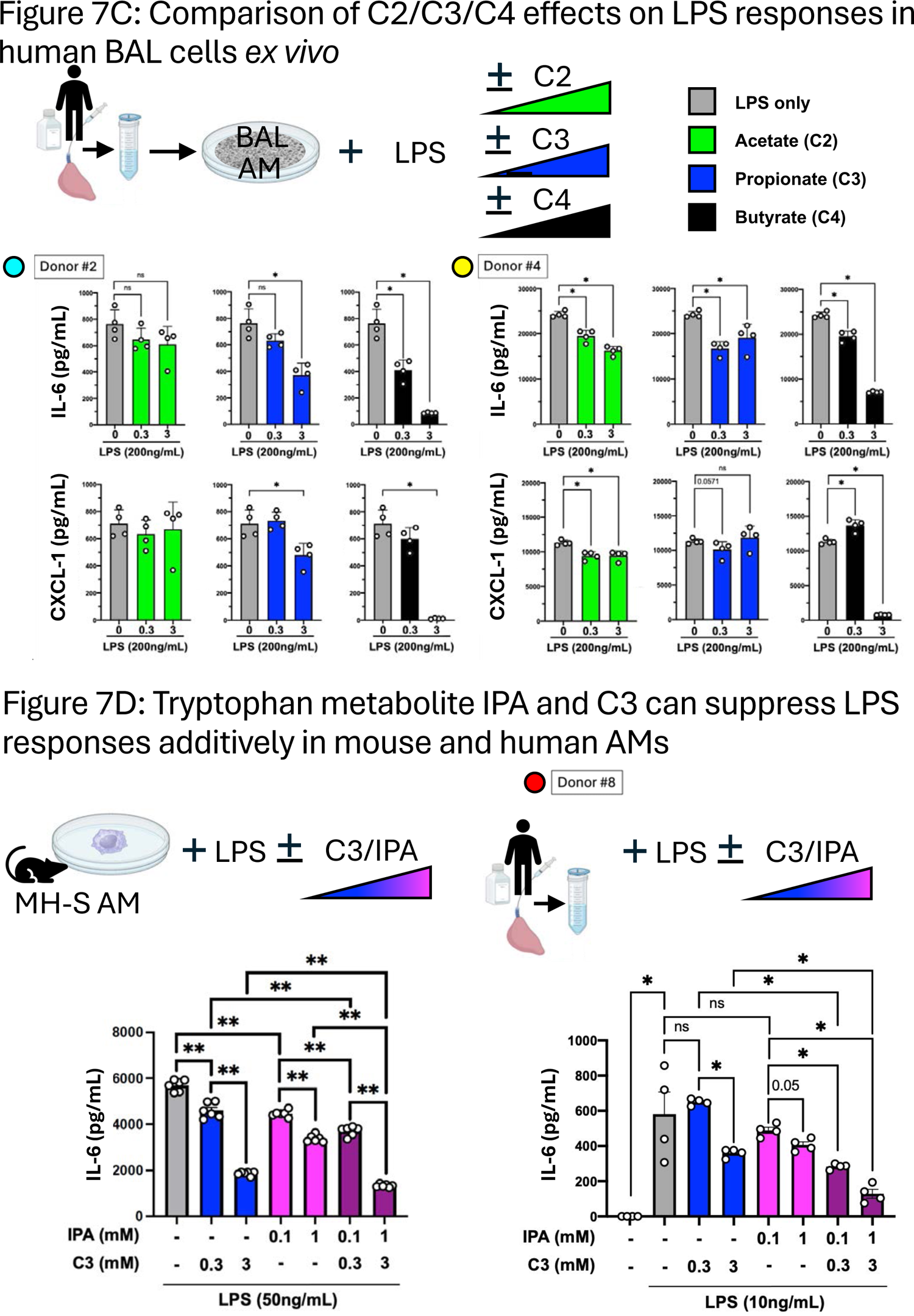
Gut Metabolites attenuates human BAL alveolar macrophage inflammatory responses to LPS *ex vivo*. (A, B) Human BAL cells (after plating for 2-12h) were imaged under brightfield and fluorescent microscopy to identify autofluorescent AMs. (B) Human BAL AMs from 5 human donor lungs were exposed to LPS (25ng/mL or 500ng/mL) in the presence of different concentrations of C3 (0mM, 0.3mM, and 3mM) overnight and IL-6 levels in supernatant measured by ELISA (top). Comparison of the responses for the set of 5 lung BAL AMs was performed by normalizing each to the % of maximum IL-6 production (bottom left). Human lung BAL AM responses (bottom right) from donor #s 1 and 4 (top) and donor #s 2, 3, and 5 (bottom) were observed as separating out into two distinct patterns. (C) Human BAL AMs were pretreated with C2, C3, or C4 (0mM, 0.3mM, and 3mM) and then exposed to LPS (200ng/mL) in the presence of the same type and dose of SCFA overnight. IL-6 and CXCL-1 levels in supernatant measured by ELISA. (D) MH-S cell line (left) and human BAL AMs (right) were pretreated with C3 (0.3 or 3 mM) and/or IPA (0.1 or 1 mM) or both at low C3+IPA or high C3+IPA concentrations as indicated, followed by overnight incubation with LPS (50 ng/mL or 10ng/mL as indicated) and the same levels of C3/IPA. IL-6 levels in the supernatant were measured by ELISA. P values: *< 0.05; **< 0.01; ***< 0.001.

First, we examined the effects of FFAR2 and FFAR3 by collecting BAL AMs from WT, FFAR2KO and FFAR3KO mice and subjecting them to *ex vivo* IR injury in the presence of propionate (C3). As shown in **Figure 6B**, the inability of AMs to signal through FFAR2 and FFAR3 had no effect on their secreted levels of IL-1β and IL-18 in the presence of high dose propionate (3mM C3), suggesting that the propionate-influenced inflammatory phenotype was not mediated by these receptors.

Next, we performed single nuclei (sn) ATACseq on whole lung tissue from mice (n=4 each) exposed to FR and FF diets for 2 weeks. We observed expected changes in the gut microbiota (**Figure S6**) but we failed to detect any major differences in transcriptional start site frequencies (TSS) (**Figure 6C, left top**) or any major genes of interest being in different open versus closed chromatin states between the FR and FF groups in all lung cell types (**Figure S7**) and specifically in AMs, our cell population of interest (**Figure 6C, left bottom**). We did however observe some differences in lung populations (**Figure 6C, right**) between the FR and FF groups, including the frequencies of non-AM macrophages (5x increase in FR), CD4^+^ T cells (3x increase in FR), B cells (3x increase in FR) and Col3a+ fibroblasts (3x decrease in FR) that reached statistical significance (**Table S4A**). Even among these cell populations there were no differences in the chromatin accessibility of the IL-1β and IL-18 loci and few if any significantly different chromatin accessibility genes or loci observed (**Tables S4B-F**). Overall, this result argued against HDACi being important via baseline alterations in lung chromatin accessibility but did not rule out the possibility that resident lung cellular population differences might account for the FR and FF lung injury responses we observed earlier.

### Pre-treatment with propionate (C3) alone did not attenuate LPS-induced inflammatory responses in alveolar macrophages in vitro

To determine when the presence of SCFAs during lung injury was essential for regulating pulmonary inflammatory responses, the MH-S AM cell line was pre-treated with C3 (0 or 1 mM) for 24h, then exposed to LPS (10 ng/ml) with or without co-treatment with C3 (1 mM) for an additional 24h. Co-treatment with C3 during LPS exposure significantly reduced IL-6 levels in the supernatants, as determined by ELISA, whereas pre-treatment with C3 alone did not produce this effect (**Figure 6D**) suggesting that C3 presence was required during the injury period and that this mechanism was unlikely to involve chromatin remodeling via HDACi.

### Rapid effect of systemic propionate on lung IR injury responses in vivo

To further test the ability of propionate to influence injury responses just by its presence during the injury period, we administered propionate IP immediately before (∼1.5h) creating lung IR injury *in vivo*. In this context, we were able to observe a clear and significant reduction in lung injury responses (**Figure 6E**). This result argues against SCFA and fiber diet reprogramming of bone marrow being required for the effects observed in attenuating lung injury. Similarly, it argues against changes in resident lung cell populations, as well as longer term reprogramming of resident lung immune cells being required for the ability of propionate to affect lung injury responses.

### Propionate metabolically reprograms lung tissue and AMs in the context of LPS injury

Our earlier published work showed that *in vitro* exposure of AM cell line (MH-S) to LPS skewed AMs towards glycolysis and this was reversed back to OX-PHOS by the presence of propionate (C3)^21^. To examine these effects in lung tissue *in situ*, we investigated whether C3 could affect the metabolism of lung tissue using *ex vivo* in precision cut lung slices (PCLS). LPS exposure in PCLS shifted metabolism from baseline OX-PHOS to favor glycolysis and the presence of C3 was able to reverse this back towards OX-PHOS (**Figure 6F**). C3 was able to partially or fully restore basal and maximal respiration, ATP-coupled respiration, and spare respiratory capacity in PCLS. To delve deeper into which cell type might be driving the PCLS metabolic responses, we compared these effects on AM vs. AT2 cells *in vitro*. We observed that AT2 cells (MLE-12 cell line) were much less metabolically active and were not significantly altered metabolically under the same conditions (**Figure 6G**). Overall, it appeared that the effects of C3 on lung tissue most closely mirrored those seen with isolated AMs.

### Human alveolar macrophages inflammatory responses in the presence of SCFAs

To extend our mouse lung findings to human alveolar macrophages and to study the effects of SCFAs on the cells that populate the airways in human lungs, we obtained BAL cells (attachment enriched AMs – confirmed by observation of cellular morphology and autofluorescence – **Figure 7A**) from human donor lungs and exposed them to lower dose and higher dose LPS injury alone or in the presence of propionate (0.3 and 3mM). We observed that similar to our murine data, C3 was able in general to attenuate the inflammatory responses of AMs and observed large degrees of variation in the magnitude of inflammation generated by low and high dose LPS as well as variable responses to propionate between 5 different donors (**Figure 7B, top**). To compare the responses among the 5 donor lung BAL AMs, we normalized them to the maximum levels of IL-6 produced and in doing so uncovered 2 distinct response patterns (**Figure 7B, bottom left**) – Donor #s 1 and 4 showed rapid responses to LPS with minimal dose responsiveness, yet demonstrated that C3 was able to suppress these responses. On the other hand, donor #s 2, 3, and 5 showed dose responsiveness to LPS and minimal response to C3 co-presence (**Figure 7B, bottom right**).

Next, we wanted to compare the effects of the three SCFAs. We observed that butyrate (C4) was most able to suppress human AM inflammatory responses the followed by propionate, as measured by IL-6 levels and CXCL-1 levels (**Figure 7C**) in the two donors tested (donor #s 2 and 4).

### Role of other gut metabolites in lung injury responses in mouse and human AMs

Other metabolites besides SCFAs are likely to also play important roles in lung injury. Given that some of the enriched taxa observed in our FR mice included those that have been reported to produce tryptophan metabolites, i.e. indoles (**Figure 2B, bottom right**; and confirmed in our metabolic analyses of FR vs FF portal blood vein levels, data not shown), we tested whether a key tryptophan metabolite, 3-indole propionic acid (IPA) could also modulate lung AM inflammatory response. Using the mouse AMs, we observed that IPA was able to also suppress LPS-mediated inflammation similar to propionate (C3) in a dose-dependent manner. Additionally, the presence of both metabolites during LPS injury increased the level of suppression (**Figure 7D, left**).

To confirm this observation in human AMs, we took BAL AMs from an independent donor lung (donor #8) and tested the effects of propionate and IPA (lower and higher doses) *ex vivo* on LPS-driven injury responses. Consistent with our *in vitro* mouse data, we observed that the combination of both low and higher doses of the two metabolites were able to suppress the levels of IL-6 produced than the single metabolites alone (**Figure 7D, right**).

## DISCUSSION

The main conclusions of the current study are that gut microbial metabolites reprogram alveolar macrophages downstream of dietary fiber intervention and alter inflammatory and host defense lung immune responses. These conclusions are based on the following experimental evidence. First, we tested the effects of a fiber-rich (FR) diet on alveolar macrophages (AMs) and lung injury responses and compared these effects against a fiber-free (FF) diet. Mice fed the FR diet exhibited distinct gut microbiota profiles and increased production of short-chain fatty acids (SCFAs) secondary to fiber fermentation. Notably, AMs from FR-fed mice showed transcriptional changes suggesting a shift toward an anti-inflammatory M2 phenotype with enriched transcription of genes involved in oxidative phosphorylation, suggestive of metabolic reprogramming. In terms of lung injury, mice on the FR diet experienced significantly reduced lung ischemia-reperfusion (IR) injury as evidenced by lower levels of inflammatory cytokines such as IL-6, IL-1β, and IL-18, particularly after two weeks on the diet. This reduction in cytokines was attributed to the modulatory effects of SCFAs produced from the FR diet, suggesting a protective role against lung injury.

Further multi-omic analyses, integrating 16S rRNA gene sequencing data, metabolomic and transcriptomic data, revealed correlations between SCFAs, gut microbiota profiles, and lung inflammatory markers, supporting the role of the gut-lung axis in regulating lung injury. Furthermore, fecal microbiota transplantation (FMT) from FR diet mice to germ-free mice replicated the reduced inflammatory response, underscoring the role of gut microbiota and propionate in mediating diet-induced changes in lung inflammation. However, extreme fiber diets impaired host defense against bacterial pneumonia, suggesting a complex balance between dietary fiber, gut microbiota, and immune responses in the lung.

Our previous work has shown that lung immunity can be influenced by the composition of the gut microbiome and that LPS and SCFAs can be transferred from the gut to the lung ^21,59^. Furthermore, our *in vitro* studies have shown that propionate can either prime or suppress alveolar macrophage inflammatory responses depending on the levels present and that enrichment of propionate-producing bacteria correlated with reduced lung IR inflammation ^27^. The importance of lung IR inflammation was also demonstrated by the fact that disruption of NLRP3 inflammasome-mediated IL-1β production or sensing was associated with worse bacterial clearance as part of a superimposed pneumonia^18^. Others have also demonstrated the importance of the gut microbiota in fighting lung infections ^60,61^ [and reviewed in ^62^], and high pectin fiber (as well as SCFAs directly) has been observed to alter lung adaptive immunity and bone remodeling ^9,24^.

Recent work has reported an alteration in type 2 inflammation following high-fiber diet with worse allergic lung responses and improved parasite clearance in mice^52^. This paper implicated ILC2s, IL-33, cholic acid, and the farnesoid X receptor (FXR) as driving the eosinophilia observed in their system. Interestingly, this group used inulin as their fiber source, but did not find significant elevations in fecal propionate and butyrate levels in their study. Other reports have shown that different fiber sources (inulin vs. pectin vs. psyllium) may have different effects on gut immunity with some fiber sources being beneficial and others detrimental depending on the injury model ^63^. We used pectin as the fiber source based on reports showing strong effects on lung adaptive immune responses and bone pathophysiology ^9,24^ and clearly observed increased SCFA levels in feces, portal blood and circulating plasma. Our work has also revealed some important differences in the regulation of IL-1β and IL-18, with only IL-1β being dependent on the presence of microbiota (also reported by others^64^), whereas both IL-1β and IL-18 in the lung appear to be modulated by the presence of acetate, propionate, heptanoate, and possibly other metabolites. Our focus on alveolar macrophages as the key cell regulating lung immune and metabolic tone is largely based on our previous work ^21^ but is also supported by work from other groups focusing on alveolar macrophages in IR lung injury ^65–67^, and gut microbiota programming of alveolar macrophages in the context of respiratory virus infection ^68^. Metabolic reprogramming of alveolar macrophages and epithelia has also been observed both locally by pathogens and through TLR4 engagement ^69,70^.

Our study begins to fill in important gaps in our understanding of how the gut microbiome influences lung injury responses through the gut-lung axis. Our findings that specific bacterial taxa can regulate the lung immunometabolic tone through the SCFAs they produce have important implications for human health, disease treatment, and prevention strategies. We can now begin to explore the effects of specific bacteria or consortia to alter lung injury responses downstream of metabolic reprogramming. Moreover, the ability of high-fiber diets to shape the gut microbiome, compared to FMT in the clinic, makes dietary interventions very attractive as a potentially powerful and simple tool to modulate lung injury responses, especially in the context of elective surgery. The direct administration of SCFAs to alter lung immunity and other organ responses has already been demonstrated by us and others^9,27^ and the broad therapeutic applicability of SCFAs and other gut metabolites needs to be further explored. Our data showing distinct responses of human lung BAL/AM responses to SCFAs between individuals tempts us to speculate that different levels of immunometabolic programming exists between individuals and that these might be driven by different dietary habits and microbiome compositions, but this theory requires further testing. Overall, compared to FMT and direct administration of SCFA (via pure compounds or foods rich in them), the use of dietary fiber to enrich for fermenting gut bacteria could be seen as more holistic, simpler, and perhaps preferable. Specific diets (not just based on fiber) could be envisioned to shape organ health, resilience, and immunity in both health, prior to injury, and in disease. However, a role for metabolites as treatment is also possible.

Reducing inflammatory responses after sterile lung IR injury may be beneficial in clinical scenarios such as organ transplantation, reperfusion after pulmonary embolism, or even hypovolemic trauma. However, much remains to be learned about the effects of a high-fiber diet and reduced lung immunometabolic tone on other common lung injuries and diseases, including pneumonia, asthma, and pulmonary fibrosis. Other groups have reported increased viral and bacterial killing by macrophages in the presence of specific SCFAs^71,72^. Our results suggest that extreme high-fiber diets dampen the ability of lung AMs and other cells to fight infection and subsequently cause worse lung damage. Our previous work has also shown that suppressing inflammation through the interruption of inflammatory signaling pathways also worsened pneumonia caused by *E.coli* and *S.aureus* lung infections^18^. While it is possible that strong suppression of inflammation is detrimental in the presence of an untreated infection, our *in vitro* data show that propionate does not indiscriminately suppress all immune responses, and perhaps at the appropriate balanced levels, may instead create an overall healthy state for alveolar macrophages and the lung as a whole.

As discussed earlier, a limitation of our study is the use of a single source of fiber (pectin). Additionally, other metabolites besides SCFAs could also be important such as lactate, primary and secondary bile acids, tryptophan derivatives (as we have also shown here), lipids, vitamins, etc. It is plausible that SCFAs could interact with any of the other classes of gut- and diet-derived metabolites and factors to influence lung immunometabolic tone. These interactions need to be investigated in a systematic and unbiased manner. Our future work will further investigate gut microbiome metabolic pathways enriched with high fiber diets and hopefully identify other metabolites that might be important in mediating the effects of dietary fiber as well as sampling the pulmonary artery blood for direct identification of the metabolites being delivered to the lung. Our work cannot formally rule out the importance of other non-AM cells in FR mouse lungs as well as the possibility however unlikely of rapid changes in chromatin accessibility occurring within 2h of lung injury initiation. Also precisely how lung resident cells utilize SCFAs as energy sources in the context of injury to direct their immunometabolic programming away from pro-inflammatory responses requires further detailed study.

The main conclusions of this study are that a high-fiber diet leads to the reprogramming of alveolar macrophages towards an anti-inflammatory state and reduces sterile lung injury via the production of short-chain fatty acids, although it may impair defense against bacterial infections, highlighting the complex interplay between diet, gut microbiota, and lung health. Specifically for the field of sterile lung injury, these findings have broad implications for the use of dietary interventions prior to general surgery, lung transplantation, and other anticipated procedures involving lung injury. Further understanding of specific bacteria that interact with and manipulate lung immunometabolic tone may allow the identification of resilient patient populations versus at-risk populations that may benefit from targeted dietary interventions (prebiotics), FMT with lung-protective taxa or combined synbiotic approaches. The use of specific metabolites, such as propionate and butyrate, to directly manipulate this system may also have a place in emergency situations where short-term manipulation to enhance or suppress lung immune responses may be in the best interest of the patient.

Overall, the use of dietary interventions and specific metabolites to modulate lung function and immunometabolic tone are likely to lead to novel therapeutic approaches that harness and leverage the gut microbiome’s ability to regulate lung inflammation.

## ACKNOWLEDGMENTS

We would like to acknowledge the following individuals for assistance with reagents, mice, advice, helpful discussions, and critical reading and editing of the manuscript: Susan Lynch (UCSF), Richard Locksley (UCSF), Mark Ansel (UCSF), Itsaso Garcia-Arcos (SUNY-Downstate Health Sciences University), Martin Valdearcos (UCSF), Judith Hellman (UCSF), Aditi Bhargava and Johannes Knapp (Aseesa, Inc.), Mervyn Maze (UCSF), Elisabetta Pusceddu (ARNAS G. Brotzu, Anesthesia and Intensive Care Unit, Liver Transplantation Center, Cagliari, Italy). Thank you to Christine Tat for assisting with some ELISA experiments. We also acknowledge the contributions of the Metabolomics Core at University of Michigan (specifically Kari Bonds and Maureen Kachmann for SCFA analyses); the BCMM (Abdur Rahim Khan), Genomics CoLab (Walter Eckalbar), and NORC (Gnotobiotic) core facilities (Jesse Turnbaugh) at UCSF. WT, control and mutant *B.theta* strains were generously provided by Eric Martens (University of Michigan). The authors would also like to acknowledge the Donor Network West for providing lungs for research to MAM; and the continuing support of THFC (COYS) on these studies.

## Abbreviations used in manuscript

SCFA: short-chain fatty acid
MCFA: medium-chain fatty acid
BCFA: branch-chain fatty acid
C2: acetate
C3: propionate
C4: butyrate
iC4: isobutyrate
2-me-C4: 2-methyl butyrate
C5: valerate
iC5: isovalerate
C6: caproate
C7: heptanoate
AM: Alveolar Macrophage
BAL: Bronchoalveolar Lavage
*B.theta*: Bacteriodes thetaiotomicron
FMT: Fecal Microbiota Transplantation
IPA: 3-indole propionic acid (aka indole propionic acid)
IR: ischemia reperfusion
FR: fiber-rich
FF: fiber-free
GF: germ-free
SPF: specific pathogen free
OX-PHOS: Oxidative Phosphorylation
PCLS: Precision Cut Lung Slices

