## Supplemental Figures for "Gut Microbiome-derived Propionate Reprograms Alveolar Macrophages Metabolically and Regulates Lung Injury Responses"

**Supplemental Figure S1:** A. Composition of diets used in study (fiber levels highlighted in bold). B. Representative image of stool and colon morphology in FF vs FR groups. C. weight changes over diet course. B. Normal chow diet composition.

### Figure S1A

**Composition of diets used in this study**

| Product # | Normal diet | | Fiber free (FF) diet | | Fiber rich (FR) diet | |
|---|---|---|---|---|---|---|
| | PicoLab Rodent Diet 20 (5058) | | Modified AIN-93G/No Cellulose(5GCX) | | Mod TestDiet® 57W5 w/ 30% Pectin, Blue (5Z6R) | |
| | gm% | kcal% | gm% | kcal% | gm% | kcal% |
| **Protein** | 21.8 | 23.217 | 18.3 | 17.900 | 18.0 | 26.900 |
| **Fat (ether extract)** | 9.0 | 21.566 | 7.1 | 15.600 | 7.0 | 23.600 |
| **Carbohydrates** | 51.8 | 55.217 | 68.2 | 66.600 | 33.2 | 49.600 |
| **Total fiber** | **2.4** | 0 | **0.0** | 0 | **35.0** | 0 |
| **Kcal/gm** | | 100 | | 100 | | 100 |
| **Energy(kcal/g)** | 4.63 | | 4.10 | | 2.68 | |

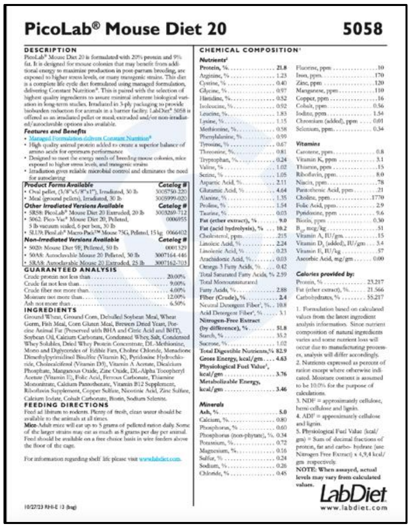
PicoLab® Mouse Diet 20 5058

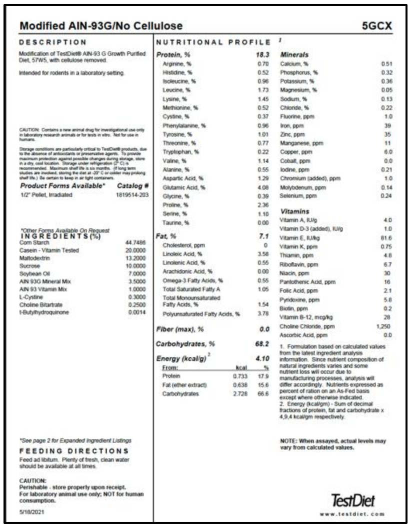
Modified AIN-93G/No Cellulose 5GCX

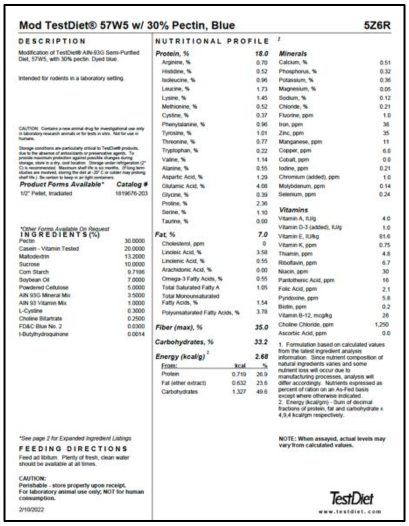
Mod TestDiet® 57W5 w/ 30% Pectin, Blue 5Z6R

### Figure S1B

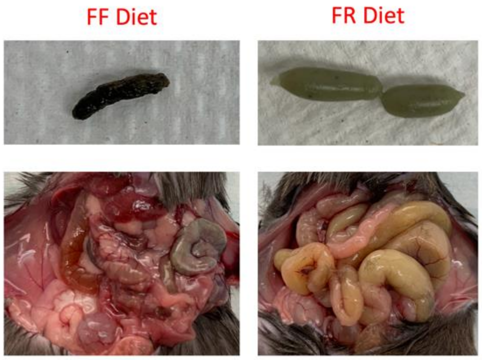


### Figure S1C

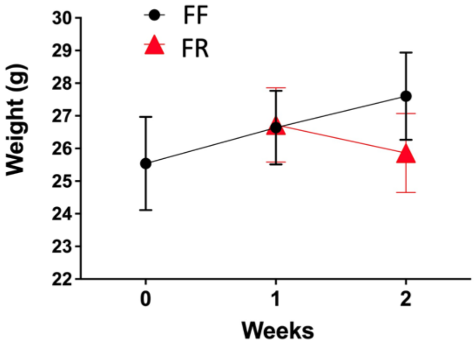

**Supplemental Figure S2: No difference in Gut leakiness between FR and FF mice.** The LBP concentrations in systemic plasma collected from mice subjected to fiber-rich diet (FR), or fiber-free diet (FF).

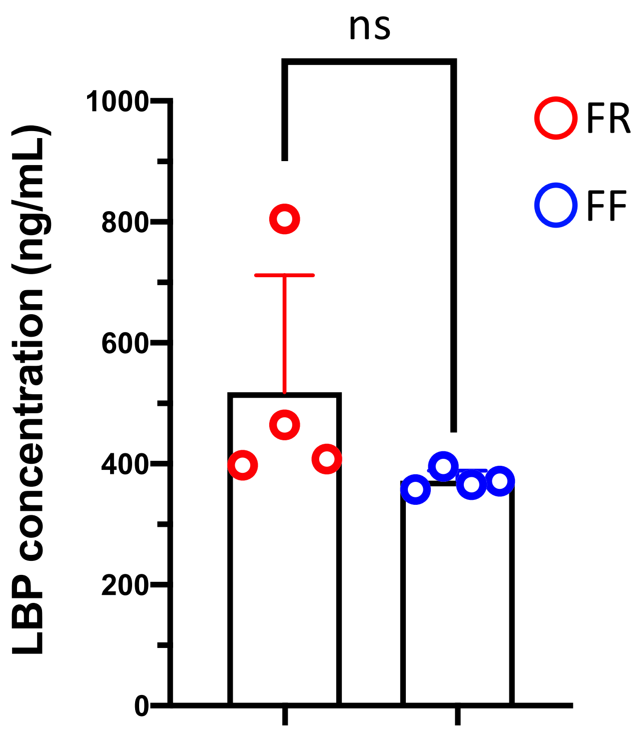

**Supplemental Table S1: Gene-set Enrichment Analyses (GSEA) for FR/FF diet mice BAL bulk RNAseq (pre-injury).** OX-PHOS genes enriched are listed for the MSigDB and KEGG databases (above two boxes). All gene sets enriched ($p<0.05$) listed in table below with the two OX-PHOS entries highlighted in yellow.

**MSigDB: OX-PHOS genes enriched in FR/FF diet mice BAL bulk RNAseq:**
COX7B;ACADVL;ACAA2;COX4I1;ETFA;PHB2;TOMM22;NNT;UQCRFS1;ACADM;DLAT;IDH3A;BCKDHA;CPT1A;ECH1;SDHC;SDHD;ATP1B1;COX6B1;POR;NDUFS8;UQCRC1;SUCLG1;VDAC1;SLC25A5;SLC25A4;NDUFB8;ECHS1;RETSAT;COX15;NDUFB5;TIMM13;ETFDH;DLST;UQCR11;ACAT1;PRDX3;LDHA;CYB5R3;POLR2F;SLC25A20;NDUFV2;NDUFV1;ATP6V1C1;NDUFA9;NDUFA5;MDH2;NDUFA4;IDH3G;GOT2;NDUFA1;IMMT;COX6C;CS;GLUD1;ALDH6A1;UQCRQ

**KEGG: OX-PHOS genes genes enriched in FR/FF diet mice BAL bulk RNAseq:**
NDUFA13;COX7B;NDUFB8;NDUFA11;COX15;NDUFB5;COX4I1;ATP5A1;ATP5K;ATP5C1;ATP5J;UQCR11;ATP5H;ATP5G3;ATP5O;ATP5G1;ATP5B;ATP5E;UQCRFS1;NDUFV3;NDUFV2;NDUFV1;ATP6V1C1;LHPP;NDUFA9;NDUFA5;NDUFA4;NDUFA1;SDHC;SDHD;COX6C;COX6B1;NDUFS8;UQCRQ;UQCRC1;ATP6V0D2

**GSEA (all three databases – KEGG, MSigDB, GO): adjusted p<0.05 pathways enriched in FR/FF diet mice BAL bulk RNAseq – OX-PHOS pathways highlighted in yellow**

| Enrichment Category | Term | Overlap | P value | Adjusted P value | Odds Ratio | Combined Score | Genes |
|---|---|---|---|---|---|---|---|
| GO_Biological_Process_2021 | cotranslational protein targeting to membrane (GO:0006613) | 40/94 | 1.4 790619 756545 3E -13 | 6.9 146147 361849 1E-10 | 5.4 030864 197 530 9 | 159 .61904 948434 2 | RPL5;RPL30;TRAM1;RPL3;RPL32;RPL10;RPL34;RPL12;RPL36A;RPL10A;RPL8;RPS4X;RPL7A;SEC61A1;RPS16;RPS15A;RPS19;SRP72;RPS18;RPL36;RPL14;RPS3;RPLP2;SSR1;RPL15;RPS27A;SIL1;RPL17;RPS13;RPS12;SEC63;RPL19;RPL21;SSR2;RPS6;RPL35A;RPSA;RPS25;RPL27;RPL29 |
| GO_Biological_Process_2021 | SRP-dependent cotranslational protein targeting to membrane (GO:0006614) | 37/90 | 4.2 435955 600739 4E -12 | 9.9 194046 216728 3E-09 | 5.0 860873 593 542 7 | 133 .18230 101785 5 | RPL5;RPL30;RPL3;RPL32;RPL10;RPL34;RPL12;RPL36A;RPL10A;RPL8;RPS4X;RPL7A;SEC61A1;RPS16;RPS15A;RPS19;SRP72;RPS18;RPL36;RPL14;RPS3;RPLP2;SEC61B;RPL15;RPS27A;RPL17;RPS13;RPS12;SEC63;RPL19;RPL21;RPS6;RPL35A;RPSA;RPS25;RPL27;RPL29 |
| MSigDB_Hallmark_2020 | Oxidative Phosphorylation | 57/200 | 4.1 390814 848022 8E -10 | 2.0 695407 424011 4E-08 | 2.9 133195 801 261 3 | 62. 943367 938359 6 | COX7B;ACADVL;ACAA2;COX4I1;ETFA;PHB2;TOMM22;NNT;UQCRFS1;ACADM;DLAT;IDH3A;BCKDHA;CPT1A;ECH1;SDHC;SDHD;ATP1B1;COX6B1;POR;NDUFS8;UQCRC1;SUCLG1;VDAC1;SLC25A5;SLC25A4;NDUFB8;ECHS1;RETSAT;COX15;NDUFB5;TIMM13;ETFDH;DLST;UQCR11;ACAT1;PRDX3;LDHA;CYB5R3;POLR2F;SLC25A20;NDUFV2;NDUFV1;ATP6V1C1;NDUFA9;NDUFA5;MDH2;NDUFA4;IDH3G;GOT2;NDUFA1;IMMT;COX6C;CS;GLUD1;ALDH6A1;UQCRQ |
| KEGG_2019_Mouse | Huntington disease | 57/192 | 7.2 418830 638037 E-11 | 2.1 725649 191411 1E-08 | 3.0 873781 880 915 7 | 72. 085818 685377 2 | NDUFA13;COX7B;NDUFA11;COX4I1;ATP5C1;ATP5G3;AP2A2;ATP5G1;DNALI1;CREB3L1;UQCRFS1;AP2M1;GPX1;SDHC;SDHD;COX6B1;CREB3;NDUFS8;PLCB4;UQCRC1;VDAC1;SLC25A5;SLC25A4;PLCB2;CREB5;DNAH3;NDUFB8;DNAH1;DCTN2;NDUFB5;DNAH5;ATP5A1;DNAH6;ITPR1;ATP5J;UQCR11;ATP5H;ATP5O;ATP5B;ATP5E;POLR2D;AP2S1;POLR2F;NDUFV3;NDUFV2;POLR2I;NDUFV1;NDUFA9;CREBBP;DNAH11;DNAH10;NDUFA5;NDUFA4;NDUFA1;COX6C;UQCRQ;DNAL1 |
| GO_Biological_Process_2021 | protein targeting to ER (GO:0045047) | 39/103 | 2.3 572307 570716 8E -11 | 3.6 733512 631033 7E-08 | 4.4 404935 443 565 2 | 108 .66308 884252 6 | RPL5;RPL30;RPL3;RPL32;RPL10;RPL34;RPL12;RPL36A;RPL10A;RPL8;RPS4X;RPL7A;SEC61A1;RPS16;RPS15A;RPS19;SRP72;RPS18;RPL36;RPL14;RPS3;RPLP2;RPL15;RPS27A;RPL17;RPS13;RPS12;SEC63;RPL19;RPL21;RPS6;RPL35A;RPSA;RPS25;SPCS2;SPCS1;CHMP4B;RPL27;RPL29 |
| MSigDB_Hallmark_2020 | Adipogenesis | 53/200 | 2.6 234451 408989 6E -08 | 6.5 586128 522473 9E-07 | 2.6 301451 456 158 5 | 45. 912319 572078 8 | COX7B;SLC27A1;ACAA2;SCP2;ALDH2;ME1;ACADM;DLAT;ACADS;IDH3A;BCKDHA;ECH1;ELOVL6;ATP1B3;SDHC;POR;TST;COL4A1;ACOX1;GPD2;CAT;UQCRC1;CMPK1;SUCLG1;CD302;ECHS1;CD151;RETSAT;AK2;UQCR11;SLC1A5;LTC4S;ADIPOR2;TOB1;C3;PRDX3;SAMM50;DHRS7B;SLC19A1;ABCA1;ESRRA;NDUFA5;MDH2;IDH3G;IMMT;CS;QDPR;DHRS7;PFKL;REEP5;BCL6;UQCRQ;LPCAT3 |
| GO_Biological_Process_2021 | cytoplasmic translation (GO:0002181) | 34/93 | 1.3 078226 185122 5E -09 | 1.5 285176 853862 E-06 | 4.1 917381 687 025 4 | 85. 741594 313405 7 | EIF4A1;RPL5;RPL30;RPL3;RPL32;RPL10;RPL34;RPL12;RPL36A;RPL10A;RPL8;RPS4X;RPL7A;RPS16;RPS15A;RPS19;RPS18;RPL36;RPL14;RPS3;RPL15;RPS27A;RPL17;RPS13;RPS12;RPL19;RPL21;RPS6;RPL35A;RPSA;RPS25;EIF3I;RPL27;RPL29 |
| GO_Biological_Process_2021 | nuclear-transcribed mRNA catabolic process, nonsense-mediated decay (GO:0000184) | 38/113 | 2.2 564527 139732 2E -09 | 2.1 097832 875649 6E-06 | 3.6 882042 742 159 3 | 73. 430199 183237 2 | RPL5;RPL30;SMG1;RPL3;RPL32;RPL10;RPL34;RPL12;EIF4A3;RPL36A;RPL10A;RPL8;RPS4X;RPL7A;RPS16;RPS15A;RPS19;PPP2R1A;RPS18;MAGOH;RPL36;RPL14;RPS3;RPLP2;RPL15;RPS27A;RPL17;RPS13;RPS12;RPL19;RPL21;RPS6;PYM1;RPL35A;RPSA;RPS25;RPL27;RPL29 |
| KEGG_2019_Mouse | Alzheimer disease | 46/175 | 2.7 519439 735386 5E -07 | 4.1 279159 603079 7E-05 | 2.5 963681 814 355 | 39. 220187 551207 2 | APP;NDUFA13;COX7B;NDUFB8;NDUFA11;NDUFB5;COX4I1;ATP5A1;ITPR1;ATP5C1;ATP5J;UQCR11;CALML4;ATP5H;ATP5G3;ATP5O;ATP5G1;APH1A;ATP5B;PPP3CA;NCSTN;PPP3R1;APH1C;ATP5E;UQCRFS1;NDUFV3;NDUFV2;NDUFV1;NDUFA9;NDUFA5;BAD;NDUFA4;NDUFA1;SDHC;SDHD;COX6C;COX6B1;NDUFS8;PLCB4;UQCRQ;UQCRC1;CALM3;CALM1;CALM2;PLCB2;GAPDH |
| GO_Biological_Process_2021 | endoplasmic reticulum to Golgi vesicle-mediated transport (GO:0006888) | 48/185 | 2.3 375232 593349 3E -07 | 0.0 001821 320206 231 8 | 2.5 520103 508 043 8 | 38. 966655 535596 5 | DCTN6;PGAP1;TMED10;ATL3;DCTN2;SAR1B;PDCD6;PROS1;GOSR2;RAB1B;DCTN3;TEX261;IER3IP1;USE1;LMAN1;ACTR1A;CAPZB;TMED3;BCAP29;TMED2;MED1;TMED7;YKT6;CTSC;TMED5;CD55;TMED4;TRAPPC3;SEC13;TMED9;F10;TRAPPC5;DYNLL2;COPZ1;CNIH1;VAMP7;TBC1D20;GOLGB1;GORASP1;COL7A1;CAPZA2;COPG1;ERGIC3;GAS6;ACTR10;CNIH4;COPE;BET1 |
| KEGG_2019_Mouse | Parkinson disease | 38/144 | 2.6 152463 927148 2E -06 | 0.0 002273 414591 895 12 | 2.6 049518 483 025 2 | 33. 484447 641291 6 | NDUFA13;COX7B;NDUFB8;NDUFA11;NDUFB5;LRRK2;COX4I1;ATP5A1;ATP5C1;ATP5J;UQCR11;ATP5H;ATP5G3;ATP5O;ATP5G1;UBE2J1;GNAI2;ATP5B;ATP5E;UQCRFS1;NDUFV3;PRKACA;NDUFV2;NDUFV1;NDUFA9;NDUFA5;NDUFA4;NDUFA1;SDHC;SDHD;COX6C;COX6B1;NDUFS8;UQCRQ;UQCRC1;VDAC1;SLC25A5;SLC25A4 |
| KEGG_2019_Mouse | Oxidative phosphorylation | 36/134 | 3.0 312194 558601 6E -06 | 0.0 002273 414591 895 12 | 2.6 683079 221 705 3 | 33. 904976 177776 4 | NDUFA13;COX7B;NDUFB8;NDUFA11;COX15;NDUFB5;COX4I1;ATP5A1;ATP5K;ATP5C1;ATP5J;UQCR11;ATP5H;ATP5G3;ATP5O;ATP5G1;ATP5B;ATP5E;UQCRFS1;NDUFV3;NDUFV2;NDUFV1;ATP6V1C1;LHPP;NDUFA9;NDUFA5;NDUFA4;NDUFA1;SDHC;SDHD;COX6C;COX6B1;NDUFS8;UQCRQ;UQCRC1;ATP6V0D2 |
| KEGG_2019_Mouse | Focal adhesion | 47/199 | 5.3 486576 473304 4E -06 | 0.0 003209 194588 398 26 | 2.2 493676 732 575 3 | 27. 304320 502597 9 | ITGB1;ROCK1;SHC1;ITGB4;ITGA2B;PIK3R3;ILK;ACTB;MYL12A;EGFR;THBS3;PPP1CB;RAP1B;CDC42;RAP1A;CCND1;AKT2;CHAD;RAC2;AKT1;RAC3;FLNA;FLNB;PIP5K1C;RAC1;VASP;VAV3;PPP1R12A;VWF;LAMB3;PRKCB;CAV2;BAD;LAMB2;CAV1;FN1;PRKCA;BRAF;PARVA;IGF1;RHOA;PPP1CA;PARVG;COL4A1;ITGA10;RAF1;MET |
| MSigDB_Hallmark_2020 | Fatty Acid Metabolism | 38/158 | 2.6 584855 850025 E-05 | 0.0 004430 809308 337 5 | 2.2 991951 151 817 9 | 24. 222408 719029 2 | GCDH;ACADVL;ACAA2;ECHS1;RETSAT;MAOA;ECI2;ETFDH;DLST;HSD17B11;LTC4S;ADIPOR2;LDHA;UROD;ME1;ACADM;ACSS1;ACADS;S100A10;CBR1;CPT1A;HSDL2;MDH2;IDH3G;ECH1;FMO1;SDHC;SDHD;ALDH3A2;ACOX1;GPD2;FASN;EHHADH;APEX1;GPD1;ALDH1A1;SUCLG1;SERINC1 |
| MSigDB_Hallmark_2020 | PI3K/AKT/mTOR Signaling | 28/105 | 4.2 258629 305675 E-05 | 0.0 005282 328663 209 37 | 2.6 357605 909 844 7 | 26. 546595 165909 7 | ARF1;RALB;PRKAA2;YWHAB;PRKAG1;SLA;PIK3R3;NOD1;IL2RG;EGFR;PPP2R1B;ARHGDIA;PRKAR2A;CFL1;AKT1;RAC1;AP2M1;ACTR3;VAV3;PLA2G12A;PRKCB;STAT2;PPP1CA;PIKFYVE;PIN1;RAF1;SQSTM1;MYD88 |
| GO_Biological_Process_2021 | nuclear-transcribed mRNA catabolic process (GO:0000956) | 44/171 | 9.3 531663 306347 2E -07 | 0.0 006246 578942 245 33 | 2.5 207760 966 440 1 | 34. 994373 483481 | RPL5;RPL30;SMG1;RPL3;RPL32;RPL10;RPL34;RPL12;EIF4A3;RPL36A;RPL10A;RPL8;RPS4X;RPL7A;RPS16;RPS15A;RPS19;PPP2R1A;RPS18;MAGOH;EXOSC8;POLR2D;RPL36;RPL14;RPS3;RPLP2;RPL15;RPS27A;RPL17;RPS13;RPS12;RPL19;RPL21;RPS6;PYM1;RPL35A;RPSA;CNOT4;RPS25;PNLDC1;DXO;RPL27;RPL29;CNOT9 |
| GO_Biological_Process_2021 | neutrophil degranulation (GO:0043312) | 95/481 | 1.1 650640 834869 8E -06 | 0.0 006391 244739 395 35 | 1.8 024569 970 281 8 | 24. 626491 335291 9 | B4GALT1;ATP8A1;HEXB;RAB3D;GDI2;AP2A2;CTSS;PYCARD;LGALS3;PSMD6;ACTR1B;GM2A;ANPEP;FTH1;STBD1;NAPRT;ITGAX;COTL1;RAC1;CEP290;CTSC;DSP;CAP1;CD53;MANBA;FCER1G;ANXA2;ARG1;SLC11A1;MCEMP1;CYBA;TMC6;KCNAB2;DDOST;RHOA;VAMP8;RAP2C;FGR;VAMP7;CLEC4D;RAB31;DOK3;SLPI;PSMA2;TMBIM1;CAT;ADAM8;CD47;CD44;ASAH1;RAB5C;TMEM63A;ROCK1;CLEC12A;PSMD13;STXBP2;UBR4;MGST1;PRCP;FPR2;CFP;AGPAT2;C3;RAP1B;TRPM2;NCSTN;NRAS;PGRMC1;CYB5R3;RAP1A;CXCR1;ALOX5;ALDH3B1;PSMB1;CXCR2;ORMDL3;CD55;TXNDC5;XRCC6;TMEM30A;FUCA1;ARPC5;PFKL;TSPAN14;LRG1;FABP5;IMPDH1;SLCO4C1;RAB18;LAMTOR2;MAN2B1;LAMTOR3;ACTR10;HSPA1B;HSPA1A |
| GO_Biological_Process_2021 | neutrophil mediated immunity (GO:0002446) | 96/488 | 1.2 304000 567819 9E -06 | 0.0 006391 244739 395 35 | 1.7 936894 894 476 6 | 24. 408833 637777 | B4GALT1;ATP8A1;HEXB;RAB3D;GDI2;AP2A2;CTSS;PYCARD;LGALS3;PSMD6;ACTR1B;GM2A;ANPEP;FTH1;STBD1;NAPRT;ITGAX;COTL1;RAC1;CEP290;CTSC;DSP;CAP1;CD53;MANBA;FCER1G;ANXA2;ARG1;SLC11A1;MCEMP1;CYBA;TMC6;KCNAB2;DDOST;RHOA;VAMP8;RAP2C;FGR;VAMP7;CLEC4D;RAB31;DOK3;SLPI;PSMA2;TMBIM1;CAT;ADAM8;CD47;CD44;ASAH1;RAB5C;TMEM63A;ROCK1;CLEC12A;PSMD13;STXBP2;UBR4;MGST1;PRCP;FPR2;CFP;AGPAT2;C3;RAP1B;TRPM2;NCSTN;NRAS;PGRMC1;CYB5R3;RAP1A;CXCR1;ALOX5;ALDH3B1;PSMB1;CXCR2;ORMDL3;CD55;TXNDC5;XRCC6;ACE;TMEM30A;FUCA1;ARPC5;PFKL;TSPAN14;LRG1;FABP5;IMPDH1;SLCO4C1;RAB18;LAMTOR2;MAN2B1;LAMTOR3;ACTR10;HSPA1B;HSPA1A |
| GO_Biological_Process_2021 | neutrophil activation involved in immune response (GO:0002283) | 95/485 | 1.6 955680 044506 4E -06 | 0.0 007926 780420 806 74 | 1.7 835547 537 040 1 | 23. 698970 890098 | B4GALT1;ATP8A1;HEXB;RAB3D;GDI2;AP2A2;CTSS;PYCARD;LGALS3;PSMD6;ACTR1B;GM2A;ANPEP;FTH1;STBD1;NAPRT;ITGAX;COTL1;RAC1;CEP290;CTSC;DSP;CAP1;CD53;MANBA;FCER1G;ANXA2;ARG1;SLC11A1;MCEMP1;CYBA;TMC6;KCNAB2;DDOST;RHOA;VAMP8;RAP2C;FGR;VAMP7;CLEC4D;RAB31;DOK3;SLPI;PSMA2;TMBIM1;CAT;ADAM8;CD47;CD44;ASAH1;RAB5C;TMEM63A;ROCK1;CLEC12A;PSMD13;STXBP2;UBR4;MGST1;PRCP;FPR2;CFP;AGPAT2;C3;RAP1B;TRPM2;NCSTN;NRAS;PGRMC1;CYB5R3;RAP1A;CXCR1;ALOX5;ALDH3B1;PSMB1;CXCR2;ORMDL3;CD55;TXNDC5;XRCC6;TMEM30A;FUCA1;ARPC5;PFKL;TSPAN14;LRG1;FABP5;IMPDH1;SLCO4C1;RAB18;LAMTOR2;MAN2B1;LAMTOR3;ACTR10;HSPA1B;HSPA1A |
| KEGG_2019_Mouse | Ribosome | 40/170 | 2.8 457711 632526 2E -05 | 0.0 012248 739146 206 4 | 2.2 346153 846 153 8 | 23. 389923 418709 9 | RPL5;RPL30;RPL3;RPL32;RPL10;MRPS14;RPL34;RPL12;MRPL16;RPL36A;MRPL17;RPL8;RPL10A;RPS4X;MRPL20;RPL7A;MRPL2;RPS16;RPS15A;RPS19;RPS18;RPL36;RPL14;RPS3;RPLP2;RPL15;RPS27A;RPL17;RPS13;RPS12;RPL19;RPL21;RPS6;RPL35A;RPSA;RPS25;MT-RNR2;MT-RNR1;RPL27;RPL29 |
| KEGG_2019_Mouse | Long-term potentiation | 21/67 | 2.8 580391 341148 3E -05 | 0.0 012248 739146 206 4 | 3.3 053004 295 702 5 | 34. 582663 265863 7 | CREBBP;PRKCB;ITPR1;PRKCA;BRAF;CALML4;PPP1CA;PPP1CB;RAP1B;PPP3CA;NRAS;PPP3R1;PLCB4;RAP1A;CALM3;PRKACA;RAF1;CALM1;PLCB2;CALM2;ATF4 |
| KEGG_2019_Mouse | Endocytosis | 56/269 | 3.8 907029 381980 4E -05 | 0.0 014590 136018 242 6 | 1.9 130510 130 132 | 19. 425762 043853 9 | ARF3;WASHC4;H2-T24;VPS29;ARF1;ZFYVE9;ARPC1B;CBLB;ARRB1;AP2A2;CAPZB;CHMP1B;CHMP1A;PSD3;AP2M1;SH3GLB2;RHOA;EPN1;RAB31;PARD3;RAB35;CHMP4B;CHMP3;CHMP6;ARF5;RAB5C;H2-M2;H2-Q6;H2-Q7;WAS;H2-Q4;NEDD4L;IL2RG;EGFR;CDC42;SNX1;GRK3;CXCR1;CXCR2;GRK6;AP2S1;PIP5K1C;LDLR;CAV2;CAV1;ARPC4;SNF8;ARPC5;PML;RAB11B;NEDD4;CAPZA2;CHMP2A;FGFR3;HSPA1B;HSPA1A |
| GO_Biological_Process_2021 | aerobic electron transport chain (GO:0019646) | 23/70 | 4.9 864806 848146 E-06 | 0.0 021192 542910 462 | 3.5 457970 580 726 9 | 43. 289856 811596 4 | NDUFA9;COX7B;NDUFA13;NDUFB8;NDUFA11;NDUFA5;COX15;NDUFB5;NDUFA4;COX4I1;NDUFA1;SDHC;UQCR11;SDHD;COX6C;COX6B1;NDUFS8;UQCRQ;UQCRC1;UQCRFS1;NDUFV3;NDUFV2;NDUFV1 |
| KEGG_2019_Mouse | Thermogenesis | 49/231 | 7.0 422685 461693 7E -05 | 0.0 023474 228487 231 2 | 1.9 567931 023 389 | 18. 708889 282813 6 | COX7B;NDUFA13;NDUFA11;COX4I1;PRKAG1;ATP5C1;ATP5G3;ATP5G1;ACTB;CREB3L1;UQCRFS1;PRKACA;CPT1A;RPS6;SDHC;SDHD;ARID1A;COX6B1;CREB3;NDUFS8;UQCRC1;CREB5;NDUFB8;PRKAA2;SMARCD3;KDM3B;COX15;NDUFB5;ATP5A1;ATP5K;ATP5J;ADCY3;UQCR11;ATP5H;ATP5O;ATP5B;NRAS;ATP5E;NDUFV3;SLC25A20;NDUFV2;NDUFV1;NDUFA9;NDUFA5;NDUFA4;NDUFA1;COX6C;SMARCA2;UQCRQ |
| GO_Biological_Process_2021 | mitochondrial ATP synthesis coupled electron transport (GO:0042775) | 23/71 | 6.5 304707 740943 1E -06 | 0.0 025441 625724 075 8 | 3.4 717280 375 120 7 | 41. 449070 479946 9 | NDUFA9;COX7B;NDUFA13;NDUFB8;NDUFA11;NDUFA5;COX15;NDUFB5;NDUFA4;COX4I1;NDUFA1;SDHC;UQCR11;SDHD;COX6C;COX6B1;NDUFS8;UQCRQ;UQCRC1;UQCRFS1;NDUFV3;NDUFV2;NDUFV1 |
| KEGG_2019_Mouse | Fluid shear stress and atherosclerosis | 34/143 | 8.7 854207 626892 9E -05 | 0.0 026356 262288 067 9 | 2.2 624402 297 009 8 | 21. 130811 313584 7 | PRKAA2;NCF1;SDC4;NCF2;GSTP2;ITGA2B;MGST1;KEAP1;MGST2;PIK3R3;CALML4;ACTB;CTSL;AKT2;SUMO3;GPC1;RAC2;AKT1;RAC3;RAC1;MAP2K7;GSTM2;GSTM1;CAV2;CAV1;CYBA;RHOA;ASS1;KLF2;SDC1;CALM3;CALM1;SQSTM1;CALM2 |
| MSigDB_Hallmark_2020 | Complement | 42/200 | 0.0 002800 702980 72 738 | 0.0 028007 029807 273 8 | 1.9 290442 457 322 | 15. 780488 429958 3 | DOCK4;GMFB;CFH;SERPINC1;DOCK9;SERPINE1;PREP;WAS;CBLB;PRCP;CLU;CTSS;GNAI2;C3;DOCK10;DPP4;LGALS3;PCLO;CTSL;S100A13;ME1;SH2B3;CTSC;CD55;FCER1G;F10;FN1;MMP12;MMP14;RABIF;MMP15;GPD2;GNB2;APOC1;IRF7;ADAM9;CALM3;CALM1;RAF1;MSRB1;HSPA1A;DGKH |
| GO_Biological_Process_2021 | rRNA metabolic process (GO:0016072) | 40/162 | 8.4 798635 674634 3E -06 | 0.0 030494 893982 993 5 | 2.3 822404 371 584 7 | 27. 819365 962274 7 | RPL5;RPL30;RPL3;DDX49;RPL32;RPL10;RPL34;RPL12;RRP1;FCF1;RPL36A;RPL10A;RPL8;FBL;RPL7A;RPS16;RPS19;PES1;EXOSC8;RPL36;RPL14;RPLP2;RRP12;RPL15;EXOSC2;RPL17;RPL19;RPL21;RPS6;WDR74;DDX54;RPL35A;IMP4;ERN2;RPS25;BOP1;EBNA1BP2;C1D;RPL27;RPL29 |
| KEGG_2019_Mouse | Gastric acid secretion | 21/74 | 0.0 001432 159036 42 857 | 0.0 039058 882811 688 4 | 2.8 676047 329 709 | 25. 381620 424221 6 | CHRM3;KCNE2;PRKCB;ITPR1;ADCY3;ATP1B3;PRKCA;CALML4;SLC4A2;ATP1A1;ATP1B1;ACTB;GNAI2;PLCB4;CALM3;PRKACA;EZR;CALM1;PLCB2;KCNK2;CALM2 |
| KEGG_2019_Mouse | Fatty acid degradation | 16/50 | 0.0 001905 708054 21 045 | 0.0 047642 701355 261 2 | 3.4 024461 269 656 4 | 29. 143607 219657 5 | GCDH;ACADVL;CPT1A;CYP4A12B;ACAA2;ECHS1;ECI2;ACAA1A;ADH5;ACAT1;ALDH3A2;ALDH2;ACOX1;EHHADH;ACADM;ACADS |
| KEGG_2019_Mouse | Fc gamma R-mediated phagocytosis | 23/87 | 0.0 002272 108064 59 223 | 0.0 052433 263029 051 4 | 2.6 014170 459 247 | 21. 824932 496652 | VAV3;VASP;NCF1;MYO10;PRKCB;ARPC1B;LIMK2;WAS;PIK3R3;PRKCA;ARPC4;ARPC5;FCGR1;CDC42;SCIN;AKT2;CFL1;RAC2;AKT1;RAC1;PIP5K1C;RAF1;PLPP1 |
| GO_Biological_Process_2021 | rRNA processing (GO:0006364) | 41/173 | 1.8 981226 927249 E-05 | 0.0 063383 739917 778 1 | 2.2 564578 675 458 2 | 24. 532345 602572 | RPL5;RPL30;RPL3;DDX49;RPL32;RPL10;RPL34;RPL12;RRP1;MAK16;FCF1;RPL36A;RPL10A;RPL8;FBL;RPL7A;SART1;RPS16;RPS19;PES1;EXOSC8;RPL36;RPL14;RPLP2;RRP12;RPL15;EXOSC2;RPL17;RPL19;RPL21;RPS6;WDR74;DDX54;RPL35A;IMP4;RPS25;BOP1;EBNA1BP2;C1D;RPL27;RPL29 |
| GO_Biological_Process_2021 | peptide biosynthetic process (GO:0043043) | 39/162 | 2.0 473759 809727 E-05 | 0.0 063809 884740 315 9 | 2.3 027092 370 049 1 | 24. 860892 867973 3 | RPL5;RPL30;RPL3;RPL32;RPL10;MRPS14;RPL34;RPL12;RPL36A;FURIN;RRBP1;RPL10A;RPL8;RPS4X;RPL7A;RPS16;RPS15A;RPS19;RPS18;RPL36;RPL14;RPS3;RPLP2;RPL15;RPS27A;RPL17;RPS13;RPS12;RPL19;GGT5;RPL21;RPS6;RPL35A;RPSA;MRPL43;RPS25;COPS5;RPL27;RPL29 |
| MSigDB_Hallmark_2020 | Protein Secretion | 23/96 | 0.0 010240 236190 95 33 | 0.0 075546 464202 162 3 | 2.2 795211 997 211 5 | 15. 692259 697975 8 | ABCA1;STX12;ARF1;TMED10;DST;CAV2;GOSR2;BNIP3;ATP1A1;EGFR;LMAN1;GOLGA4;VAMP7;TMX1;AP2S1;TMED2;ERGIC3;YKT6;AP2M1;CTSC;COPE;VAMP3;BET1 |
| MSigDB_Hallmark_2020 | mTORC1 Signaling | 40/200 | 0.0 010576 504988 30 27 | 0.0 075546 464202 162 3 | 1.8 125 | 12. 418715 931452 5 | PPP1R15A;TOMM40;PSMD13;PIK3R3;SLA;ENO1;SLC7A11;SLC1A5;HMGCR;IFI30;CFP;ADD3;CORO1A;ADIPOR2;LDHA;SERP1;NAMPT;ME1;SSR1;SC5D;ABCF2;YKT6;LDLR;CTSC;ACTR3;TPI1;IGFBP5;TXNRD1;IMMT;ELOVL6;SYTL2;QDPR;SLC7A5;UFM1;PFKL;COPS5;PSMC4;CD9;SQSTM1;GAPDH |
| KEGG_2019_Mouse | Chemokine signaling pathway | 40/197 | 0.0 007795 289234 25 977 | 0.0 167041 912162 709 | 1.8 474522 292 993 6 | 13. 221884 477095 3 | ITK;CXCL9;NCF1;ROCK1;SHC1;WAS;PIK3R3;ADCY3;ARRB1;CXCL3;GNAI2;RAP1B;CDC42;NRAS;GRK3;RAP1A;CXCR1;AKT2;CXCR2;GRK6;RAC2;AKT1;RAC3;RAC1;PRKACA;VAV3;CCL25;PRKCB;BAD;STAT2;STAT3;BRAF;RHOA;FGR;PLCB4;PARD3;GNB2;GNB1;RAF1;PLCB2 |
| KEGG_2019_Mouse | Citrate cycle (TCA cycle) | 11/32 | 0.0 009673 070489 62 897 | 0.0 186482 579650 658 | 3.7 822541 120 194 5 | 26. 252605 312686 6 | CS;PCX;MDH2;IDH3G;DLST;SDHC;SUCLG1;DLAT;SDHD;FH1;IDH3A |
| KEGG_2019_Mouse | Renal cell carcinoma | 18/68 | 0.0 010332 352119 31 77 | 0.0 186482 579650 658 | 2.6 026424 442 609 4 | 17. 893324 047785 5 | ARNT2;CREBBP;EGLN2;BAD;PIK3R3;BRAF;FH1;RAP1B;CDC42;NRAS;RAP1A;AKT2;AKT1;RAC1;ELOC;VHL;RAF1;MET |
| KEGG_2019_Mouse | Glucagon signaling pathway | 24/102 | 0.0 010567 346180 20 4 | 0.0 186482 579650 658 | 2.2 264391 237 901 2 | 15. 256833 674802 7 | CPT1A;CREBBP;PRKAA2;PRMT1;ITPR1;PRKAG1;CALML4;PPP3CA;CREB3;PPP3R1;LDHA;PFKL;G6PC3;PLCB4;CREB3L1;AKT2;AKT1;CALM3;PRKACA;CALM1;CALM2;PLCB2;ATF4;CREB5 |
| GO_Biological_Process_2021 | cellular response to oxidative stress (GO:0034599) | 31/125 | 7.6 644377 574382 3E -05 | 0.0 223945 290725 148 | 2.3 910601 202 935 8 | 22. 658485 048681 3 | VKORC1L1;PRKAA2;NCF1;SRXN1;NCF2;LRRK2;KEAP1;PRR5L;PTPRK;EGFR;ARNTL;PRDX3;PRDX2;ALDH3B1;LONP1;RPS3;AKT1;SLC25A24;GPX1;TXNRD1;TPM1;GPX8;SLC4A11;CYBA;VRK2;SELENOS;CAT;CPEB2;HSPA1B;ATF4;HSPA1A |
| GO_Biological_Process_2021 | fatty acid beta-oxidation (GO:0006635) | 17/52 | 8.9 810297 082343 5E -05 | 0.0 246978 316976 445 | 3.5 130593 714 993 2 | 32. 734022 982756 1 | GCDH;ACADVL;ACAA2;MECR;ECHS1;ECI2;ETFDH;ETFA;ACAT1;SLC25A17;SCP2;ACOX1;EHHADH;ACADM;ABCD1;ACADS;PPARD |
| KEGG_2019_Mouse | Regulation of actin cytoskeleton | 42/217 | 0.0 015747 065761 68 38 | 0.0 262451 096028 064 | 1.7 399499 582 985 8 | 11. 229091 251892 6 | ITGB1;CHRM3;ROCK1;ITGB4;ARPC1B;ITGA2B;WAS;LPAR2;PIK3R3;BRK1;ACTB;MYL12A;EGFR;PPP1CB;CDC42;NRAS;SCIN;TMSB4X;CFL1;RAC2;ITGAX;RAC3;MYH14;PIP5K1C;RAC1;VAV3;PPP1R12A;LIMK2;FN1;BRAF;ARPC4;ARPC5;RHOA;PPP1CA;ENAH;PIKFYVE;DIAPH3;ITGA10;EZR;RAF1;FGFR3;PFN2 |
| KEGG_2019_Mouse | Mineral absorption | 13/44 | 0.0 017113 770048 47 82 | 0.0 270217 421818 077 | 3.0 287890 266 757 | 19. 294770 155128 | SLC31A1;SLC40A1;SLC30A1;MT2;ATP1B3;ATP1A1;MT1;ATP1B1;SLC8A1;FTL1;HEPH;FTH1;STEAP2 |
| GO_Biological_Process_2021 | protein N-linked glycosylation via asparagine (GO:0018279) | 12/30 | 0.0 001095 789742 51 26 | 0.0 284600 947013 69 | 4.8 165842 943 437 7 | 43. 921782 146559 2 | DPM1;LMAN1;ST6GAL1;RPN2;TUSC3;DAD1;DERL3;ALG5;MGAT1;MGAT2;DDOST;UBE2J1 |
| KEGG_2019_Mouse | Platelet activation | 27/125 | 0.0 020830 236894 27 04 | 0.0 312453 553414 056 | 1.9 937667 565 990 3 | 12. 309385 850045 5 | ITGB1;ROCK1;ITGA2B;ITPR1;PIK3R3;ADCY3;ACTB;MYL12A;GNAI2;PTGS1;PPP1CB;RAP1B;RAP1A;AKT2;AKT1;PRKACA;VASP;P2RY12;PPP1R12A;FCER1G;VWF;RHOA;PPP1CA;VAMP8;PLCB4;TBXAS1;PLCB2 |
| KEGG_2019_Mouse | Non-alcoholic fatty liver disease (NAFLD) | 31/151 | 0.0 023952 408584 10 67 | 0.0 342177 265487 238 | 1.8 702089 387 020 9 | 11. 285348 471653 1 | NDUFA13;COX7B;NDUFB8;PRKAA2;NDUFA11;NDUFB5;COX4I1;PRKAG1;PIK3R3;UQCR11;ADIPOR2;CDC42;AKT2;AKT1;UQCRFS1;NDUFV3;RAC1;NDUFV2;NDUFV1;NDUFA9;NDUFA5;NDUFA4;NDUFA1;SDHC;SDHD;COX6C;COX6B1;NDUFS8;UQCRQ;UQCRC1;ATF4 |
| GO_Biological_Process_2021 | cellular response to chemical stress (GO:0062197) | 26/101 | 0.0 001470 229696 94 224 | 0.0 361753 885958 157 | 2.5 109638 221 485 8 | 22. 159059 189712 1 | VKORC1L1;PRKAA2;NCF1;SRXN1;NCF2;LRRK2;KEAP1;PRR5L;PRDX3;PRDX2;ALDH3B1;LONP1;SLC25A24;GPX1;CAV1;TXNRD1;GPX8;SLC4A11;CYBA;VRK2;SELENOS;CAT;CPEB2;HSPA1B;ATF4;HSPA1A |
| KEGG_2019_Mouse | Leukocyte transendothelial migration | 25/115 | 0.0 027235 796271 29 1 | 0.0 371397 221881 241 | 2.0 094317 920 404 9 | 11. 867318 804023 3 | ITGB1;ITK;NCF1;ROCK1;NCF2;TXK;PIK3R3;F11R;ACTB;MYL12A;GNAI2;RAP1B;CDC42;RAP1A;RASSF5;RAC2;RAC1;VASP;VAV3;PRKCB;CYBA;PRKCA;RHOA;CLDN3;EZR |
| GO_Biological_Process_2021 | peptidyl-asparagine modification (GO:0018196) | 12/31 | 0.0 001590 344399 02 511 | 0.0 371743 003272 119 | 4.5 628197 346 744 1 | 39. 908199 877263 8 | DPM1;LMAN1;ST6GAL1;RPN2;TUSC3;DAD1;DERL3;ALG5;MGAT1;MGAT2;DDOST;UBE2J1 |
| KEGG_2019_Mouse | Bacterial invasion of epithelial cells | 18/74 | 0.0 028893 877891 24 62 | 0.0 376876 668146 689 | 2.3 229916 243 954 2 | 13. 581859 839948 9 | ITGB1;SHC1;CAV2;ARPC1B;CAV1;WAS;FN1;ILK;PIK3R3;ARPC4;ARPC5;ACTB;RHOA;SEPT8;CDC42;SEPT9;RAC1;MET |
| KEGG_2019_Mouse | VEGF signaling pathway | 15/58 | 0.0 033556 926971 82 51 | 0.0 419461 587147 814 | 2.5 198273 795 252 9 | 14. 355701 162534 1 | PRKCB;BAD;PIK3R3;PRKCA;CDC42;PPP3CA;NRAS;PPP3R1;MAPKAPK3;AKT2;RAC2;AKT1;RAC3;RAC1;RAF1 |
| MSigDB_Hallmark_2020 | Cholesterol Homeostasis | 17/74 | 0.0 068979 069947 96 55 | 0.0 431119 187174 785 | 2.1 544337 525 613 5 | 10. 721619 816852 8 | GSTM2;PCYT2;ECH1;GPX8;HMGCR;CLU;LSS;LGALS3;ATXN2;ALCAM;FABP5;FASN;SC5D;MAL2;CD9;LDLR;CPEB2 |
| GO_Biological_Process_2021 | astral microtubule organization (GO:0030953) | 6/9 | 0.0 001980 897548 32 943 | 0.0 440985 525640 004 | 14. 426458 504 519 3 | 123 .01138 676782 2 | DLG1;NUMA1;CENPJ;LIMK2;EZR;CLASP1 |

**Figure S3: Summary of all immune tone and lung injury measurements for all 3 groups.** Comprehensive data from all measurements conducted including all three groups: FF, FR (1 week) and FR (2 weeks) with individual mouse #s annotated. Plasma and lung lung injury measurements and left and right lung inflammation marker measurements (A) in mice after lung IR injury. SCFA measurements in stool and portal blood (B) and plasma and Right Lung (C). n=8-15 mice in each group depending on the specific transcriptomic/proteomic/metabolite measurement.

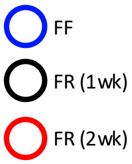


S3A

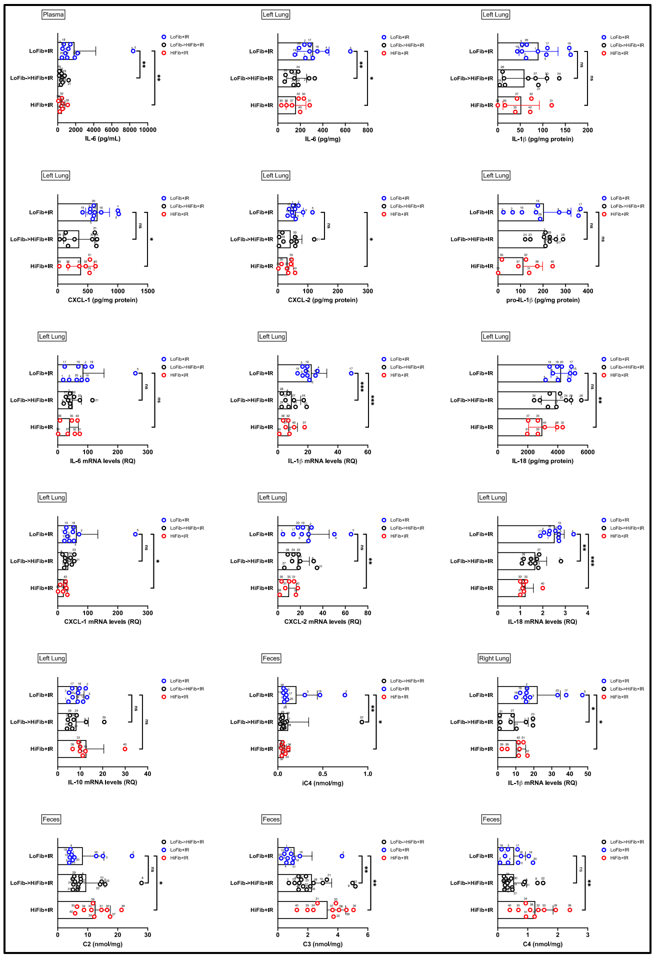

S3B

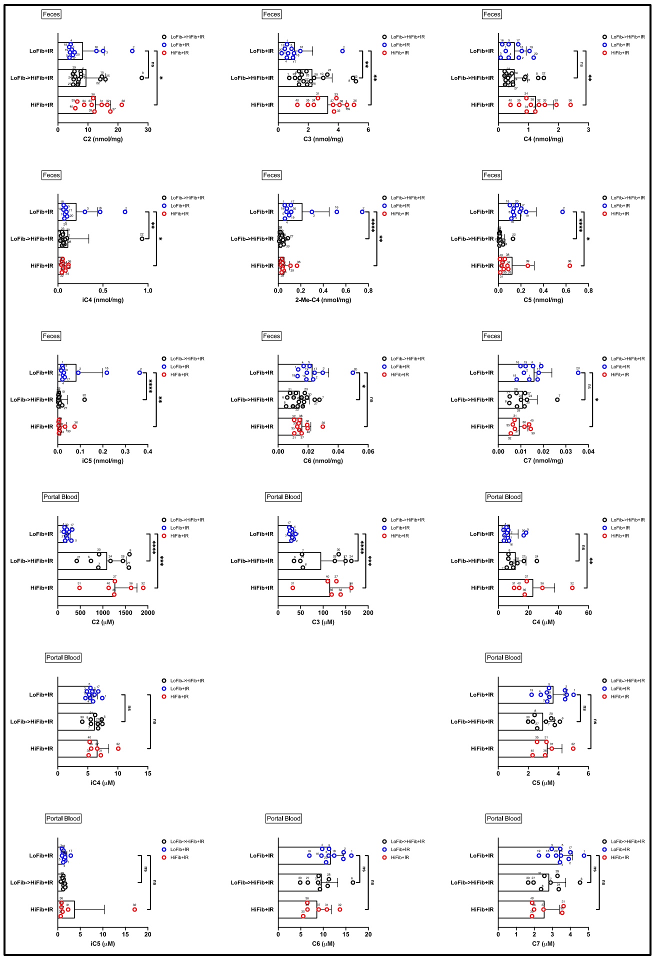

S3C

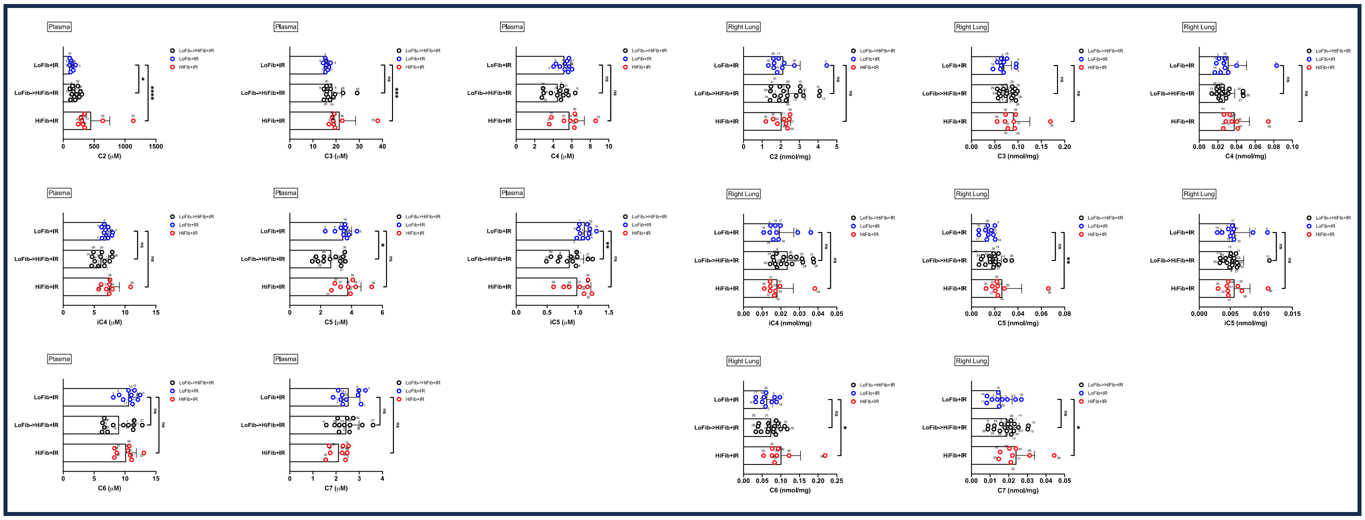

**Supplemental Table S2: Correlation table (all Groups) and including 16S data.** Summary of individual correlation coefficients between bacteria phyla, stool SCFAs, immune tone and lung injury markers. n=26-27 mice.

Correlation Table (SILVA)

| Symbol | Actinobacteriota | Bacteria_Total | Bacteroidota | Firmicutes | F/ B SILVA | OTU_Count | Proteobacteria | Verrucomicrobiota | Caproate (F) | Heptanoate (F) | Propionate (F) | CXCL1 ELISA (LL) | IL-1b ELISA (LL) | IL-1b qPCR (LL) | IL-1B qPCR (RL) | IL-18 qPCR (LL) |
|---|---|---|---|---|---|---|---|---|---|---|---|---|---|---|---|---|
| "Actinobacteriota*" | | −0.379* | | | | | −0.376* | | | | | 0.587** | 0.396* | 0.49* | | 0.475* |
| "Bacteroidota*" | | | | | | | | −0.415* | −0.429* | −0.468** | 0.394* | | −0.442* | | −0.536** | |
| "Firmicutes*" | | | | | | | −0.485** | | 0.375* | 0.451** | −0.358* | 0.612** | 0.441* | 0.518** | 0.462* | |
| "Proteobacteria*" | −0.405* | | | −0.521** | −0.511** | | | −0.523** | −0.412* | | | −0.609** | −0.458* | | | −0.452* |
| "Verrucomicrobiota*" | | 0.371* | | | | 0.479** | −0.521** | | 0.339* | | | | | 0.471* | | |
| FECES | | | | | | | | | | | | | | | | |
| Acetate | | | | | | | | | | | | | | | | |
| Propionate | | | | −0.448** | | −0.418* | | | | −0.496** | | | −0.4* | | −0.446* | |
| Butyrate | | | | | | | | | | | | | | | | −0.427* |
| Isobutyrate | | | | | | | | | | | 0.339* | | | | | |
| 2-methylbutyrate | 0.377* | | −0.441* | 0.488** | 0.42* | | −0.375* | | | | | 0.404* | | | | |
| Valerate | 0.439* | | −0.467** | 0.384* | 0.408* | | | | 0.349* | | | | | | | |
| Isovalerate | 0.382* | | −0.386* | 0.416* | 0.386* | | −0.345* | | | | | 0.407* | | | | |
| Caproate | | | −0.344* | 0.417* | 0.356* | | | | | | | | | | 0.395* | |
| Heptanoate | | | −0.34* | 0.502** | 0.411* | | | | | | −0.496** | | | | 0.549** | 0.573** |
| LEFT LUNG | | | | | | | | | | | | | | | | |
| CXCL1 ELISA | 0.562** | | −0.571** | 0.531** | | | −0.569** | | | | | | | | | 0.403* |
| CXCL1 qPCR | | | | 0.402* | | | | | | | | | | | 0.584** | |
| CXCL2 ELISA | | | | | | | −0.508** | | | | | | | 0.437* | | |
| CXCL2 qPCR | | | −0.398* | 0.477* | 0.487* | | | | | | | 0.452* | 0.419* | | | 0.526** |
| IL-1b ELISA | 0.413* | | | 0.391* | 0.481* | | −0.418* | | | | −0.4* | | | 0.593** | | |
| IL-1b qPCR | 0.573** | | −0.607** | 0.537** | 0.497* | | | 0.464* | | | | | 0.593** | | | 0.431* |
| IL-6 ELISA | 0.405* | | −0.461* | 0.558** | 0.445* | | −0.414* | | | | −0.476* | 0.564** | 0.594** | 0.437* | | 0.469* |
| IL-6 qPCR | | | | 0.471* | | | | | | | | 0.416* | 0.428* | | 0.491* | |
| IL-10 qPCR | | | | | | | | | | | | | | | | |
| IL-18 ELISA | | | | | | | | | | | | | | | | |
| IL-18 qPCR | 0.488* | | −0.584** | | | 0.457* | | | | 0.573** | | 0.403* | | 0.431* | 0.398* | |
| pro-IL-1b ELISA | | | | | | | | | | | | | 0.41* | 0.402* | | |

**Figure S4: Correlations between immune tone and lung injury markers, SCFAs, and bacterial taxa (all Groups).** Significant positive correlations between cytokines/chemokines, between left and right lung IL-1b levels (row 1 and 2), and between metabolites (row 3) and negative correlation between heptanoate and propionate (row 4). n=26-27 mice as shown on each graph.

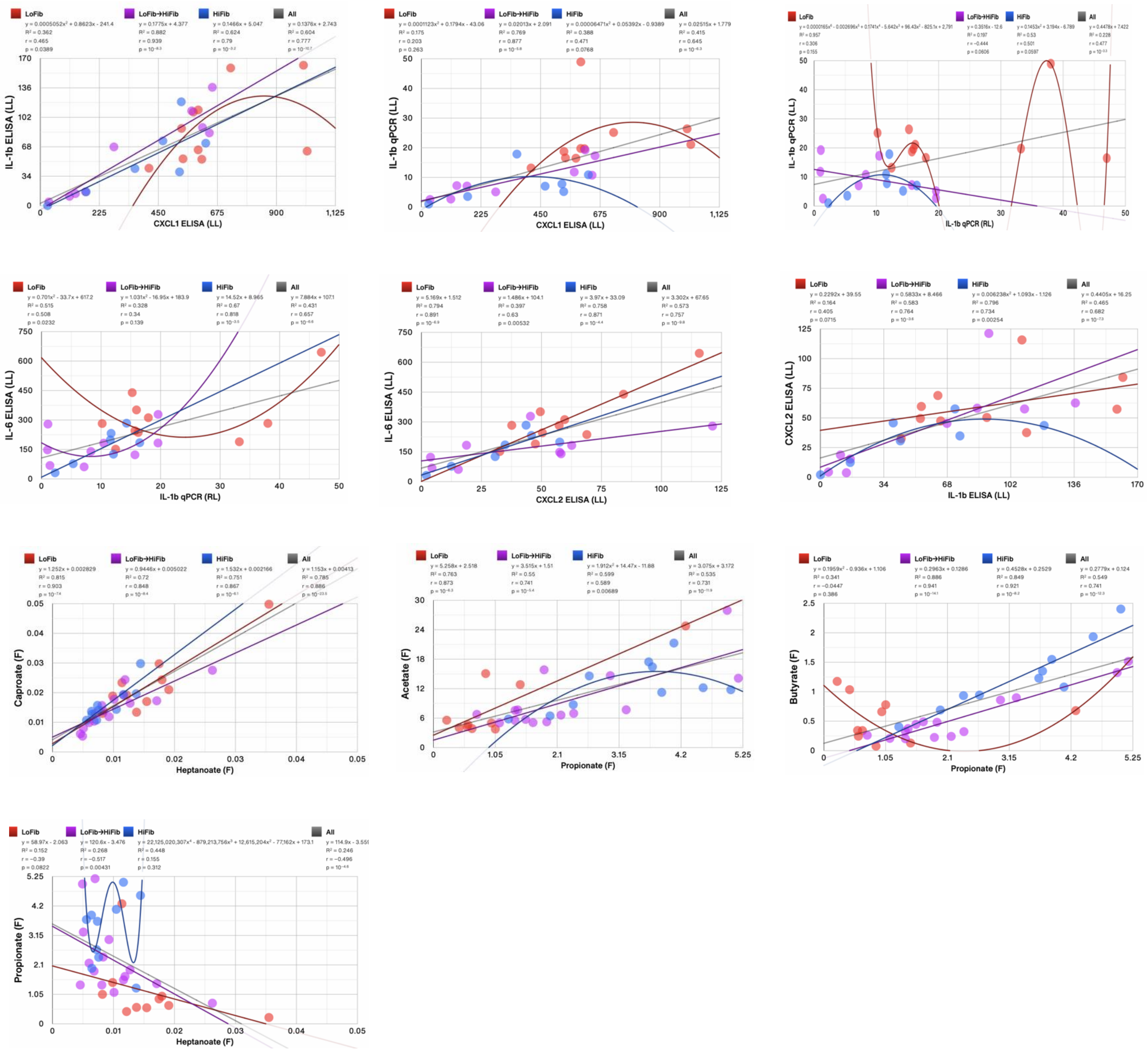

**Figure S5: FMT transfer of FF/FR Lung Injury phenotype to GF mice. Comparing the Left (Directly Injured) vs. Right (Indirectly Injured) Lungs.** Cytokines and chemokine levels measured by qPCR from lung tissue homogenates.

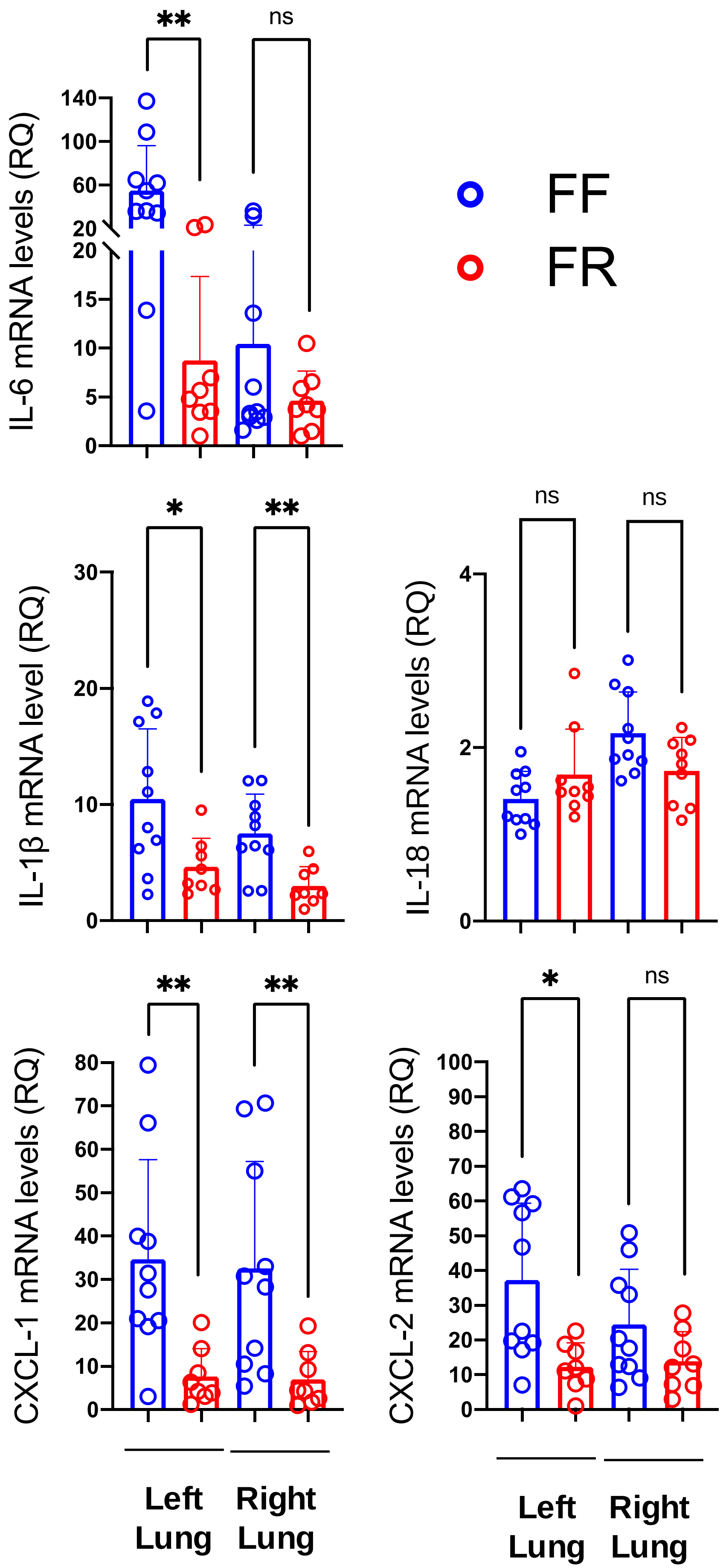

**Supplemental Table S3: *B.theta* strain information**

| Relation | Strain | Genotype | Phenotype/Role |
|---|---|---|---|
| **Parent strain** | VPI-5482 | Wild-type, intact genome | Standard reference strain; fully functional; sensitive to 2-FdU. NCBI Reference Genome: NC_004663 |
| **Genetic mutant parent strain** | DeltaTDK | *tdk* gene (BT_4729 Thymidine Kinase Gene) deleted | Gene ID: BT_4729 in VPI-5482 genome Resistant to 2-FdU; used as a platform for genetic engineering; serves as the "wild-type" in experimental contexts. |
| **Propionate mutant strain** | Delta1686-89 | *tdk* gene deleted, BT_1686-89 deleted | Propionate metabolism deficient; used to study SCFA metabolism and its effects on microbial and host physiology. |

**Figure S6: 16S rRNA sequence analysis of FR and FF mice gut microbiota from snATACseq experiment.** Family level comparison between n=4 mice gut microbiota in each group.

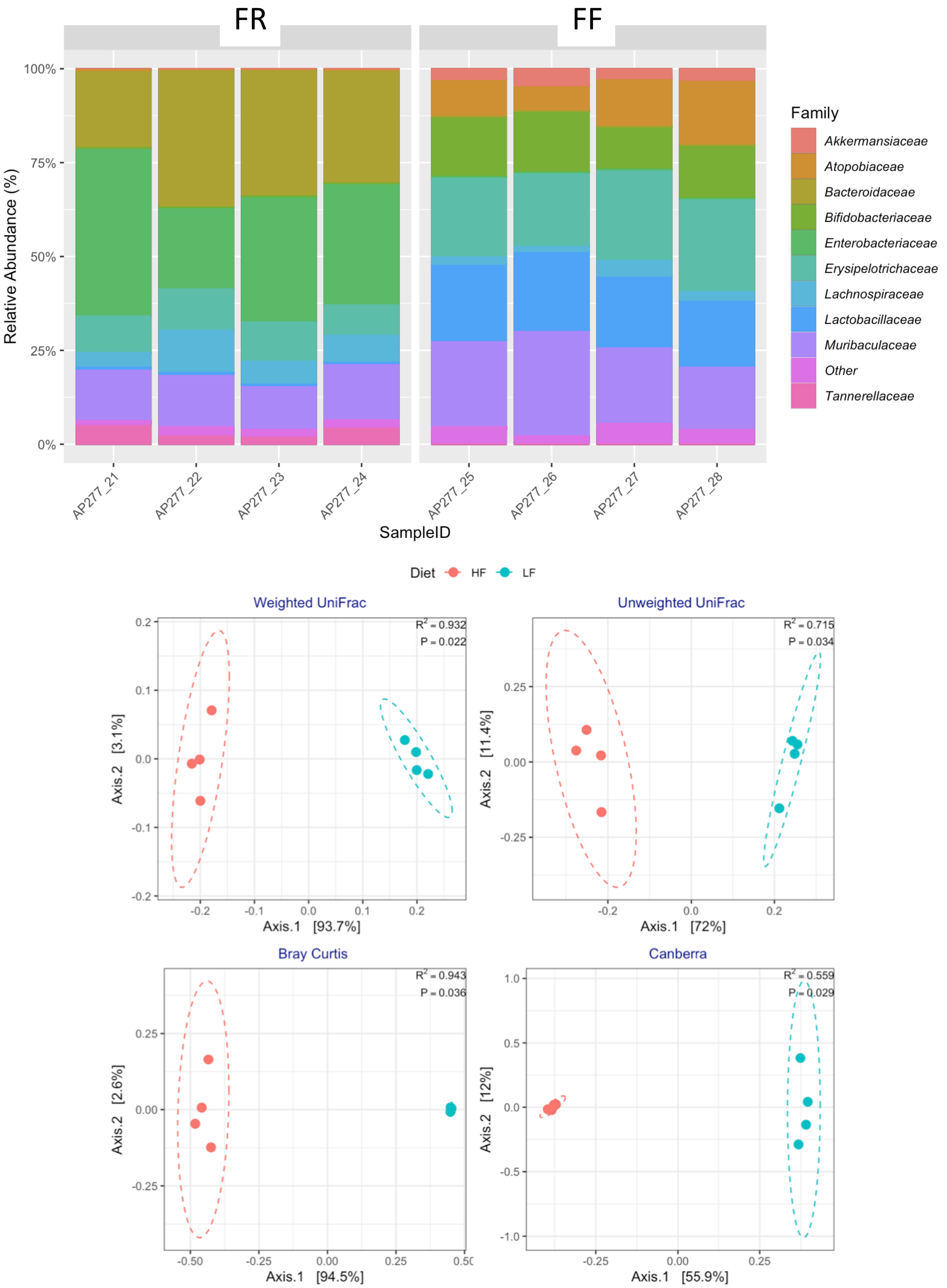

**Figure S7: Manhattan Plots for Chromatin Accessibility loci from snATACseq.**

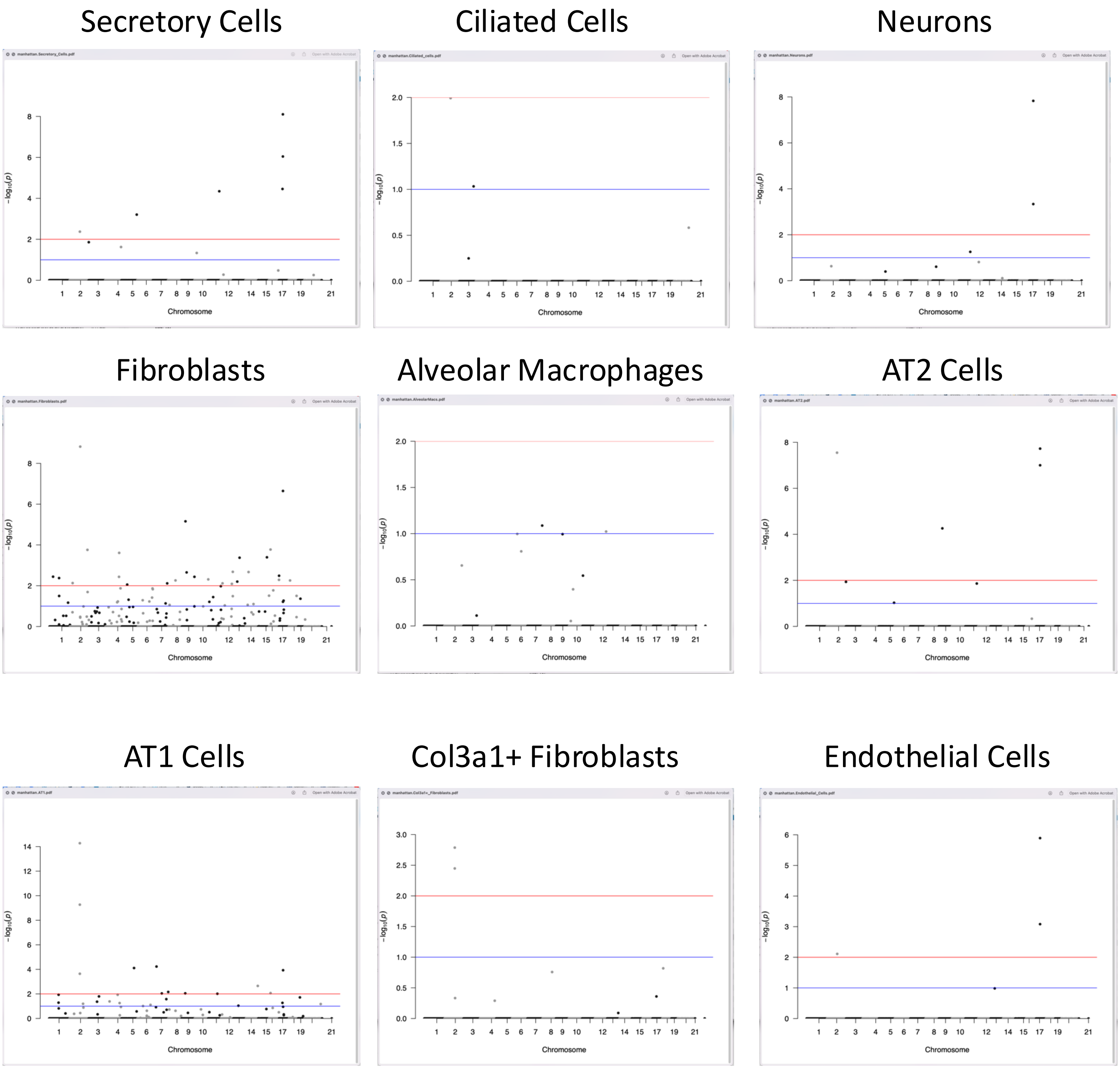

**Table S4: Cell Populations from digested lungs of FR and FF mice (pre-injury) from snATACseq analysis.** (A) Cell types that were significantly different are highlighted in yellow. (B) Differentially expressed genes with adjusted p values <0.1 for each cell type identified. (C) Number of Differentially sequenced ATAC chromatin segments with adjusted p values <0.1

#### Table S4A

| BaselineProp.clusters | BaselineProp.Freq | PropMean.HighFiber | PropMean.LowFiber | PropRatio | Tstatistic | P.Value | FDR |
|---|---|---|---|---|---|---|---|
| Macrophages | 0.0641549402330156 | 0.0799086280096742 | 0.0151448576652441 | 5.27628781834355 | 5.07910491440752 | 2.25293885393785e-06 | 3.15411439551299e-05 |
| CD4_T_cell | 0.052050234528673 | 0.059440250737777 | 0.0185911074675084 | 3.19724098425338 | 3.43411094617065 | 0.000925663644030091 | 0.00647964550821064 |
| B_cell | 0.0494779845665002 | 0.0585447798984303 | 0.0214024589587154 | 2.73542306570293 | 3.17298942313917 | 0.00210818430297654 | 0.00791508806846228 |
| Col3a1+_Fibroblasts | 0.0680889695869269 | 0.0421565480396635 | 0.109397686892566 | 0.38535136561949 | -3.15008691582251 | 0.00226145373384637 | 0.00791508806846228 |
| Alveolar Macs | 0.127099409895597 | 0.100771353106679 | 0.170008543833376 | 0.592742875354807 | -1.8168421948071 | 0.0728078201794595 | 0.203861896502486 |
| Mast_cells | 0.0220910879104252 | 0.0259456327949061 | 0.0070893501465127 | 3.65980410879676 | 1.96859318259878 | 0.0961911342133788 | 0.224445979831217 |
| Fibroblasts | 0.115297321833863 | 0.0931846963794225 | 0.148313211538281 | 0.628296666311285 | -1.37751201140774 | 0.172013279238302 | 0.344026558476604 |
| AT1 | 0.0969889544560448 | 0.0876001099811132 | 0.113472416566582 | 0.771994751074262 | -0.715401940669935 | 0.476344551623919 | 0.753011270496573 |
| Mesothelial_Cells | 0.0140717203812982 | 0.0119536196483763 | 0.0149230486592283 | 0.801017266735526 | -0.697239063789537 | 0.487578477408955 | 0.753011270496573 |
| Neurons | 0.0832198517173551 | 0.105061403468425 | 0.0806240255146501 | 1.30310292493811 | 0.595114674872549 | 0.553366677731809 | 0.753011270496573 |
| Endothelial_Cells | 0.10425177787865 | 0.121397443925048 | 0.0991189326849993 | 1.224765447292 | 0.538506358827145 | 0.591651712533022 | 0.753011270496573 |
| Ciliated_cells | 0.0484188228173703 | 0.0494491605878084 | 0.0498935863467694 | 0.99109252728653 | 0.401130232533909 | 0.690789910656814 | 0.781324434701315 |
| AT2 | 0.0868512634286579 | 0.0950587383572102 | 0.0829293404880106 | 1.1462618392697 | 0.352273379254484 | 0.725515546508364 | 0.781324434701315 |
| Secretory_Cells | 0.0679376607656226 | 0.0695276350654654 | 0.0690914332375567 | 1.00631339961365 | -0.0285153688864745 | 0.97731874977729 | 0.97731874977729 |

#### Table S4B

| cellType | Significant Gene Count |
|---|---|
| Endothelial_Cells | 2 |
| Col3a1+_Fibroblasts | 3 |
| CD4_T_cell | 1 |
| AT1 | 13 |
| AT2 | 4 |
| Alveolar_Macs | 1 |
| B_cell | 0 |
| AlveolarMacs | 8 |
| Fibroblasts | 34 |
| Neurons | 2 |
| Macrophages | 0 |
| Ciliated_cells | 2 |
| Mast_cells | 0 |
| Secretory_Cells | 1 |
| Mesothelial_Cells | 1 |

#### Table S4C

| cellType | Significant Gene Count |
|---|---|
| Endothelial_Cells | 3 |
| Col3a1+_Fibroblasts | 2 |
| CD4_T_cell | 0 |
| AT1 | 27 |
| AT2 | 7 |
| Alveolar_Macs | 0 |
| B_cell | 0 |
| AlveolarMacs | 3 |
| Fibroblasts | 53 |
| Neurons | 3 |
| Macrophages | 0 |
| Ciliated_cells | 2 |
| Mast_cells | 0 |
| Secretory_Cells | 9 |
| Mesothelial_Cells | 0 |

Supplemental Table S4D Differentially expressed genes with adjusted p values <0.1 or top hit for each cell type identified

| Ciliated Cells | p_val | avg_log2FC | pct.1 | pct.2 | p_val_adj |
|---|---|---|---|---|---|
| Gm10800 | 8.45E-08 | 0.56792722 | 0.973 | 0.917 | 0.00184337 |
| Sfi1 | 3.15E-06 | 0.53869866 | 0.968 | 0.917 | 0.06864342 |
| Smim20 | 4.61E-06 | -1.037833 | 0.293 | 0.621 | 0.10066109 |

| Endothelial Cells | p_val | avg_log2FC | pct.1 | pct.2 | p_val_adj |
|---|---|---|---|---|---|
| Gm10800 | 4.14E-08 | 0.37904698 | 0.986 | 0.974 | 0.00090343 |
| Prl8a9 | 1.76E-06 | -2.2512669 | 0.034 | 0.136 | 0.0383567 |

| Endothelial Cells | p_val | avg_log2FC | pct.1 | pct.2 | p_val_adj |
|---|---|---|---|---|---|
| Gm10800 | 4.24E-10 | 0.6181366 | 0.955 | 0.89 | 9.25E-06 |

| Macrophages | p_val | avg_log2FC | pct.1 | pct.2 | p_val_adj |
|---|---|---|---|---|---|
| Slc5a8 | 6.36E-05 | 3.11274674 | 0.284 | 0.05 | 1 |

| Neurons | p_val | avg_log2FC | pct.1 | pct.2 | p_val_adj |
|---|---|---|---|---|---|
| Gm10800 | 3.58E-07 | 0.44388716 | 0.983 | 0.98 | 0.00780578 |
| Ftsj3 | 3.81E-06 | -1.044998 | 0.224 | 0.436 | 0.08324653 |

| CD4 T Cells | p_val | avg_log2FC | pct.1 | pct.2 | p_val_adj |
|---|---|---|---|---|---|
| Mterf4 | 1.51E-06 | -1.8794499 | 0.113 | 0.481 | 0.03294199 |

| B Cells | p_val | avg_log2FC | pct.1 | pct.2 | p_val_adj |
|---|---|---|---|---|---|
| Gm16506 | 6.92E-05 | -2.2721327 | 0.049 | 0.226 | 1 |


| Ciliated Cells | p_val | avg_log2FC | pct.1 | pct.2 | p_val_adj |
|---|---|---|---|---|---|
| Gm10800 | 8.45E-08 | 0.56792722 | 0.973 | 0.917 | 0.00184337 |
| Sfi1 | 3.15E-06 | 0.53869866 | 0.968 | 0.917 | 0.06864342 |
| Smim20 | 4.61E-06 | -1.037833 | 0.293 | 0.621 | 0.10066109 |


| Mast Cells | p_val | avg_log2FC | pct.1 | pct.2 | p_val_adj |
|---|---|---|---|---|---|
| Gm773 | 4.01E-05 | -3.4859614 | 0.054 | 0.438 | 0.87582308 |

| Mesothelial Cells | p_val | avg_log2FC | pct.1 | pct.2 | p_val_adj |
|---|---|---|---|---|---|
| Oasl1 | 2.73E-06 | -3.7321952 | 0.038 | 0.415 | 0.05961844 |

Supplemental Table S4D Differentially expressed genes with adjusted p values <0.1 or top hit for each cell type identified

| AT1 | p_val | avg_log2FC | pct.1 | pct.2 | p_val_adj |
|---|---|---|---|---|---|
| Gm10800 | 3.37E-22 | 1.09882049 | 0.891 | 0.747 | 7.35E-18 |
| Gm10801 | 3.78E-09 | 0.53812844 | 0.894 | 0.795 | 8.25E-05 |
| Fah | 9.66E-08 | -0.7829363 | 0.413 | 0.634 | 0.00210871 |
| Drg1 | 2.50E-07 | 0.51154178 | 0.894 | 0.872 | 0.00545063 |
| Sfi1 | 2.93E-07 | 0.5009006 | 0.886 | 0.839 | 0.00639399 |
| Gm21738 | 4.13E-07 | 1.00793447 | 0.481 | 0.388 | 0.00900328 |
| Hrct1 | 6.61E-07 | -1.016074 | 0.188 | 0.385 | 0.01442993 |
| Rsph1 | 1.41E-06 | -1.1631479 | 0.201 | 0.377 | 0.03085256 |
| Tor3a | 1.88E-06 | -0.5698206 | 0.522 | 0.729 | 0.04106224 |
| Gramd2 | 2.43E-06 | -0.4901285 | 0.69 | 0.828 | 0.05292233 |
| Gm10840 | 2.56E-06 | -0.9740725 | 0.201 | 0.392 | 0.05580245 |
| Pip4k2a | 2.56E-06 | 0.79724203 | 0.606 | 0.549 | 0.05591921 |
| Rrp36 | 3.98E-06 | -1.1500449 | 0.101 | 0.271 | 0.08684494 |

| AT2 | p_val | avg_log2FC | pct.1 | pct.2 | p_val_adj |
|---|---|---|---|---|---|
| Gm10800 | 1.58E-15 | 0.6378608 | 0.977 | 0.977 | 3.44E-11 |
| Drg1 | 5.15E-08 | 0.45680282 | 0.963 | 0.904 | 0.00112404 |
| Sfi1 | 1.19E-06 | 0.41123116 | 0.938 | 0.922 | 0.0258644 |
| Gm21738 | 2.45E-06 | 0.60591827 | 0.761 | 0.74 | 0.05340354 |

| AlveolarMacs | p_val | avg_log2FC | pct.1 | pct.2 | p_val_adj |
|---|---|---|---|---|---|
| Skint6 | 8.15E-08 | 0.92074443 | 0.737 | 0.593 | 0.00177925 |
| Adamtsl1 | 5.23E-07 | 0.80258909 | 0.758 | 0.657 | 0.01141027 |
| Myo5b | 1.75E-06 | 1.02808589 | 0.716 | 0.61 | 0.03827214 |
| Il12rb2 | 1.90E-06 | -0.7073646 | 0.395 | 0.663 | 0.04156586 |
| Fam49a | 2.42E-06 | -0.5545763 | 0.658 | 0.837 | 0.05290664 |
| Rtn4 | 2.53E-06 | -0.4956005 | 0.689 | 0.86 | 0.05528988 |
| Calcoco1 | 3.67E-06 | -0.890162 | 0.274 | 0.513 | 0.08006858 |
| Phlda1 | 3.98E-06 | -0.6907483 | 0.305 | 0.573 | 0.08692424 |

| Col1a1+ Fibroblasts | p_val | avg_log2FC | pct.1 | pct.2 | p_val_adj |
|---|---|---|---|---|---|
| Gm10800 | 1.22E-08 | 0.91004407 | 0.87 | 0.721 | 0.00026657 |
| Tspan9 | 1.58E-07 | -0.5363486 | 0.903 | 0.962 | 0.00345675 |
| Lrfn5 | 1.26E-06 | 0.97046371 | 0.676 | 0.442 | 0.02758949 |

Supplemental Table S4D Differentially expressed genes with adjusted p values <0.1 or top hit for each cell type identified

| Fibroblasts | p_val | avg_log2FC | pct.1 | pct.2 | p_val_adj |
|---|---|---|---|---|---|
| Gm10800 | 2.06E-13 | 0.66117124 | 0.917 | 0.764 | 4.49E-09 |
| Slc22a13 | 1.85E-09 | -1.4430269 | 0.134 | 0.31 | 4.04E-05 |
| Slc29a1 | 2.42E-09 | -0.9072987 | 0.368 | 0.586 | 5.28E-05 |
| Slc7a10 | 8.06E-09 | -0.8362386 | 0.438 | 0.63 | 0.00017597 |
| Mta1 | 1.11E-08 | -0.8220345 | 0.401 | 0.597 | 0.0002429 |
| Serpina3n | 1.69E-08 | -1.2832704 | 0.096 | 0.268 | 0.00036836 |
| Crip2 | 4.18E-08 | -0.8646236 | 0.32 | 0.518 | 0.00091196 |
| Siah2 | 8.91E-08 | -0.7102475 | 0.322 | 0.54 | 0.00194405 |
| Hif3a | 1.45E-07 | -0.5839225 | 0.471 | 0.688 | 0.00316101 |
| Pdzrn4 | 1.64E-07 | 0.71374901 | 0.788 | 0.655 | 0.00358156 |
| Gas1 | 3.14E-07 | -0.9599364 | 0.217 | 0.405 | 0.00684282 |
| Rnase10 | 3.59E-07 | -0.8020289 | 0.254 | 0.458 | 0.00782372 |
| Pex5l | 4.81E-07 | 0.78047252 | 0.66 | 0.54 | 0.01048769 |
| Tmem241 | 4.97E-07 | -0.5346199 | 0.657 | 0.784 | 0.01083553 |
| Tmigd1 | 5.44E-07 | -0.9546125 | 0.164 | 0.334 | 0.01186338 |
| Nxph1 | 8.33E-07 | 0.51962801 | 0.7 | 0.603 | 0.01817245 |
| Zbtb16 | 8.35E-07 | -0.4788074 | 0.854 | 0.882 | 0.01822505 |
| Mroh2a | 9.30E-07 | 0.66963062 | 0.751 | 0.652 | 0.02029348 |
| Ddc | 1.06E-06 | -0.5300688 | 0.592 | 0.751 | 0.0232049 |
| Htra4 | 1.12E-06 | -0.7712884 | 0.181 | 0.367 | 0.02451405 |
| Chchd5 | 1.25E-06 | -0.7351588 | 0.207 | 0.416 | 0.02717745 |
| Lrp3 | 1.40E-06 | -0.9774273 | 0.285 | 0.441 | 0.03057343 |
| Sfi1 | 1.43E-06 | 0.49682509 | 0.821 | 0.751 | 0.03111153 |
| Styx | 1.61E-06 | -0.8896195 | 0.224 | 0.411 | 0.03523635 |
| Hs3st5 | 1.76E-06 | 0.78275767 | 0.668 | 0.584 | 0.03833655 |
| Dync1i1 | 2.14E-06 | 0.71669438 | 0.723 | 0.597 | 0.04667499 |
| Gm21738 | 2.26E-06 | 0.64317718 | 0.574 | 0.455 | 0.0494238 |
| Nat6 | 2.46E-06 | -0.9733163 | 0.202 | 0.356 | 0.05359857 |
| Vat1l | 2.46E-06 | 0.88020805 | 0.539 | 0.427 | 0.05378393 |
| Tmem52 | 2.54E-06 | -1.5744588 | 0.076 | 0.203 | 0.05546009 |
| Kcnh2 | 3.10E-06 | -0.4894116 | 0.554 | 0.737 | 0.06756673 |
| Il21r | 3.38E-06 | -0.8366558 | 0.36 | 0.534 | 0.07375502 |
| Grhl2 | 3.61E-06 | 0.98604855 | 0.481 | 0.373 | 0.07868866 |
| Cd2 | 4.52E-06 | -0.8615392 | 0.204 | 0.378 | 0.09855076 |

Supplemental Table S4E: Number of Differentially sequenced ATAC chromatin segments with adjusted p values <0.1 or top hit only.

| Endothelial Cells | p_val | avg_log2FC | pct.1 | pct.2 | p_val_adj |
|---|---|---|---|---|---|
| chr17-39842818-39846951 | 5.48E-12 | 0.58190718 | 0.95 | 0.897 | 1.28E-06 |
| chr17-39847439-39848940 | 3.54E-09 | 0.62101982 | 0.765 | 0.688 | 0.00082965 |
| chr2-98666906-98667526 | 3.31E-08 | 0.29469385 | 0.974 | 0.967 | 0.00775961 |

| Neurons | p_val | avg_log2FC | pct.1 | pct.2 | p_val_adj |
|---|---|---|---|---|---|
| chr17-39842818-39846951 | 6.34E-14 | 0.51260263 | 0.989 | 0.995 | 1.49E-08 |
| chr17-39847439-39848940 | 1.97E-09 | 0.48588032 | 0.937 | 0.891 | 0.00046234 |
| chr11-109010949-109012578 | 2.37E-07 | 0.56263328 | 0.75 | 0.678 | 0.05560453 |

| CD4 T Cells | p_val | avg_log2FC | pct.1 | pct.2 | p_val_adj |
|---|---|---|---|---|---|
| chr3-128255365-128256293 | 1.39E-06 | -12.024677 | 0 | 0.115 | 0.3268215 |

| B Cells | p_val | avg_log2FC | pct.1 | pct.2 | p_val_adj |
|---|---|---|---|---|---|
| chr1-119482661-119483807 | 9.88E-07 | -11.487034 | 0 | 0.113 | 0.23163936 |

| **Ciliated Cells** | **p_val** | **avg_log2FC** | **pct.1** | **pct.2** | **p_val_adj** |
|---|---|---|---|---|---|
| chr2-98666906-98667526 | 4.34E-08 | 0.54858768 | 0.952 | 0.879 | 0.01015909 |
| chr3-138843002-138844292 | 3.96E-07 | -10.903807 | 0 | 0.106 | 0.09282599 |

| Mesothelial Cells | p_val | avg_log2FC | pct.1 | pct.2 | p_val_adj |
|---|---|---|---|---|---|
| chr5-111674930-111676918 | 2.79E-06 | -12.756341 | 0 | 0.293 | 0.65479161 |

| Macrophages | p_val | avg_log2FC | pct.1 | pct.2 | p_val_adj |
|---|---|---|---|---|---|
| chr8-46496841-46498230 | 5.62E-06 | -2.267465 | 0.07 | 0.375 | 1 |

### Supplemental Table S4E: Number of Differentially sequenced ATAC chromatin segments with adjusted p values <0.1 or top hit only.

| AT1 | p_val | avg_log2FC | pct.1 | pct.2 | p_val_adj |
|---|---|---|---|---|---|
| chr2-98666906-98667526 | 2.26E-20 | 0.96543322 | 0.845 | 0.681 | 5.31E-15 |
| chr2-98666067-98666807 | 2.29E-15 | 0.53987352 | 0.976 | 0.923 | 5.37E-10 |
| chr7-35112559-35114545 | 2.54E-10 | -2.5062086 | 0.03 | 0.187 | 5.94E-05 |
| chr5-119650936-119652216 | 3.33E-10 | -2.1182959 | 0.046 | 0.216 | 7.81E-05 |
| chr17-39842818-39846951 | 5.04E-10 | 0.77406254 | 0.902 | 0.74 | 0.00011804 |
| chr2-98662103-98663137 | 9.77E-10 | 0.50079758 | 0.894 | 0.788 | 0.00022904 |
| chr14-118000919-118002398 | 9.56E-09 | -1.8512331 | 0.054 | 0.22 | 0.00224074 |
| chr7-144597913-144598951 | 2.95E-08 | -2.9525561 | 0.014 | 0.121 | 0.00691628 |
| chr16-17795058-17797913 | 3.67E-08 | -1.4551232 | 0.098 | 0.278 | 0.00858965 |
| chr9-35304729-35306073 | 3.85E-08 | 0.79712037 | 0.579 | 0.447 | 0.00902947 |
| chr7-84605208-84607160 | 3.91E-08 | -0.7892221 | 0.315 | 0.553 | 0.00915183 |
| chr11-86960701-86963361 | 4.15E-08 | -1.11973 | 0.16 | 0.385 | 0.00972927 |
| chr1-92437443-92439004 | 4.99E-08 | -1.1244352 | 0.158 | 0.377 | 0.01169841 |
| chr4-119462406-119463945 | 5.05E-08 | -1.8688671 | 0.049 | 0.201 | 0.01183479 |
| chr3-101109495-101111093 | 6.82E-08 | -1.9172725 | 0.043 | 0.194 | 0.01599044 |
| chr19-17836678-17838545 | 8.23E-08 | -1.9721057 | 0.043 | 0.183 | 0.0192865 |
| chr7-126921886-126923365 | 1.14E-07 | -1.3815584 | 0.095 | 0.278 | 0.02670116 |
| chr4-43725714-43731824 | 1.75E-07 | -0.7993511 | 0.299 | 0.52 | 0.04106232 |
| chr3-83067422-83069137 | 1.84E-07 | -1.0906466 | 0.155 | 0.355 | 0.04319636 |
| chr1-92030144-92031247 | 2.21E-07 | -1.5804839 | 0.068 | 0.223 | 0.05179198 |
| chr17-33828289-33830049 | 2.30E-07 | -1.8775898 | 0.043 | 0.179 | 0.05386551 |
| chr4-137695090-137696707 | 2.38E-07 | -1.8088032 | 0.049 | 0.19 | 0.05576943 |
| chr2-128698063-128699549 | 2.71E-07 | -1.101312 | 0.144 | 0.374 | 0.06362335 |
| chrX-157597366-157599144 | 2.87E-07 | -1.883634 | 0.043 | 0.176 | 0.06715582 |
| chr6-125330464-125332226 | 3.34E-07 | -1.6577818 | 0.06 | 0.205 | 0.07834497 |
| chr13-48851292-48852723 | 3.91E-07 | -1.5835026 | 0.065 | 0.216 | 0.09164402 |
| chr6-55256066-55257393 | 4.13E-07 | -1.0084715 | 0.168 | 0.385 | 0.09683537 |

| AT2 | p_val | avg_log2FC | pct.1 | pct.2 | p_val_adj |
|---|---|---|---|---|---|
| chr17-39842818-39846951 | 8.07E-14 | 0.70923746 | 0.963 | 0.922 | 1.89E-08 |
| chr2-98666906-98667526 | 1.21E-13 | 0.48451339 | 0.969 | 0.968 | 2.84E-08 |
| chr17-39847439-39848940 | 4.26E-13 | 0.76239297 | 0.854 | 0.799 | 9.98E-08 |
| chr9-35304729-35306073 | 2.36E-10 | 0.56723633 | 0.854 | 0.753 | 5.52E-05 |
| chr3-5860222-5861130 | 4.97E-08 | 0.84456667 | 0.53 | 0.425 | 0.01164381 |
| chr11-109010949-109012578 | 5.86E-08 | 0.75047731 | 0.611 | 0.498 | 0.01374344 |
| chr5-146260501-146261808 | 4.01E-07 | 0.87104963 | 0.479 | 0.361 | 0.09397559 |

| Mast Cells | p_val | avg_log2FC | pct.1 | pct.2 | p_val_adj |
|---|---|---|---|---|---|
| chr14-61396648-61397641 | 1.11E-06 | -5.7195666 | 0.008 | 0.375 | 0.25928079 |

Supplemental Table S4E: Number of Differentially sequenced ATAC chromatin segments with adjusted p values <0.1 or top hit only.

| AlveolarMacs | p_val | avg_log2FC | pct.1 | pct.2 | p_val_adj |
|---|---|---|---|---|---|
| JH584304.1-58407-60889 | 3.36E-07 | -0.2989984 | 0.963 | 0.983 | 0.07871267 |
| chr7-140873926-140875278 | 3.49E-07 | -2.0317749 | 0.047 | 0.213 | 0.08182869 |
| chr12-104815037-104816057 | 4.05E-07 | -1.4184601 | 0.111 | 0.313 | 0.09502105 |

| Col1a1+ Fibroblasts | p_val | avg_log2FC | pct.1 | pct.2 | p_val_adj |
|---|---|---|---|---|---|
| chr2-98666906-98667526 | 6.97E-09 | 0.82419074 | 0.849 | 0.653 | 0.00163362 |
| chr2-98666067-98666807 | 1.52E-08 | 0.70022639 | 0.876 | 0.774 | 0.0035694 |

| AT1 | p_val | avg_log2FC | pct.1 | pct.2 | p_val_adj |
|---|---|---|---|---|---|
| chr17-39842818-39846951 | 3.37E-14 | 0.84839027 | 0.966 | 0.929 | 7.90E-09 |
| chr17-39847439-39848940 | 3.86E-12 | 0.87780018 | 0.82 | 0.797 | 9.04E-07 |
| chr17-36230622-36232240 | 1.49E-10 | 1.12543015 | 0.577 | 0.401 | 3.49E-05 |
| chr11-109010949-109012578 | 1.93E-10 | 1.10506994 | 0.573 | 0.44 | 4.52E-05 |
| chr5-146260501-146261808 | 2.66E-09 | 1.13509194 | 0.532 | 0.335 | 0.00062431 |
| chr2-98666906-98667526 | 1.81E-08 | 0.54475496 | 0.91 | 0.863 | 0.00423892 |
| chr3-5860222-5861130 | 5.92E-08 | 0.99365203 | 0.524 | 0.368 | 0.01387226 |
| chr4-151609967-151611196 | 1.02E-07 | -4.8186827 | 0.004 | 0.104 | 0.02384669 |
| chr10-24835626-24836919 | 1.99E-07 | -3.909238 | 0.007 | 0.115 | 0.04654712 |

Supplemental Table S4E: Number of Differentially sequenced ATAC chromatin segments with adjusted p values <0.1 or top hit only.

| Fibroblasts | p_val | avg_log2FC | pct.1 | pct.2 | p_val_adj |
|---|---|---|---|---|---|
| chr2-98666906-98667526 | 6.46E-15 | 0.6544296 | 0.882 | 0.723 | 1.51E-09 |
| chr17-39842818-39846951 | 9.62E-13 | 0.96049451 | 0.798 | 0.652 | 2.26E-07 |
| chr9-35304729-35306073 | 2.96E-11 | 0.72114329 | 0.685 | 0.526 | 6.93E-06 |
| chr16-18339396-18340758 | 7.18E-10 | -1.4383208 | 0.101 | 0.285 | 0.0001682 |
| chr2-166280424-166281354 | 7.43E-10 | -2.5397972 | 0.025 | 0.142 | 0.00017412 |
| chr4-130389017-130390308 | 1.05E-09 | -2.1614836 | 0.035 | 0.173 | 0.00024521 |
| chr15-84697143-84698304 | 1.72E-09 | -1.8820886 | 0.048 | 0.2 | 0.00040384 |
| chr13-60522503-60524050 | 1.81E-09 | -1.9524571 | 0.043 | 0.192 | 0.00042336 |
| chr12-113131725-113134303 | 8.88E-09 | -0.988717 | 0.204 | 0.414 | 0.00208003 |
| chr14-32826747-32829394 | 9.09E-09 | -0.925907 | 0.217 | 0.436 | 0.00213026 |
| chr9-45917025-45918322 | 9.49E-09 | -1.1114441 | 0.154 | 0.348 | 0.00222321 |
| chr17-3463299-3464769 | 1.38E-08 | -2.6217386 | 0.018 | 0.123 | 0.00323765 |
| chr1-36570556-36573310 | 1.54E-08 | -0.9353775 | 0.209 | 0.433 | 0.00360979 |
| chr9-119197140-119198278 | 1.56E-08 | -1.8950709 | 0.045 | 0.173 | 0.00365622 |
| chr4-128733165-128734825 | 1.56E-08 | -0.8805968 | 0.237 | 0.46 | 0.00366232 |
| chr1-92672567-92674446 | 1.79E-08 | -1.1828997 | 0.131 | 0.312 | 0.0042056 |
| chr10-127599857-127601158 | 2.08E-08 | -1.1040331 | 0.151 | 0.34 | 0.00487844 |
| chr16-97751024-97752673 | 2.19E-08 | -1.2722634 | 0.108 | 0.288 | 0.0051389 |
| chr8-68167800-68170293 | 2.34E-08 | -1.8830046 | 0.045 | 0.17 | 0.00548507 |
| chr18-14330710-14332423 | 2.35E-08 | -0.6106242 | 0.403 | 0.633 | 0.00551828 |
| chr13-38878394-38879702 | 2.68E-08 | -2.0670119 | 0.033 | 0.153 | 0.00628408 |
| chr2-27554068-27555069 | 3.16E-08 | -2.90454 | 0.013 | 0.104 | 0.00740638 |
| chr12-16869042-16869871 | 3.25E-08 | -1.9744577 | 0.035 | 0.162 | 0.00762315 |
| chr7-132447956-132449346 | 3.26E-08 | -1.7764666 | 0.048 | 0.184 | 0.00764832 |
| chr12-104406096-104407842 | 3.46E-08 | -1.6508124 | 0.058 | 0.2 | 0.00809702 |
| chr5-52523792-52525289 | 3.76E-08 | -1.6225951 | 0.058 | 0.205 | 0.0088012 |
| chr11-119430796-119432353 | 4.49E-08 | -1.5097686 | 0.073 | 0.219 | 0.01052154 |
| chr4-142756055-142757352 | 5.71E-08 | -1.3241422 | 0.093 | 0.26 | 0.01338909 |
| chr6-140999670-141001649 | 5.79E-08 | -1.5015459 | 0.076 | 0.216 | 0.01356334 |
| chr6-143367185-143368449 | 6.33E-08 | -2.1082531 | 0.03 | 0.142 | 0.01484058 |
| chr16-43945316-43946432 | 7.18E-08 | -1.6994142 | 0.053 | 0.184 | 0.0168255 |
| chr2-167843829-167845408 | 8.70E-08 | -1.1064008 | 0.136 | 0.315 | 0.02038655 |
| chr16-4652399-4653728 | 1.31E-07 | -1.6486865 | 0.055 | 0.186 | 0.03071163 |
| chr18-75502291-75504436 | 1.34E-07 | -0.7477088 | 0.29 | 0.501 | 0.03142737 |
| chr1-93209735-93211429 | 1.35E-07 | -1.2014265 | 0.108 | 0.288 | 0.03170083 |
| chr12-113141948-113143681 | 1.39E-07 | -1.0014795 | 0.171 | 0.353 | 0.03246702 |
| chr11-47705004-47706220 | 1.54E-07 | -1.7350843 | 0.048 | 0.173 | 0.03605931 |
| chr6-136906020-136907874 | 1.63E-07 | -1.137594 | 0.128 | 0.29 | 0.03813241 |
| chr19-24307293-24308784 | 1.85E-07 | -1.945982 | 0.035 | 0.151 | 0.0433634 |
| chr5-65456482-65459036 | 2.07E-07 | -1.144343 | 0.121 | 0.288 | 0.04841435 |
| chr6-49095822-49097613 | 2.19E-07 | -0.6784696 | 0.312 | 0.542 | 0.05133466 |
| chr17-45583460-45586366 | 2.30E-07 | -1.0635095 | 0.149 | 0.321 | 0.05378701 |
| chr4-117846805-117847842 | 2.52E-07 | -1.7491943 | 0.043 | 0.167 | 0.05909187 |
| chr6-111584373-111585533 | 2.52E-07 | -0.8674691 | 0.212 | 0.389 | 0.05916818 |
| chr17-39847439-39848940 | 2.61E-07 | 0.89125903 | 0.494 | 0.389 | 0.06124624 |
| chr11-11856425-11857560 | 2.63E-07 | -1.2572616 | 0.098 | 0.249 | 0.0615866 |
| chr4-111624212-111625660 | 2.76E-07 | -1.5419556 | 0.06 | 0.192 | 0.06463287 |
| chr1-177014257-177015266 | 2.89E-07 | -1.7883126 | 0.043 | 0.159 | 0.06775598 |
| chr7-117493916-117495761 | 3.13E-07 | -0.9985542 | 0.159 | 0.334 | 0.07345963 |
| chr14-79034135-79035904 | 3.39E-07 | -1.9332915 | 0.035 | 0.142 | 0.0793698 |
| chr14-24747413-24748971 | 3.70E-07 | -1.1649663 | 0.116 | 0.271 | 0.08664045 |
| chr8-12788226-12789435 | 3.72E-07 | -1.4125725 | 0.073 | 0.211 | 0.08728403 |
| chr12-38136093-38137358 | 3.93E-07 | -1.0957756 | 0.134 | 0.285 | 0.09216294 |

Supplemental Table S4F: IL-1b and IL-18 ATAC seq differences between FR and FF lungs (all cell types combined, pseudobulk analysis)

| | p_val | avg_log2FC | pct.1 | pct.2 | p_val_adj |
|---|---|---|---|---|---|
| Il1b | 0.21248498 | 0.38126016 | 0.102 | 0.095 | 1 |
| Il18 | 0.19556494 | 0.67364294 | 0.057 | 0.05 | 1 |

| | p_val | avg_log2FC | pct.1 | pct.2 | p_val_adj |
|---|---|---|---|---|---|
| **il1b chr2-129,206,490–129,213,059** | | | | | |
| chr2-129206405-129207502 | 0.00010375 | -0.5754807 | 0.028 | 0.047 | 1 |
| chr2-129209899-129210819 | 0.0100254 | -0.4468081 | 0.023 | 0.034 | 1 |
| chr2-129207932-129208895 | 0.07132926 | -0.3216594 | 0.018 | 0.025 | 1 |
| chr2-129212226-129213333 | 0.01171048 | -0.3707286 | 0.024 | 0.035 | 1 |
| **il18 chr9-50,466,127 and 50,493,140** | | | | | |
| chr9-50467079-50468445 | 0.03935128 | -0.3111103 | 0.029 | 0.039 | 1 |
| chr9-50470068-50471506 | 0.01072068 | -0.2808366 | 0.043 | 0.058 | 1 |
| chr9-50478300-50480249 | 1.99E-08 | -0.59697 | 0.069 | 0.109 | 0.00466045 |
| chr9-50482277-50483455 | 0.00016391 | -0.536613 | 0.032 | 0.051 | 1 |
| chr9-50490313-50492497 | 3.35E-05 | -0.4877965 | 0.041 | 0.065 | 1 |
